# CytoGate-Bench: an LLM benchmark for cross-panel cell gating in cytometry

**DOI:** 10.64898/2026.08.24.746336

**Authors:** Jaesik Kim, Byounghan Lee, Namhyuk Ahn, Matei Ionita, Michelle L. McKeague, Matthew E. Lee, Chang-Uk Jeong, Sokratis A. Apostolidis, Amy E. Baxter, Shwetank, Allison R. Greenplate, E. John Wherry, Kyung-Ah Sohn, Dokyoon Kim

**Author notes:** These authors contributed equally (Jaesik Kim, Byounghan Lee).

## Abstract

In cytometry, the workhorse single-cell technology of clinical immunology, every study defines its own antibody panel and cell-type vocabulary, so a classifier trained on one cannot annotate the next. Immunologists instead annotate by manual gating, splitting one parent population at a time on a two-marker plot, down an expert-defined hierarchy. We introduce CytoGate-Bench, a benchmark that reformulates this per-step procedure as a zero-shot, panel-agnostic task for large language models. It comprises 23,646 expert-annotated instances re-curated from 11 public flow- and mass-cytometry cohorts spanning eight marker panels. Across six open- and closed-weight backbones, the strongest formulation draws one rectangular gate per candidate and falls within the range of trained, panel-specialized baselines. It degrades less under distribution shift. Walking the hierarchy stepwise outperforms predicting every cell type at once. Ablations trace the signal to the data distribution shape and curated marker priors. However, adding vision or a self-verification loop systematically tightens gates.

**THE BIGGER PICTURE:** Immunology laboratories worldwide profile blood and tissue with cytometry, an instrument family that measures dozens of protein markers on millions of individual cells. Before any biology can be read out, every cell must be assigned an identity, a step still dominated by manual “gating,” in which an expert draws boundaries on a sequence of two-marker plots, following a documented, hierarchical protocol. Automating this step has remained difficult because every study measures a different marker panel and names a different set of cell types, so conventional machine-learning models must be retrained for each new study.

Large language models (LLMs) promise a different route, a single general-purpose model that reads the expert’s protocol and the data and makes each gating decision directly, with no study-specific training. This work contributes a public benchmark that tests precisely that ability across 11 human cohorts. The result is a statement of feasibility rather than superiority. Off-the-shelf models already score in the range of study-specific trained models and tolerate the day-to-day variation that degrades them. That capability matters most for new or small studies, for which no labeled training data exists. Walking the expert’s hierarchy one decision at a time also outperforms asking the model to name every cell type in a single pass, evidence that the structure of expert practice matters more than the scale of the question. These results come from a deliberately minimal setup, untuned models drawing simple rectangular gates, so we read them as a floor rather than a ceiling. Cytometry-aware training, richer gate geometries, and better-calibrated visual feedback are open avenues, and the benchmark gives that progress a fixed yardstick. Sustained progress would give laboratories analysts that keep pace with evolving marker panels without retraining, while leaving a decision trail an immunologist can audit.

## INTRODUCTION

The *AI co-scientist* paradigm increasingly automates scientific workflows with Large Language Model (LLM) agents^1^. Whereas conventional automation fixes one predictive model per task, a single general-purpose model adapts to each analysis through instructions and context. The shift is already underway in single-cell biology, where agents^2,3^ plan and run transcriptomic analyses. Computational cytometry, however, has neither an LLM-based method nor a standardized benchmark, and LLM agents have not yet been applied to *manual gating*, the domain’s definitive cell-type annotation and the entry point for downstream immunological analyses. Automating it is thus a clear first milestone.

The core obstacle is that cytometry studies do not share a common feature (marker) space. Each study defines its own marker panel and its own cell-type label set, with new markers added as panels are extended (Figure 1A, top). A classifier trained on one study’s panel is bound to that fixed input and label vocabulary (*flat annotation*), so it cannot transfer to a study whose panel differs, and each new study needs its own model retrained from scratch. Annotating cells in a way that transfers across panels remains the open bottleneck.

**Figure 1.**
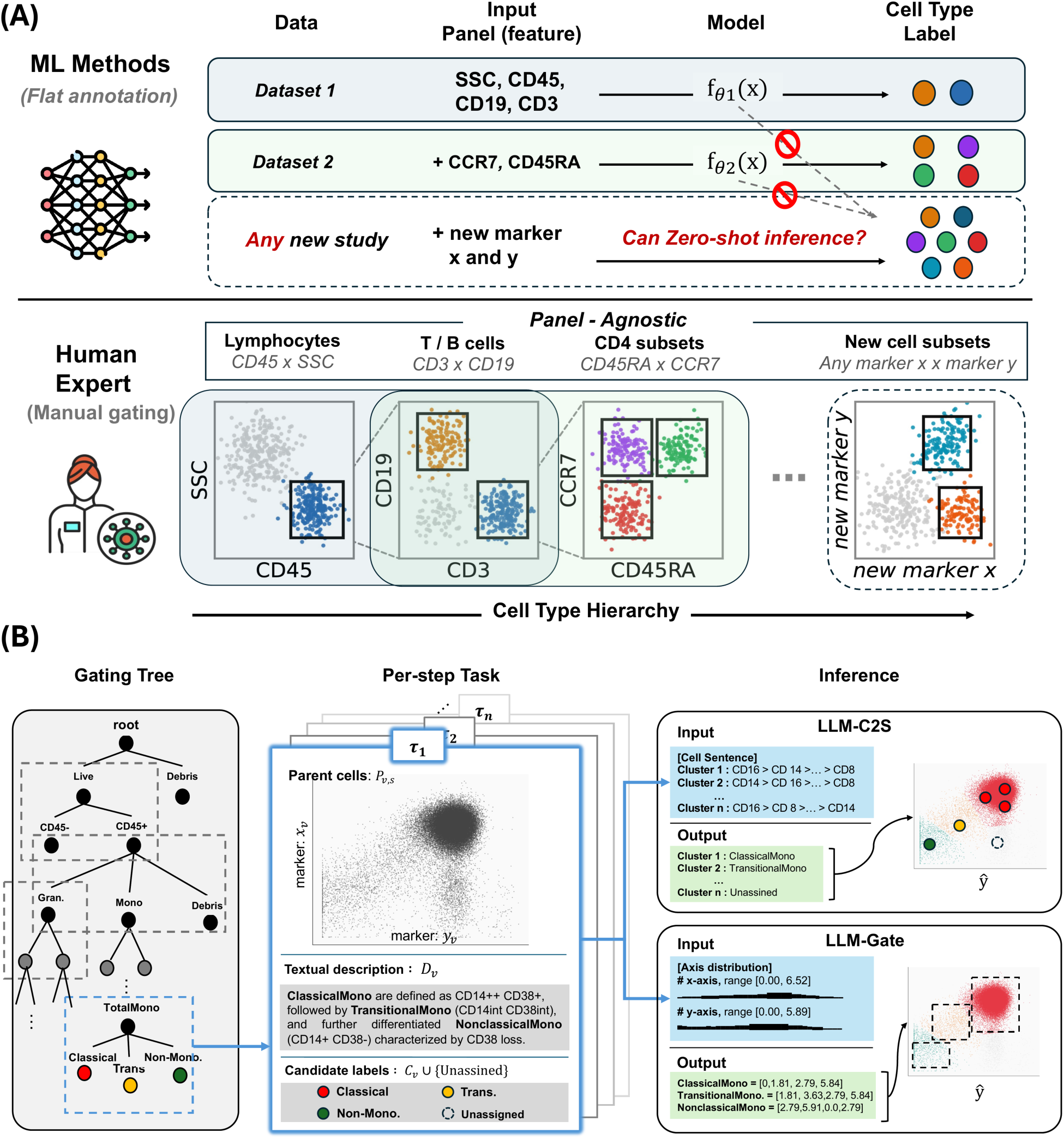
Per-step gating transfers across marker panels and defines the annotation task. (A) Panel heterogeneity and the two annotation regimes. Top: per-study flat annotation binds a model to one study’s marker panel and label set, so it does not transfer across panels. Bottom: per-step manual gating makes one local decision per node of an expert gating tree and transfers across panels by construction. (B) The expert gating tree fixes, at each internal node, the parent population, one marker pair, the candidate children, and curated marker priors; the per-step task is to assign every cell of the parent to one candidate (or *Unassigned*). Inputs (*P_v_*_,*s*_, *x_v_*, *y_v_*, *C_v_*, *D_v_*) vary across nodes; methods are evaluated zero-shot per node. See also supplemental sections S1, S3, and S4.

Expert practice already avoids this limitation. Cell types form a *strict taxonomic hierarchy*, e.g., CD4^+^ T cell → memory CD4^+^ T cell, in which each type is defined by its parent population and by the marker pair that separates it from its siblings. Manual gating turns this definition into a procedure, in which an immunologist walks the hierarchy top-down, reading one biaxial marker-pair scatter per node to split the current population into subtypes, such as CD45 × side scatter (SSC) to gate lymphocytes and then CD3 × CD19 for T versus B cells (Figure 1A, bottom). We formulate each local decision as a *step*, and chaining steps from root to leaf yields a cell’s full hierarchical label. Because the hierarchy branches along parallel analytical axes, a single cell can follow more than one branch and therefore carries several labels at once. A CD4^+^ T cell may be *memory* under CCR7 × CD45RA and *activated* under HLA-DR × CD38, both correct. Because a step operates on a single marker pair and the shape of the data rather than a fixed global feature vector, per-step gating is panel-agnostic by construction, even handling marker pairs not seen before.

This makes LLMs a natural fit. An LLM can read a per-step description together with the empirical distribution on a biaxial plot, ground them in biomedical priors, and emulate the expert’s local decision, generalizing zero-shot across cohorts. Two questions follow. First, can off-the-shelf LLMs match domain-specific supervised models zero-shot, on panels they were never trained on? Second, if so, does that success come from recalling memorized marker-name associations, or from reading the shape of the data itself, as an immunologist would?

Cytometry annotation has been studied predominantly as flat classification. Supervised classifiers^4–6^ and recent foundation models^7,8^ bind to a fixed marker space defined at training time and are brittle under panel shifts. Unsupervised clustering^9,10^ sidesteps this but still needs labor-intensive per-cohort labeling. Established benchmarks^11,12^ score flat readouts on single panels. A recent evaluation of 23 clustering and four auto-gating tools across six cohorts ends in a practical recommendation flowchart, keyed to whether the analyst has prior marker knowledge or labeled data from the target study^13^. Every branch of that flowchart terminates in one of two outcomes. Unsupervised clustering returns partitions without labels, and auto-gating requires a marker table or labeled cells from the very study being analyzed. Labeled annotation without study-specific input appears on no branch.

Per-step methods preserve the hierarchical structure of gating but remain panel-bound. UNITO^14^ and flowMagic^15^ fit model weights to specific training panels, whereas flowDensity^16^ runs fully unsupervised but has no notion of marker semantics and still requires a gating strategy hand-specified per cohort. To our knowledge, no method performs panel-agnostic, zero-shot per-step gating, assigning every cell to the correct child of a *given* expert gating tree without panel-specific training. We expand this comparison and motivate per-step as the natural annotation primitive in the supplemental information (section S1).

Prior LLM work on single-cell data has centered on single-cell RNA sequencing (scRNA-seq). One line serializes molecular profiles into text and prompts a model for a label, rendering each cell as a “cell sentence” of top-expressed genes (Cell2Sentence^17^, C2S-Scale^18^, CellVerse^19^) or labeling whole clusters from their top marker genes (GPTCelltype^20^, Cell-o1^21^). A second line casts the LLM as an agent that plans bioinformatics workflows and invokes annotation as one routine^2,3,22–27^. In both, the input is a text-serialized list of gene names.

In short, prior methods either fix the marker vocabulary at training time or, in the scRNA-seq LLM setting, read text-serialized gene names to emit a single per-cell or per-cluster label. CytoGate-Bench departs on both axes, framing annotation as zero-shot, panel-agnostic per-step gating that reads the empirical marker distribution under a given expert gating tree. No prior method addresses this regime, including our closest relative Cell-o1.

To answer the two questions above, we introduce **CytoGate-Bench**, a benchmark that frames hierarchical manual gating as an LLM task. It re-curates 11 public cohorts spanning eight panels into 23,646 per-step instances. We evaluate two formulations, *LLM-C2S* (clusters the parent population and labels each cluster from its cell sentence) and *LLM-Gate* (draws one axis-aligned rectangle per candidate). Both run zero-shot across six open- and closed-weight backbones and a domain-tuned reference (Cell- o1^21^). We further augment *LLM-Gate* with two variants that address the backbone as a vision-language model (VLM). *VLM-Gate* adds the scatter plot as an image, and *Agent- Gate* adds a tool loop that renders and refines its predicted boundaries. Across all 11 cohorts, the best LLM-Gate configuration scores within the range spanned by trained, panel-specialized baselines without any panel-specific training, and degrades less when populations are depleted or channels drift. Prompt ablations locate that signal primarily in the shape of the parent distribution, with curated marker priors contributing, and show that adding an image or a self-verification loop makes the model tighten its gates rather than place them better.

## RESULTS

### CytoGate-Bench reformulates expert gating as a panel-agnostic, per-step LLM task

We re-curate 11 public flow-cytometry and mass-cytometry (CyTOF) cohorts (∼835 samples) into a harmonized per-step format in which every gating-tree node is paired with its marker pair, candidate cell types, curated marker priors, and expert ground truth, yielding 23,646 evaluation instances (Table 1, supplemental information, section S2.1). The cohorts vary in panel, donor, tissue, and lineage target, and this variation makes the benchmark a demanding test of cross-panel generalization.

**Table 1.**
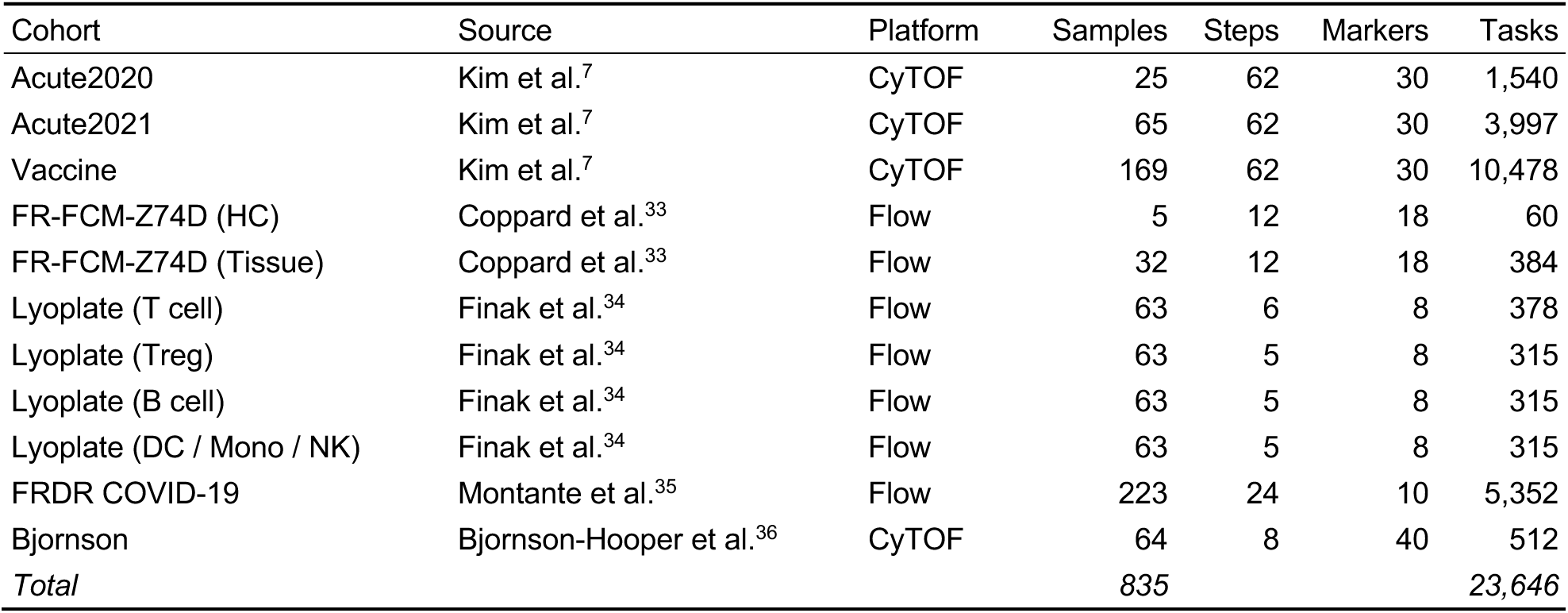
The 11 curated cohorts. *Source* names the study each cohort is re-curated from; *Tasks* = per-step instances (samples × steps, excluding steps whose parent population is empty in a given sample). CyTOF, mass cytometry; HC, healthy control; DC, dendritic cell; Mono, monocyte; NK, natural killer cell.

| Cohort | Source | Platform | Samples | Steps | Markers | Tasks |
| --- | --- | --- | --- | --- | --- | --- |
| Acute2020 | Kim et al.<sup>7</sup> | CyTOF | 25 | 62 | 30 | 1,540 |
| Acute2021 | Kim et al.<sup>7</sup> | CyTOF | 65 | 62 | 30 | 3,997 |
| Vaccine | Kim et al.<sup>7</sup> | CyTOF | 169 | 62 | 30 | 10,478 |
| FR-FCM-Z74D (HC) | Coppard et al.<sup>33</sup> | Flow | 5 | 12 | 18 | 60 |
| FR-FCM-Z74D (Tissue) | Coppard et al.<sup>33</sup> | Flow | 32 | 12 | 18 | 384 |
| Lyoplate (T cell) | Finak et al.<sup>34</sup> | Flow | 63 | 6 | 8 | 378 |
| Lyoplate (Treg) | Finak et al.<sup>34</sup> | Flow | 63 | 5 | 8 | 315 |
| Lyoplate (B cell) | Finak et al.<sup>34</sup> | Flow | 63 | 5 | 8 | 315 |
| Lyoplate (DC / Mono / NK) | Finak et al.<sup>34</sup> | Flow | 63 | 5 | 8 | 315 |
| FRDR COVID-19 | Montante et al.<sup>35</sup> | Flow | 223 | 24 | 10 | 5,352 |
| Bjornson | Bjornson-Hooper et al.<sup>36</sup> | CyTOF | 64 | 8 | 40 | 512 |
| *Total* |  |  | *835* |  |  | *23,646* |

Figure 1B (left, center) illustrates the unit of evaluation in CytoGate-Bench. The expert-specified *gating tree* organizes the cell-type hierarchy as a rooted tree. The root represents all cells in a sample, and each internal node represents a population that the expert further splits into a small set of children. We define a **per-step task** as the local decision at one such node, assigning each cell reaching the node to one of its candidate children. A benchmark instance is one per-step task on one sample, and chaining instances from root to leaf reconstructs the full hierarchical annotation, with per-step error compounding only linearly in tree depth (supplemental section S3), so per-step accuracy is our headline metric.

A single step spans 10^2^–10^5^ cells, far beyond any LLM context window, so a method is defined entirely by how it reduces the parent population *P_v_*_,*s*_ to something the model can read. We evaluate two such encodings, holding the rest of the task fixed. *LLM-C2S* follows the group-then-annotate scheme common in single-cell analysis, clustering the parent population and labeling one cluster at a time from its *cell sentence*^17^, after Cell- o1^21^. *LLM-Gate* skips clustering and prompts the model for one axis-aligned rectangle per candidate from per-axis histograms given as text rather than as a rendered image^28^, mirroring how an immunologist places gates on a biaxial plot (Figure 1B). Implementations, prompts, and propagation rules are in the methods and in supplemental sections S4 and S5.

### Zero-shot LLM gating scores in the range of trained, panel-specialized baselines

We compare three method groups on a shared held-out test split. The *trained baselines*, UNITO^14^ and cyMAE^7^, are fit per cohort. The LLM-class methods, *LLM-C2S* and *LLM-Gate*, run zero-shot. We report **F1 macro**, **Balanced accuracy** (BA), and **Hull intersection-over-union** (Hull IoU), averaged within each cohort and then across the 11 cohorts with equal weight. Cohort split, method roster and backbones, generation settings, and metric definitions are in the methods and supplemental section S6.

Among the LLM methods, LLM-Gate is the strongest formulation (Table 2). It holds the best LLM-class value on F1 macro (GPT-5.4, 0.676) and on Hull IoU (Gemma 4 31B, 0.614), ahead of the clustering-based LLM-C2S rows.

**Table 2.** Zero-shot per-step gating evaluation. Per-cohort F1 macro / balanced accuracy (BA) / Hull IoU; *Mean* = equal-weight mean over the 11 cohorts. Bold = best per column; underline = best LLM-class value per column. ^†^ = closed-weight backbone. Cohort abbreviations: A2020/A2021 = Acute2020/2021, Vacc = Vaccine, Bj = Bjornson, Z74D-H/T = FR-FCM-Z74D HC/Tissue, FRDR = FRDR COVID-19, LP- T/Tr/B/D = Lyoplate T cell/Treg/B cell/DC-Mono-NK.

| Cohort | A2020 |  |  | A2021 |  |  | Vacc |  |  | Bj |  |  | Z74D-H |  |  | Z74D-T |  |  |
| --- | --- | --- | --- | --- | --- | --- | --- | --- | --- | --- | --- | --- | --- | --- | --- | --- | --- | --- |
| Metric | F1 | BA | IoU | F1 | BA | IoU | F1 | BA | IoU | F1 | BA | IoU | F1 | BA | IoU | F1 | BA | IoU |
| *Trained baselines* |  |  |  |  |  |  |  |  |  |  |  |  |  |  |  |  |  |  |
| UNITO (2D pair)<sup>a</sup> | **0.677** | 0.760 | 0.622 | **0.753** | **0.828** | **0.755** | **0.808** | **0.846** | **0.832** | **0.929** | **0.928** | **0.913** | 0.559 | 0.629 | 0.647 | 0.446 | 0.505 | 0.473 |
| cyMAE (all-marker)<sup>a</sup> | 0.451 | 0.561 | 0.379 | 0.555 | 0.691 | 0.501 | 0.612 | 0.747 | 0.568 | 0.811 | 0.864 | 0.691 | 0.542 | 0.612 | 0.497 | 0.448 | 0.562 | 0.464 |
| *LLM-C2S* |  |  |  |  |  |  |  |  |  |  |  |  |  |  |  |  |  |  |
| *density-grid (flowDensity)* |  |  |  |  |  |  |  |  |  |  |  |  |  |  |  |  |  |  |
| Cell-o1 (7B) | 0.340 | 0.416 | 0.280 | 0.336 | 0.419 | 0.285 | 0.338 | 0.426 | 0.288 | 0.424 | 0.487 | 0.376 | 0.338 | 0.395 | 0.339 | 0.316 | 0.388 | 0.303 |
| Qwen3.5-4B | 0.515 | 0.610 | 0.454 | 0.534 | 0.646 | 0.477 | 0.542 | 0.648 | 0.482 | 0.751 | 0.764 | 0.743 | 0.591 | 0.633 | 0.634 | 0.517 | 0.618 | 0.550 |
| Qwen3.5-27B | 0.592 | 0.700 | 0.515 | 0.601 | 0.721 | 0.533 | 0.584 | 0.692 | 0.517 | 0.807 | 0.821 | 0.783 | 0.619 | 0.637 | 0.672 | 0.545 | 0.634 | 0.584 |
| Qwen3.6-27B | 0.582 | 0.691 | 0.508 | 0.592 | 0.710 | 0.527 | 0.577 | 0.683 | 0.513 | 0.780 | 0.793 | 0.772 | 0.621 | 0.654 | 0.669 | 0.542 | 0.633 | 0.575 |
| Gemma 4 26B A4B | 0.576 | 0.680 | 0.505 | 0.586 | 0.709 | 0.518 | 0.570 | 0.679 | 0.501 | 0.741 | 0.758 | 0.708 | 0.622 | 0.655 | 0.671 | 0.534 | 0.633 | 0.569 |
| Gemma 4 31B | 0.587 | 0.692 | 0.513 | 0.599 | 0.712 | 0.537 | 0.585 | 0.682 | 0.525 | 0.798 | 0.805 | 0.783 | 0.628 | 0.659 | 0.678 | 0.544 | 0.631 | 0.584 |
| GPT-5.4<sup>†</sup> | 0.568 | 0.674 | 0.498 | 0.583 | 0.701 | 0.524 | 0.568 | 0.674 | 0.515 | 0.795 | 0.802 | 0.780 | 0.630 | 0.647 | 0.688 | 0.536 | 0.633 | 0.581 |
| *clustering (FlowSOM)* |  |  |  |  |  |  |  |  |  |  |  |  |  |  |  |  |  |  |
| Cell-o1 (7B) | 0.319 | 0.385 | 0.254 | 0.317 | 0.396 | 0.249 | 0.316 | 0.394 | 0.260 | 0.375 | 0.460 | 0.314 | 0.243 | 0.323 | 0.217 | 0.273 | 0.341 | 0.277 |
| Qwen3.5-4B | 0.633 | 0.736 | 0.603 | 0.629 | 0.742 | 0.603 | 0.635 | 0.741 | 0.609 | 0.682 | 0.703 | 0.605 | 0.536 | 0.598 | 0.591 | 0.447 | 0.559 | 0.480 |
| Qwen3.5-27B | 0.636 | 0.734 | 0.603 | 0.640 | 0.758 | 0.613 | 0.660 | 0.766 | 0.635 | 0.794 | 0.809 | 0.711 | 0.588 | 0.632 | 0.630 | 0.542 | **0.650** | 0.597 |
| Qwen3.6-27B | 0.569 | 0.659 | 0.546 | 0.606 | 0.718 | 0.595 | 0.632 | 0.739 | 0.618 | 0.737 | 0.758 | 0.671 | 0.567 | 0.628 | 0.617 | 0.500 | 0.617 | 0.521 |
| Gemma 4 26B A4B | 0.652 | 0.754 | 0.629 | 0.645 | 0.764 | 0.634 | 0.655 | 0.762 | 0.634 | 0.727 | 0.741 | 0.643 | 0.571 | 0.630 | 0.629 | 0.505 | 0.621 | 0.527 |
| Gemma 4 31B | 0.659 | 0.762 | **0.634** | 0.656 | 0.772 | **0.648** | 0.670 | 0.774 | **0.659** | 0.766 | 0.768 | 0.696 | 0.577 | 0.623 | 0.628 | 0.522 | 0.630 | 0.548 |
| GPT-5.4<sup>†</sup> | 0.615 | 0.734 | 0.582 | 0.612 | 0.743 | 0.589 | 0.610 | 0.741 | 0.594 | 0.684 | 0.710 | 0.589 | 0.575 | 0.612 | 0.604 | 0.496 | 0.608 | 0.514 |
| *LLM-Gate* |  |  |  |  |  |  |  |  |  |  |  |  |  |  |  |  |  |  |
| Cell-o1 (7B) | 0.411 | 0.516 | 0.260 | 0.421 | 0.531 | 0.274 | 0.412 | 0.510 | 0.268 | 0.317 | 0.444 | 0.265 | 0.340 | 0.445 | 0.307 | 0.234 | 0.353 | 0.189 |
| Qwen3.5-4B | 0.643 | 0.744 | 0.567 | 0.638 | 0.756 | 0.566 | 0.653 | 0.754 | 0.590 | 0.757 | 0.782 | 0.707 | 0.597 | 0.642 | 0.621 | 0.528 | 0.615 | 0.556 |
| Qwen3.5-27B | **0.667** | 0.764 | 0.608 | **0.658** | 0.772 | 0.602 | **0.676** | 0.773 | 0.633 | 0.816 | 0.815 | **0.785** | 0.633 | 0.668 | 0.683 | **0.557** | 0.639 | **0.613** |
| Qwen3.6-27B | 0.661 | 0.762 | 0.605 | 0.650 | 0.768 | 0.596 | 0.672 | 0.773 | 0.636 | 0.813 | 0.814 | 0.782 | 0.620 | 0.663 | 0.664 | 0.550 | 0.639 | 0.597 |
| Gemma 4 26B A4B | 0.627 | 0.739 | 0.555 | 0.619 | 0.742 | 0.540 | 0.636 | 0.745 | 0.585 | 0.801 | 0.803 | 0.767 | 0.625 | 0.672 | 0.662 | 0.534 | 0.631 | 0.578 |
| Gemma 4 31B | 0.659 | **0.767** | 0.606 | 0.648 | 0.767 | 0.599 | **0.676** | 0.780 | 0.644 | 0.811 | 0.809 | 0.782 | 0.643 | 0.684 | **0.692** | 0.540 | 0.629 | 0.594 |
| GPT-5.4<sup>†</sup> | 0.660 | **0.767** | 0.570 | 0.644 | **0.774** | 0.562 | 0.671 | **0.783** | 0.614 | **0.821** | **0.848** | 0.774 | **0.650** | **0.705** | 0.678 | 0.554 | 0.639 | 0.564 |

| Cohort | FRDR |  |  | LP-T |  |  | LP-Tr |  |  | LP-B |  |  | LP-D |  |  | Mean |  |  |
| --- | --- | --- | --- | --- | --- | --- | --- | --- | --- | --- | --- | --- | --- | --- | --- | --- | --- | --- |
| Metric | F1 | BA | IoU | F1 | BA | IoU | F1 | BA | IoU | F1 | BA | IoU | F1 | BA | IoU | F1 | BA | IoU |
| *Trained baselines* |  |  |  |  |  |  |  |  |  |  |  |  |  |  |  |  |  |  |
| UNITO (2D pair)<sup>a</sup> | 0.586 | 0.709 | 0.562 | **0.739** | **0.873** | **0.798** | 0.639 | 0.646 | 0.477 | 0.715 | 0.808 | **0.598** | 0.741 | 0.789 | 0.640 | **0.690** | 0.757 | **0.665** |
| cyMAE (all-marker)<sup>a</sup> | **0.764** | **0.882** | **0.767** | 0.687 | 0.870 | 0.702 | **0.828** | **0.919** | **0.798** | **0.741** | **0.881** | 0.562 | **0.778** | **0.911** | 0.646 | 0.656 | **0.773** | 0.598 |
| *LLM-C2S* |  |  |  |  |  |  |  |  |  |  |  |  |  |  |  |  |  |  |
| *density-grid (flowDensity)* |  |  |  |  |  |  |  |  |  |  |  |  |  |  |  |  |  |  |
| Cell-o1 (7B) | 0.444 | 0.538 | 0.413 | 0.260 | 0.323 | 0.206 | 0.464 | 0.530 | 0.313 | 0.420 | 0.557 | 0.267 | 0.427 | 0.495 | 0.307 | 0.373 | 0.452 | 0.307 |
| Qwen3.5-4B | 0.632 | 0.737 | 0.630 | 0.614 | 0.763 | 0.624 | 0.638 | 0.697 | 0.501 | 0.604 | 0.740 | 0.433 | 0.642 | 0.774 | 0.585 | 0.598 | 0.694 | 0.556 |
| Qwen3.5-27B | 0.657 | 0.760 | 0.648 | 0.635 | 0.779 | 0.657 | 0.653 | 0.716 | 0.536 | 0.652 | 0.781 | 0.480 | **0.744** | **0.865** | **0.657** | 0.644 | 0.737 | 0.598 |
| Qwen3.6-27B | 0.660 | 0.752 | 0.665 | 0.622 | 0.780 | 0.635 | 0.649 | 0.706 | 0.521 | 0.652 | 0.773 | 0.484 | 0.712 | 0.829 | 0.634 | 0.635 | 0.728 | 0.591 |
| Gemma 4 26B A4B | 0.657 | 0.750 | 0.657 | 0.630 | 0.775 | 0.645 | 0.651 | 0.708 | 0.521 | 0.612 | 0.748 | 0.440 | 0.692 | 0.820 | 0.618 | 0.625 | 0.720 | 0.578 |
| Gemma 4 31B | 0.663 | 0.754 | 0.672 | 0.627 | 0.764 | 0.652 | 0.651 | 0.712 | 0.524 | 0.654 | 0.780 | 0.481 | 0.740 | 0.862 | 0.656 | 0.643 | 0.732 | 0.600 |
| GPT-5.4<sup>†</sup> | 0.651 | 0.752 | 0.643 | 0.620 | 0.779 | 0.630 | 0.645 | 0.700 | 0.490 | 0.616 | 0.756 | 0.425 | 0.741 | 0.839 | 0.625 | 0.632 | 0.723 | 0.582 |
| *clustering (FlowSOM)* |  |  |  |  |  |  |  |  |  |  |  |  |  |  |  |  |  |  |
| Cell-o1 (7B) | 0.414 | 0.525 | 0.388 | 0.245 | 0.314 | 0.160 | 0.379 | 0.421 | 0.185 | 0.370 | 0.479 | 0.203 | 0.317 | 0.387 | 0.180 | 0.324 | 0.402 | 0.244 |
| Qwen3.5-4B | 0.644 | 0.764 | 0.638 | 0.354 | 0.508 | 0.254 | 0.513 | 0.568 | 0.233 | 0.509 | 0.675 | 0.333 | 0.448 | 0.547 | 0.302 | 0.548 | 0.649 | 0.477 |
| Qwen3.5-27B | 0.694 | **0.811** | 0.683 | 0.546 | 0.675 | 0.479 | 0.590 | 0.657 | 0.407 | 0.659 | 0.802 | 0.498 | 0.613 | 0.702 | 0.459 | 0.633 | 0.727 | 0.574 |
| Qwen3.6-27B | 0.672 | 0.788 | 0.681 | 0.373 | 0.548 | 0.277 | 0.515 | 0.577 | 0.268 | 0.581 | 0.735 | 0.371 | 0.529 | 0.610 | 0.359 | 0.571 | 0.671 | 0.502 |
| Gemma 4 26B A4B | 0.693 | 0.797 | 0.694 | 0.495 | 0.622 | 0.422 | 0.571 | 0.635 | 0.352 | 0.591 | 0.743 | 0.400 | 0.546 | 0.644 | 0.380 | 0.605 | 0.701 | 0.540 |
| Gemma 4 31B | 0.705 | 0.808 | **0.719** | 0.535 | 0.648 | 0.466 | 0.573 | 0.639 | 0.363 | 0.643 | 0.786 | 0.460 | 0.636 | 0.727 | 0.467 | 0.631 | 0.722 | 0.572 |
| GPT-5.4<sup>†</sup> | 0.585 | 0.704 | 0.594 | 0.333 | 0.514 | 0.222 | 0.542 | 0.598 | 0.317 | 0.590 | 0.741 | 0.374 | 0.528 | 0.605 | 0.316 | 0.561 | 0.665 | 0.481 |
| *LLM-Gate* |  |  |  |  |  |  |  |  |  |  |  |  |  |  |  |  |  |  |
| Cell-o1 (7B) | 0.457 | 0.608 | 0.381 | 0.201 | 0.286 | 0.171 | 0.353 | 0.411 | 0.143 | 0.345 | 0.483 | 0.189 | 0.310 | 0.403 | 0.154 | 0.346 | 0.454 | 0.236 |
| Qwen3.5-4B | 0.637 | 0.756 | 0.554 | 0.540 | 0.675 | 0.546 | 0.641 | 0.693 | 0.514 | 0.636 | 0.737 | 0.505 | 0.592 | 0.658 | 0.442 | 0.624 | 0.710 | 0.561 |
| Qwen3.5-27B | 0.681 | 0.778 | 0.636 | 0.627 | 0.759 | 0.636 | 0.619 | 0.667 | 0.496 | 0.658 | 0.771 | 0.546 | 0.618 | 0.709 | 0.484 | 0.656 | 0.738 | 0.611 |
| Qwen3.6-27B | 0.686 | 0.790 | 0.638 | **0.654** | **0.790** | **0.670** | 0.629 | 0.676 | 0.506 | 0.668 | 0.787 | 0.547 | 0.620 | 0.714 | 0.491 | 0.657 | 0.743 | 0.612 |
| Gemma 4 26B A4B | 0.677 | 0.780 | 0.634 | 0.627 | 0.769 | 0.636 | 0.629 | 0.679 | 0.513 | 0.650 | 0.776 | 0.528 | 0.618 | 0.709 | 0.474 | 0.640 | 0.731 | 0.588 |
| Gemma 4 31B | **0.710** | 0.806 | 0.673 | 0.619 | 0.740 | 0.624 | 0.621 | 0.667 | 0.499 | 0.649 | 0.773 | 0.525 | 0.637 | 0.726 | 0.517 | 0.656 | 0.741 | **0.614** |
| GPT-5.4<sup>†</sup> | 0.650 | 0.761 | 0.549 | 0.652 | 0.776 | 0.663 | **0.720** | **0.778** | **0.584** | **0.708** | **0.839** | **0.579** | 0.708 | 0.776 | 0.551 | **0.676** | **0.768** | 0.608 |

<sup>a</sup>Fit per cohort on the train+val split of the target panel (see methods). All other rows are zero-shot.

Against the trained baselines, the data support a shared range rather than an ordering, answering our first question. The two baselines divide the metrics between them. UNITO is stronger on mean F1 macro (0.690) and Hull IoU (0.665), whereas cyMAE is highest on balanced accuracy (0.773) yet has the lowest F1 macro of the methods compared (0.656). GPT-5.4 LLM-Gate trails UNITO on F1 macro (0.676 vs. 0.690) and Hull IoU (0.608 vs. 0.665) but exceeds it on balanced accuracy (0.768 vs. 0.757). A Wilcoxon signed-rank test over the 11 cohort means resolves none of these gaps, separating UNITO neither from GPT-5.4 LLM-Gate (*p* = 0.46/0.83/0.15 for F1 macro / BA / Hull IoU) nor from Qwen3.6-27B, the best open-weight backbone on F1 macro (*p* = 0.21/0.64/0.13). Supplemental section S6.5 reports the full tests. On all three metrics the best zero-shot configuration, given no panel-specific training, lands between the two baselines fit on every target panel. The trained baselines do retain a visible edge on the small, shallow-tree flow panels, where fitting per cohort pays off.

Two patterns hold across the table. *(i) Scale helps within the range we test.* Moving from 4B to 27–31B backbones improves every LLM formulation, though within the 27– 31B tier model family and generation matter more than parameter count. *(ii) Domain fine-tuning on the wrong modality hurts.* Cell-o1, fine-tuned from Qwen2.5-7B on transcriptomic data, ranks lowest in every formulation and trails even the 4B general-purpose backbones, suggesting that RNA-seq-tuned cell-type reasoning does not transfer to cytometry.

The case study in Figure 2 exposes a characteristic failure per formulation. LLM-Gate’s axis-aligned rectangles do not cleanly trace curved dense regions, clustering LLM-C2S misses easy, well-separated splits, and density-grid LLM-C2S struggles with transitional populations that lack a clear density peak.

**Figure 2.**
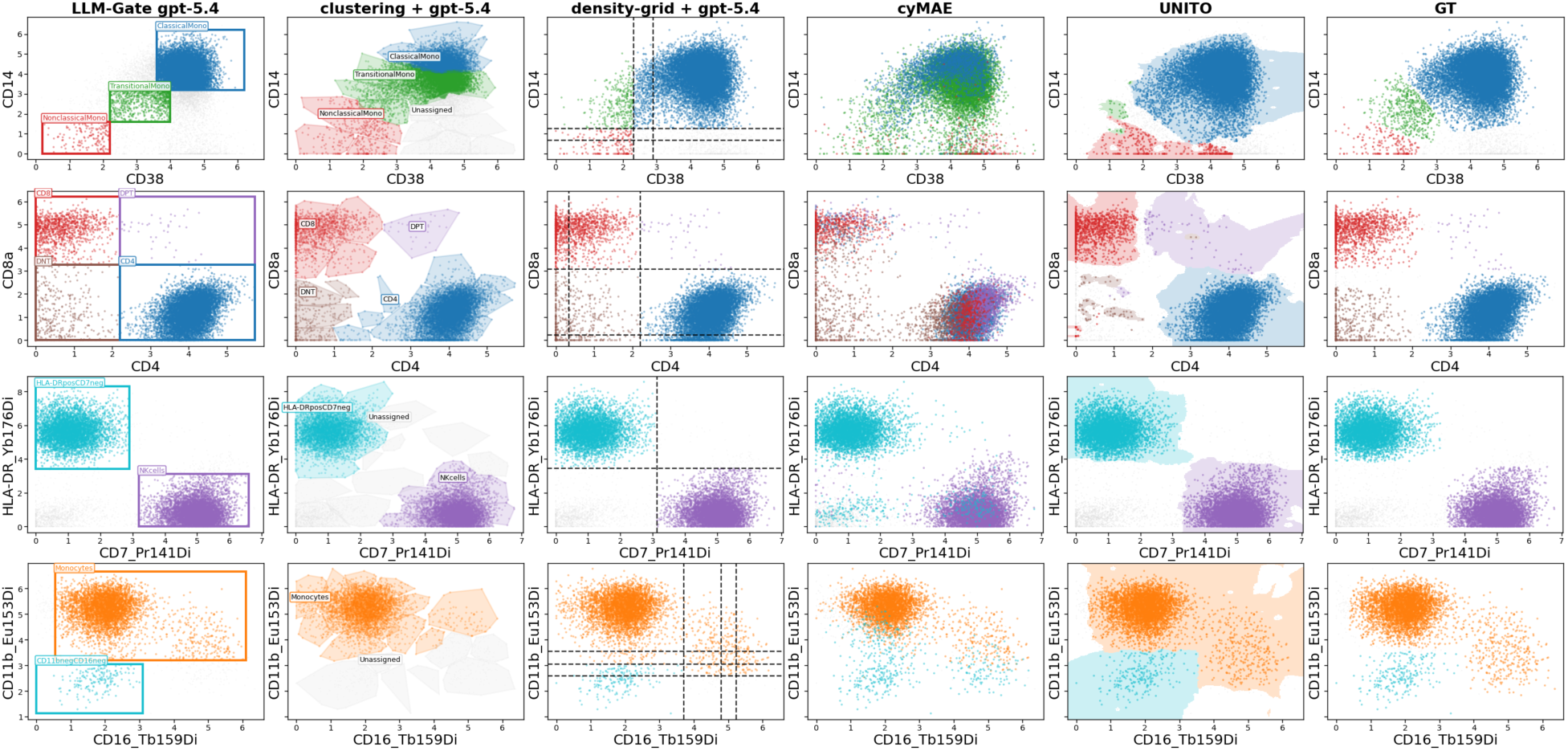
Each formulation fails in a characteristic way on hard steps. Rows: Acute2020 steps 18, 29 and Bjornson steps 06, 07 (full sample IDs in Figure S1). Columns (L→R): LLM-Gate, LLM-C2S (clustering, density-grid), cyMAE, UNITO (faint background = decision region), ground truth. Each panel shows one (sample, step) benchmark instance, with one point per cell of the parent population; LLM backbone: GPT-5.4.

### Vision input and self-verification loops induce a systematic gate-tightening bias

LLM-Gate presents the parent population as a one-dimensional histogram per axis, so the model never sees the joint structure an immunologist reads off a biaxial plot, that is, the clusters, gradients, and boundaries. We test whether supplying that two-dimensional shape directly helps, in two increments of visual grounding, on one closed-weight (GPT- 5.4) and one open-weight (Qwen3.5-27B) backbone.

**VLM-Gate** adds the image, extending the LLM-Gate prompt with a rendered two-dimensional scatter of the parent population in the (*x_v_*, *y_v_*) plane while leaving text and schema unchanged. **Agent-Gate** keeps that same prompt and schema and adds self-verification, a tool that re-renders the model’s proposed rectangle on the scatter so the model can inspect and revise its gate over up to four turns. Tool schema and turn- control details are in the supplemental information (section S7).

Adding visual grounding does not reliably improve annotation, in keeping with the trouble VLMs have reading charts faithfully^29^, and the self-verification loop degrades it on both backbones. On GPT-5.4 (Figure 3A), cohort-mean F1 macro is flat between LLM-Gate and VLM-Gate (both 0.676, with LLM-Gate first on 7 of 11 cohorts), while Agent-Gate falls −0.013 behind. Cohort-mean Hull IoU drops monotonically across the three variants (0.608 → 0.582 → 0.548), with LLM-Gate ranking first on 9 of the 11 cohorts and Agent-Gate on none. On the open-weight Qwen3.5-27B the Agent-Gate degradation replicates but the effect of the image reverses. Agent-Gate is again last on both metrics (−0.014 F1 macro, −0.037 Hull IoU relative to LLM-Gate), whereas VLM-Gate edges LLM-Gate on both (0.663 vs. 0.656; 0.617 vs. 0.613). Supplying the image is therefore backbone-dependent, while adding the self-verification loop is uniformly harmful.

**Figure 3.**
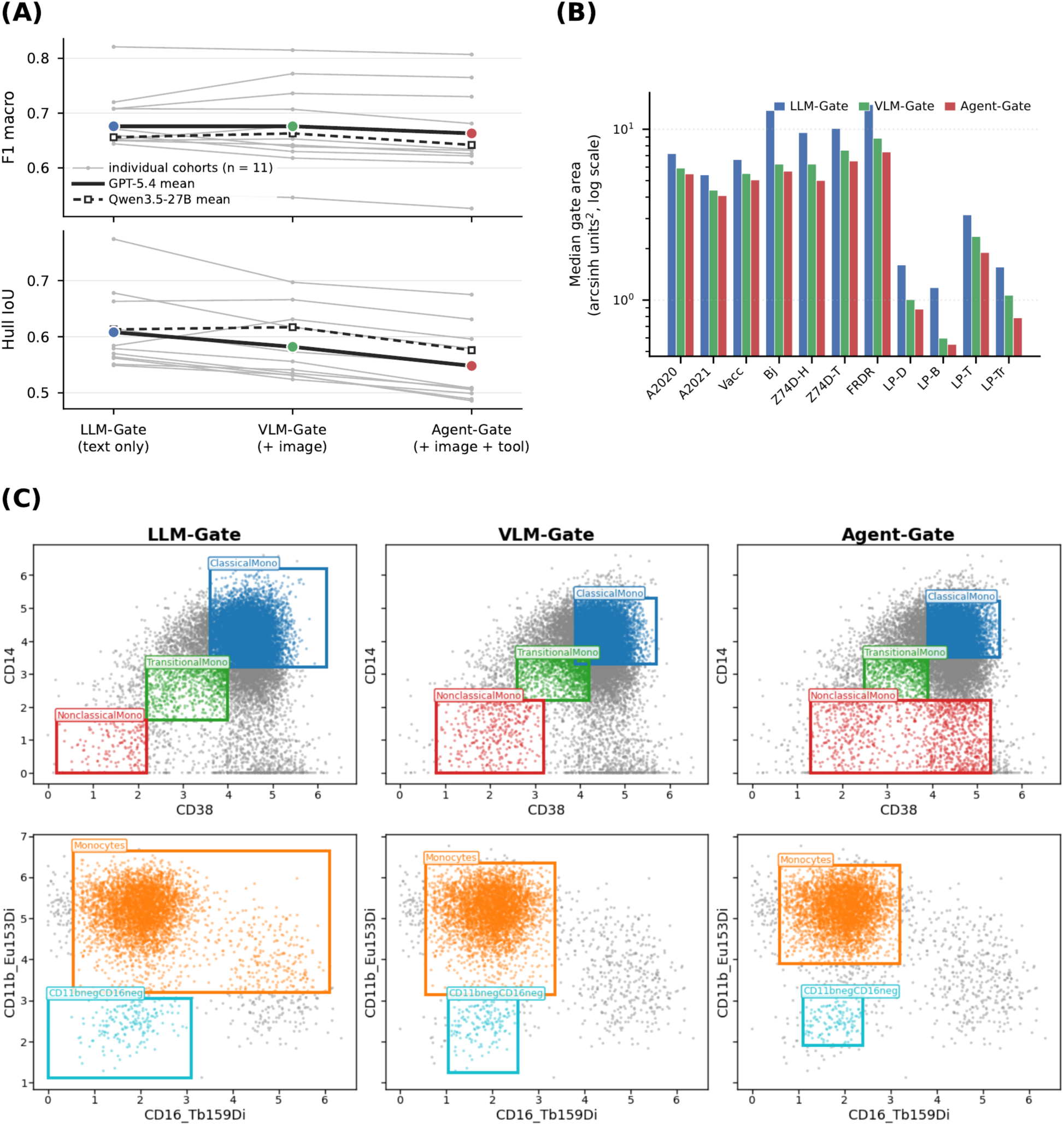
Visual and agentic feedback systematically tighten predicted gates. (A) Cohort-mean F1 macro (top) and Hull IoU (bottom) across the three Gate variants. Thin gray lines are the 11 individual cohorts on GPT-5.4; the solid line is their equal- weight mean and the dashed line the Qwen3.5-27B mean, for which only the across- cohort mean line is drawn (per-cohort values in Table S9). F1 macro is flat from LLM- Gate to VLM-Gate and falls only at Agent-Gate, whereas Hull IoU declines across all three. (B) Median predicted-rectangle area in the (*x_v_*, *y_v_*) arcsinh plane (log-scaled axis) per cohort, over the *n* = 23,041 (cohort, sample, step, category) instances on which all three methods returned a valid rectangle (GPT-5.4). Bars are medians of a paired comparison; the ordering LLM-Gate > VLM-Gate > Agent-Gate is monotone across all 11 cohorts. (C) Per-category rectangles on two representative steps, Acute2020 sample 994570 step 18 (CD38 × CD14) and Bjornson sample R15W11 step 7 (CD16 × CD11b). Gray = parent cells; colored = cells assigned to each candidate by rectangle containment, boundary in matching color (GPT-5.4). Cohort abbreviations as in Table 2. See also Table S10 and Figure S1, which shows all four case steps.

The falling Hull IoU traces to a systematic *gate-tightening* bias, consistent with the limits of unaided LLM self-correction^30^. Across 23,041 paired (cohort, sample, step, category) instances where all three methods returned a valid rectangle, median rectangle area in the (*x_v_*, *y_v_*) arcsinh plane falls to 0.86 × that of LLM-Gate for VLM-Gate, and to 0.79 × for Agent-Gate, with Agent-Gate smaller than LLM-Gate on 72% of paired instances. The shrinkage is monotone across all 11 cohorts (Figure 3B). The bias is not an artifact of the closed-weight backbone. On Qwen3.5-27B, Agent-Gate’s median rectangle is smaller than LLM-Gate’s in all 11 cohorts, by 3–62% (Table S10, supplemental section S7). A tighter rectangle truncates the periphery of the ground-truth cluster, which directly lowers Hull IoU. The truncated cells, however, default to *Unassigned* rather than being misrouted to another candidate, so label-level F1 takes only a mild penalty (Figure 3C; all four case steps in Figure S1).

### Distribution shape carries the dominant signal, with curated marker priors contributing

Turning to our second question, we ablate the LLM-Gate prompt to identify which components carry the signal. The prompt has three components, namely the curated marker priors (*D_v_*), the parent-distribution shape conveyed as one-dimensional histograms (*H*), and the peak/valley density landmarks (*P*). We remove one component at a time (Table 3), holding the rectangle-output formulation fixed.

**Table 3.**
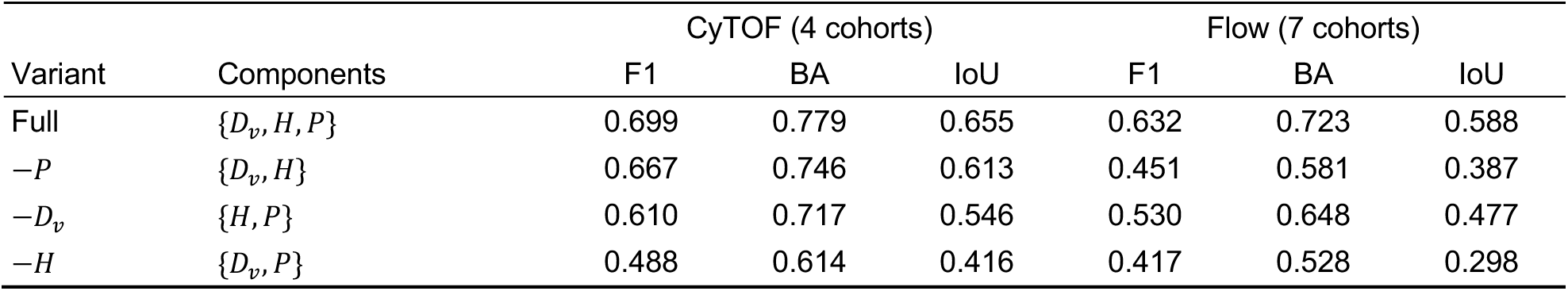
LLM-Gate input ablation. (Qwen3.6-27B). CyTOF = mean over Acute2020/2021/Vaccine, Bjornson; Flow = mean over seven flow cohorts (Z74D-H/T, FRDR, LP-T/Tr/B/D). *Full* is the LLM-Gate configuration of Table 2. *D_v_*, curated marker priors; *H*, per-axis histograms; *P*, peak/valley density landmarks. Cohort abbreviations as in Table 2.

Distribution shape *H* is the dominant signal in both modalities. Removing the histogram yields the largest drop (−0.211 on CyTOF, −0.215 on flow) (Table 3). Removing the curated marker priors *D_v_* costs a comparable amount on both modalities (−0.089 on CyTOF, −0.102 on flow), confirming that supplying each candidate’s expected marker levels carries signal beyond the marker-name knowledge stored in the model’s weights. The peak/valley landmarks *P* exhibit sharply modality-dependent behavior. Removing *P* has a small effect on CyTOF (−0.032) but produces a large drop on flow (−0.181), mirroring how chart and table reasoning hinges on reading the plotted values^29,31^. Flow panels have fewer candidates per step and sharper unimodal or bimodal distributions, so the explicit peak and valley coordinates act as direct gate-boundary cues that the histogram bars alone do not resolve.

### Zero-shot LLMs degrade less than trained baselines under distribution shift

In cytometry, distribution shift is the norm rather than the exception. Cohorts differ in batch, site, instrument, and patient population, so any cross-cohort method must stay reliable under it. Having localized LLM-Gate’s signal to distribution shape, we ask whether that signal survives such shift. We study two perturbations a panel-agnostic method must absorb without retraining. In **(a) population depletion**, the target candidate’s prevalence in the parent drops sharply. In **(b) calibration drift**, the same panel produces measurably different signal across batches. We test each with nine curated clinical or technical scenarios (supplemental section S8.1).

Each trained baseline is vulnerable to one perturbation family, whereas the LLM-Gate rows absorb both. At *ρ* = 1% depletion (Figure 4A), Qwen3.6-27B LLM-Gate retains the highest recall on the depleted candidate (median 0.91), ahead of cyMAE (0.86) and GPT-5.4 LLM-Gate (0.84), while UNITO drops furthest (median 0.73), including one scenario it fails at every severity (0.00 recall already at *ρ* = 100%, Table S13). On non-depleted candidates UNITO leads on median but drops to 0.03 on the rituximab outlier.

**Figure 4.**
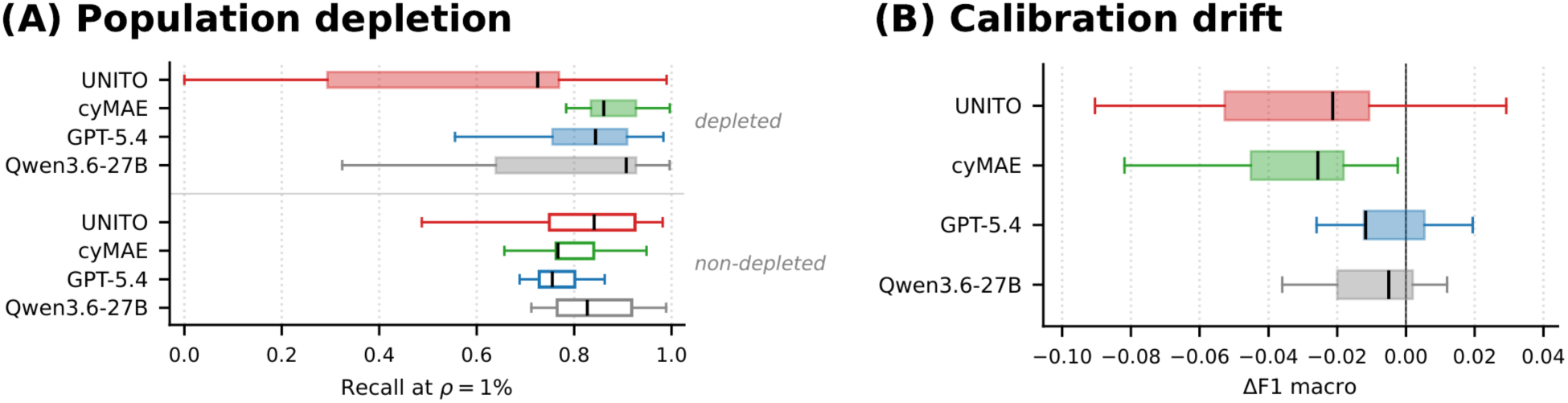
Zero-shot LLM-Gate absorbs both perturbation families that degrade the trained baselines. (A) Recall on the depleted candidate (filled boxes) versus the non-depleted candidates at the same gating step (hollow boxes), at depletion severity *ρ* = 1%. (B) ΔF1 macro under calibration drift (perturbed minus nominal). In both panels each box summarizes the nine curated scenarios of that perturbation family (*n* = 9 scenarios per box). Scenario definitions are in supplemental section S8.1. See also Tables S13 and S15.

Calibration drift reverses which baseline suffers (Figure 4B). Here cyMAE degrades on 9/9 scenarios (mean Δ*F*1 −0.040, worst −0.132 on lineage_dim) and UNITO on 8/9 (−0.027), while the two LLM-Gate rows degrade least (−0.014 each for GPT-5.4 and Qwen3.6-27B; per-scenario numbers in Table S15). Channel rescaling perturbs the per-cell embedding that cyMAE relies on and shifts the absolute intensities that UNITO’s fitted gates are pinned to, whereas LLM-Gate reads the current per-axis distribution and places its gates relative to it.

### Walking the gating tree stepwise outperforms flat annotation

Hierarchical gating improves panel transferability and interpretability, but its sequential structure accumulates error. Mistakes compound from one step to the next as depth grows (the per-step-to-leaf error bound is formalized in supplemental section S3, Corollary S2). We test whether this trade-off is favorable when the target is a full leaf-level label, comparing a flat and a hierarchical annotator on the same leaf vocabulary across all 11 cohorts and two backbones. The three cohorts sharing the deep 62-step tree are swept at three depths, namely L1 (16 lineage leaves), L2 (38 identity leaves), and L3 (the full 83-leaf tree), while the remaining cohorts are evaluated at their full native depth, giving 17 cohort-depth settings. Because a cell can carry several leaf labels at once, leaf annotation is fundamentally a multi-label task, and we report multi-label macro and micro F1.

*Flat C2S* (Cell-o1-style^21^) clusters the sample with FlowSOM^9^ (with *K* set to the leaf count at each depth) and labels each cluster in a single LLM call. The *LLM-Gate cascade* instead walks the gating tree, using its own predicted parent at each step, so no ground-truth labels leak. Both are run with GPT-5.4 and with the open-weight Qwen3.6-27B. The cell-sentence prompt is in supplemental section S9.1.

The cascade outperforms Flat C2S in 16 of the 17 cohort-depth settings, on both metrics and with both backbones (Figure 5A). FRDR COVID-19 is the single exception, where the flat harness wins on both metrics under both backbones. Error does accumulate with depth. On Acute2020’s deep tree, cascade F1 macro falls from 0.541 at L1 to 0.240 at L3 as the leaf vocabulary expands and the surviving leaves become intrinsically rarer. The degradation is nevertheless graceful rather than catastrophic. Cascade F1 micro over the same range stays nearly flat (0.843/0.813/0.805, the Acute2020 curve in Figure 5B) against Flat C2S’s 0.455/0.537/0.460 (Tables S17–S19). FlowSOM clusters cannot resolve the leaf vocabulary even when *K* is matched to the leaf count at each depth. This is the behavior Corollary S2 predicts, where depth enters the leaf-error bound as a multiplier rather than an exponent. Per-sample breakdowns, per-cell uniform manifold approximation and projection (UMAP) plots, confusion matrices, and cascade traces are in supplemental section S9.

**Figure 5.**
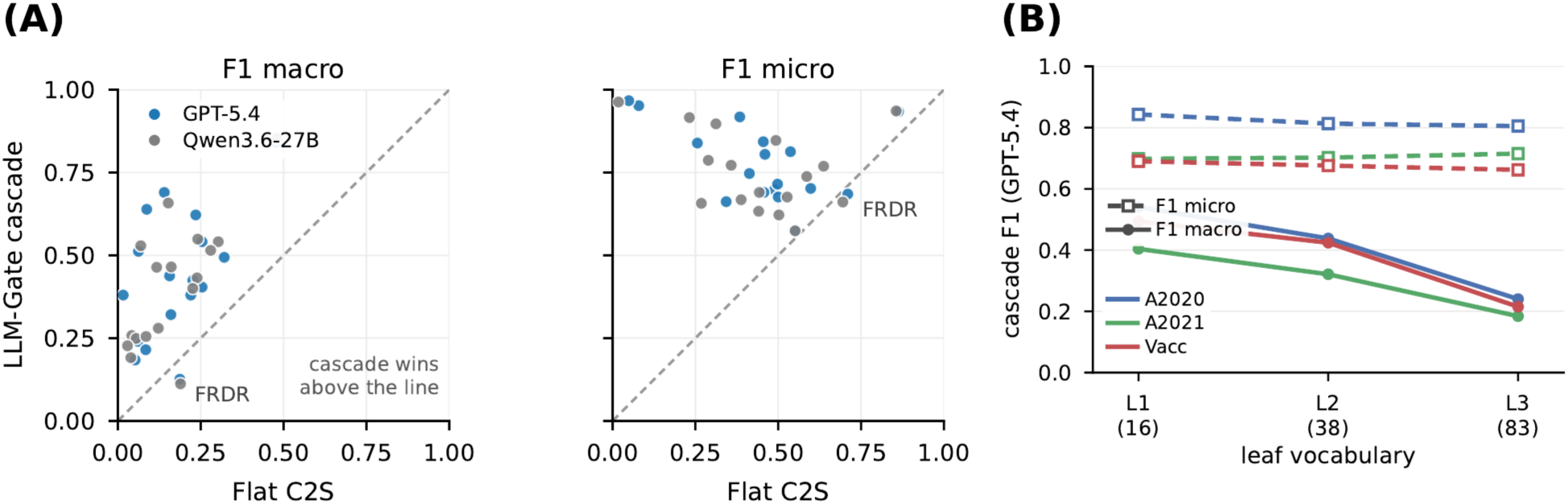
Walking the gating tree stepwise outperforms flat annotation. (A) Leaf-level multi-label F1 macro (left) and F1 micro (right) for Flat C2S (*x*) against the LLM-Gate cascade (*y*) on each of the 17 cohort-depth settings, for both backbones; points above the dashed identity line are settings the cascade wins. The cascade wins 16 of 17 settings on both metrics and both backbones; FRDR COVID-19, labeled, is the single exception. (B) Cascade F1 across the three depth cuts of the 62-step tree, for the three cohorts that share it (GPT-5.4). F1 macro falls as the leaf vocabulary expands from 16 to 83 leaves, whereas F1 micro stays nearly flat, so the accumulated error is concentrated in the rare leaves that macro-averaging weights equally. See also supplemental section S9.

## DISCUSSION

Off-the-shelf models, given no cytometry-specific training, place gates in the range of models fit to every target panel, and we read this as a statement about the task rather than about the models. Once an expert gating tree fixes what each step is asking (which cells are in play, which two markers to read, which children are on offer), what remains is largely a reading of the shape in front of the model. That reading is something a general-purpose model can already do. CytoGate-Bench makes it measurable, and the picture holds across six backbones and 11 cohorts.

The same reading explains the robustness result. A model that places its gates relative to the distribution it is currently shown has nothing pinned to absolute intensities or to a fitted embedding. Those anchors are what fail when a population is depleted or a channel drifts, and each trained baseline breaks on the perturbation that invalidates its own fit. It explains why the cell-sentence formulation trails. Serializing a cluster into a list of markers discards the shape that carries the signal, so pairing that serialization with an explicit shape summary is an obvious extension. It also explains why walking the tree outperforms annotating flatly. A flat call must settle the granularity of the answer and the distinctions among cell types at once, whereas each step of the cascade is a single bounded question posed on a view an expert would recognize.

What the reading does not explain is the effect of adding vision. Showing the model the plot it has been describing in numbers, or letting it re-render and revise its own boundary, moves its gates inward rather than into better places, and the shrinkage holds on both backbones we test. Self-verification here buys conservatism, not accuracy, and calibrating it is a prerequisite for visually grounded gating. Beyond cytometry, the bias is a clean case of vision-text disagreement hurting a numerically grounded task, in which handing a text-only model a rendered image of the data made it more conservative rather than more accurate.

For cytometry practice, LLM-based per-step gating under a given expert tree keeps annotation auditable. Every cell’s label is the composition of local, inspectable decisions on standard biaxial views, the same trace an immunologist would review. The zero-shot regime evaluated here is also a floor rather than a ceiling for what LLMs can offer. We deliberately withhold panel-specific training to test transferability, and that regime matches the setting laboratories face as panels evolve across studies. Where labeled data exist, cytometry-aware training of an LLM on top of the per-step formulation could push accuracy further without giving up the audit trail.

### Limitations of the study

Two limits govern how the reported scores should be read. The first is the ground truth itself. Our labels come from one expert gating per cohort, and expert gating is not perfectly reproducible. In a published comparison of five independent raters annotating a shared peripheral blood mononuclear cell (PBMC) library^13^, pairwise agreement ranged from *κ* = 0.44 to 0.86 and fell to 0.42 in the deeper layers of the tree. The same study’s raters also disagreed on which populations to gate at all, annotating 39 versus 24 subpopulations, with several gated by a single rater. Supplying the gating tree removes that coarser source of disagreement and leaves the within-step boundary decision, which is what we score, so absolute scores should be read against this residual disagreement rather than against 1.0. The second is statistical power. The cross-cohort comparison rests on a Wilcoxon signed-rank test over only 11 paired cohort means. The test separates the best LLM from UNITO on none of the three metrics, but at *n* = 11 that is a failure to detect a difference rather than evidence of equivalence.

Several choices bound the scope of the claims. Methods receive the expert-defined gating tree as input, so we do not evaluate whether a method can construct the gating strategy itself. The LLM-based methods run under a deliberately minimal harness (off-the-shelf models, no fine-tuning, axis-aligned rectangular gates, and one generation attempt per step), so the reported numbers are a floor rather than a ceiling. The visual- modality results, VLM-Gate and Agent-Gate, are obtained on two backbones, one closed- and one open-weight, rather than across the VLM family. The two agree that the self-verification loop tightens gates and lowers Hull IoU, but disagree on whether the image alone helps. The closed-weight GPT-5.4 rows additionally depend on a commercial API and may not reproduce exactly as the served model is updated. The open-weight rows are reproducible from pinned checkpoints, with their seed variance reported in Table S5.

Finally, the benchmark is scoped to per-step annotation accuracy in cytometry. It does not yet probe downstream translational use, such as rare cell-type detection or biomarker discovery in a clinical study. The per-step, two-marker gating primitive is also specific to cytometry and does not transfer directly to untargeted high-plex assays such as scRNA-seq, which type cells by whole-transcriptome clustering and multi-gene signatures instead. Cost is a practical limit of its own, since the deepest gating trees require one LLM call per tree node, which amounts to a median of roughly 4.6 minutes per sample at the full 62-step depth (supplemental section S6.6).

## METHODS

### Cohort curation and provenance

The 11 cohorts are re-curated public flow / mass-cytometry studies, each released through one of four manual-gating ecosystems (OMIQ workflow, FlowJo WSP, R `flowWorkspace GatingSet`, or pre-computed label CSV). Table S3 records the source repository, gating tool, and number of original FCS samples per cohort, together with the per-cohort exclusions imposed during harmonization.

### Preprocessing

All antibody / fluorophore-conjugated channels (*protein*), together with viability dye, DNA intercalator, normalization-bead, Gaussian event-shape, and background isotope channels, are transformed by arcsinh(*x*/cofactor) before storage. The cofactor is platform-specific, 5 for mass cytometry (CyTOF) and 150 for conventional flow cytometry, following the standard cytometry conventions for each modality. Light-scatter (`FSC-*`, `SSC-*`) and technical (`Time`, `Event_length`) channels are not arcsinh-transformed. Instead, they are clipped to the per-sample [1,99] percentile range and min-max scaled to [0,10]. All channels are stored as `float16` parquet columns with zstd compression, identical across cohorts. The cofactor used for each cohort is recorded per cohort alongside the channel-type assignment, and all gate-polygon vertex coordinates are stored in the same arcsinh space, so no further transform is needed at evaluation time to align gates with cells.

Mass cytometry uses isotopic mass tags rather than fluorophores and therefore has no spectral spillover. The CyTOF cohorts (Acute2020/2021, Vaccine, Bjornson) contribute compensation-free expression. The conventional-flow cohorts inherit per-cohort compensation matrices from their source workspaces. Cohorts harvested from FlowJo workspaces (FRDR COVID-19, FR-FCM-Z74D) are extracted with `flowWorkspace` in R, which applies the workspace-stored compensation matrix to the linear-scale FCS values before our arcsinh transform. Lyoplate enters our pipeline through legacy `GatingSetList` objects whose stored expression is already compensated. The Acute2020/2021 and Vaccine OMIQ workflows include manual cleanup gates upstream of cell-type annotation (bead removal, debris, doublet, viability), which are recorded as the first three to seven steps of each cohort’s gating tree rather than discarded. Per-step accuracy thus rewards methods that handle cleanup gates correctly.

### Per-step task definition

#### Notation

Let C denote the set of 11 curated cohorts. For each cohort *c* ∈ C:

- *M_c_* is the antibody-panel marker set, with |*M_c_*| ∈ [8,40].
- *T_c_* is the expert-specified gating tree, a rooted tree whose root *r_c_* represents the full sample, and where each internal node *v* has a finite set of children *C_v_*, the *candidate cell types* at *v*.
- Each internal node *v* carries an ordered marker pair (*x_v_*, *y_v_*) ∈ *M_c_* × *M_c_* and *curated cell-type-specific marker priors D_v_* = {*d_k_*}*_k_*_∈_*_Cv_*, one textual description per candidate stating its expected phenotype on the focal markers (e.g., classical monocytes as CD38 high, CD14 high).
- *S_c_* is the sample set, where each sample *s* ∈ *S_c_* has cell count *n_s_* and expression matrix *X_s_* ∈ ℝ*^ns^*^×|*M*^*^c^*^|^.

For each internal node *v* and sample *s*, the *parent population P_v_*_,*s*_ ⊆ {1, …, *n_s_*} is the set of cell indices previously assigned to *v*. The expert provides a per-cell ground-truth label for this local step:

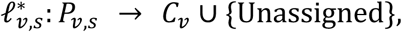

where Unassigned absorbs cells the expert declines to place in any candidate cell type (e.g., ambiguous, borderline, or residual signals).

#### Benchmark instance

A benchmark instance is a triple *τ* = (*c*, *s*, *v*) specifying a cohort, a sample, and an internal node. Each instance is stored as a standalone record with four canonical fields:

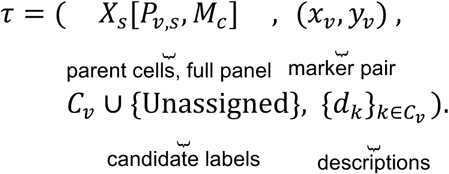

The method receives the full panel *M_c_* together with the focal marker pair (*x_v_*, *y_v_*) that defines the step and a *textual description d_k_* per candidate, each curated from the heuristics immunologists apply when gating on biaxial plots (Figure 1B).

#### Task

Given the four fields of *τ* (parent cells, marker pair, candidate labels, textual descriptions), a method *f* produces a per-cell prediction:

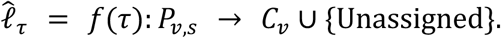

The method must assign every parent cell to one of the candidate labels.

#### Composition

If every per-step rule along a cell’s root-to-leaf path returns the next node on that path, walking the tree returns that leaf (Proposition S1). Composing per-step decisions therefore recovers the leaf-level label a flat annotator emits, and conversely every flat classifier is realizable per-step, so the formulation costs no expressiveness (Corollary S3; proofs in supplemental section S3).

#### Scope

The gating tree is an input, not an output. Throughout, *panel-agnostic* means that a method needs no panel-specific training to route cells under a tree it is given. It does not mean discovering populations the tree does not name. Constructing the tree itself is outside the benchmark’s scope.

### Gate extraction and per-cell ground truth

Gate definitions are harvested from each cohort’s source format and serialized to a uniform per-sample JSON keyed by step index, with each step listing one or more gates as a polygon (or rectangle expanded to four vertices) in the same arcsinh-transformed marker space as the stored expression columns. Three extraction paths are used (OMIQ REST API; FlowJo WSP parsed with `flowWorkspace` in R; legacy `GatingSetList` objects), detailed in supplemental section S2.5. The extracted per-sample gates are validated against the gating tree by checking that every step’s parent expression resolves to a well-defined parent mask and that polygon vertex coordinates lie within the per-sample [1,99] percentile range of their channel. Samples whose gates fail to extract, for example through a missing ground-truth export or a filename mismatch, or that contain a degenerate gate are excluded (Table S3, “Exclusions” column). A step whose parent population is empty in an otherwise valid sample is not an exclusion. The sample is retained and that step contributes no evaluation instance, as noted in the Table 1 caption.

The gating tree defines one annotation step per (parent population, marker pair) combination. For each cell that lies in a step’s parent mask, the cell is checked against every child gate of that step in order, and the first matching child becomes the cell’s per-step label. Cells in the parent that match no listed child are left as missing (`NaN`) for that step and propagate to *Unassigned* downstream, as specified in the task definition above. We follow a strict *gate-positive only* convention. Only categories with an explicit gate in the source data appear in the gating tree, and we do not invent derived negative categories (*e.g.*, *Dead*, *Doublets*, *Other*, *Non-Treg*) for unmatched cells. The single exception is when a negative subset is itself used as a downstream parent (e.g., `non_Tfh` parents step 6 in some panels), in which case the source must already provide an explicit gate for it. This discipline ensures every non-NaN label in our parquet outputs is grounded in the original source-recorded gating, never inferred from omission, and that per-step accuracy is computed against the same labels the original analyst would have inspected.

Samples are partitioned into donor-stratified train/val/test splits targeting a 1:1:2 ratio (seed 42). Realized split sizes deviate from the target where donor grouping constrains the assignment, as in the Lyoplate panels, or where the cohort is small, as in FR-FCM- Z74D HC. Per-cohort channel counts and split sizes are in Table S2 (supplemental section S2.1).

### LLM-C2S and LLM-Gate implementations

Both LLM-class methods hold the rest of the per-step task fixed (the candidate set *C_v_*, the descriptions *D_v_*, the marker pair (*x_v_*, *y_v_*), and the system prompt) and differ only in how the parent population *P_v_*_,*s*_ is encoded for the model. LLM-C2S clusters the parent population in the (*x_v_*, *y_v_*) plane with one of two primitives, the *density-grid* variant (flowDensity^16^) and the *clustering* variant (FlowSOM^9^), which form the two row families of Table 2. It then serializes each cluster’s representative cell as a cell sentence^17^ and labels all clusters of a step in a single call, following Cell-o1^21^. Cluster labels then propagate to member cells. LLM-Gate summarizes the parent population as one histogram per axis, given as text, together with kernel-density peak/valley landmarks, and prompts the model to output one axis-aligned rectangle per candidate. Each cell takes the label of the rectangle containing it, the smallest such rectangle when several overlap, and cells covered by no rectangle default to *Unassigned*. The verbatim production prompts, the JSON output schema validated against, and the per-paradigm output-cardinality and propagation rules are reproduced in supplemental sections S4 and S5.

### Baselines, backbones, and training regime

Table S4 lists the three method groups compared throughout the results, together with the input view each method consumes per step, whether it is fit on cohort labels or evaluated zero-shot, and the backbones used. The 1:1:2 train/val/test partition is donor-stratified with seed 42 (supplemental section S2.1, Table S2). The *trained baselines* (UNITO, cyMAE) are fit per cohort on train+val and evaluated on test, while *LLM-class methods* (LLM-C2S, LLM-Gate, VLM-Gate, Agent-Gate) see no labels and operate purely on the per-step inputs (*P_v_*_,*s*_, *x_v_*, *y_v_*, *C_v_*, *D_v_*) formalized in the per-step task definition above.

Cell-o1 (7B)^21^ is a Qwen2.5-7B fine-tuned on transcriptomic cell-type reasoning. It is included as a domain-tuned reference and is the only fine-tuned LLM in the roster.

### Evaluation metrics and aggregation

Per-cohort scores are computed by averaging per-step F1 macro, balanced accuracy (BA), and Hull IoU over all (sample, step) instances in the cohort’s test split. The *Mean* column of Table 2 averages these per-cohort scores with equal weight across the 11 cohorts (cohort, not task, is the aggregation unit, so panel diversity is not dominated by the sample-heavy CyTOF cohorts). F1 macro is the unweighted mean of per-candidate F1 over *C_v_* ∪ {Unassigned}. BA is the mean of per-candidate recall over the candidates of that set with nonzero ground-truth support in the instance, following the scikit-learn convention. Hull IoU is the intersection-over-union between the convex hulls of predicted and ground-truth cells assigned to the same candidate in the (*x_v_*, *y_v_*) arcsinh plane, with each point set trimmed to its per-axis [1,99] percentile range before the hull is computed to reduce outlier sensitivity.

### Inference and generation settings

All LLM backbones are run zero-shot, with a maximum completion budget of 32768 tokens and a single generation attempt per step. On a parse failure every cell of the step is predicted *Unassigned* (supplemental section S5). Open-weight reasoning backbones (Qwen3.5/3.6, Gemma 4) additionally receive a thinking-token budget of 8096 (`tb8096` variant). Decoding follows each backbone’s recommended setting. The closed-weight GPT-5.4 is greedy (temperature 0), whereas the open-weight reasoning backbones sample at temperature 1.0, top-*p* 0.95, and top-*k* 20 (Qwen) / 64 (Gemma), since greedy decoding degrades these reasoning modes. The open-weight rows are therefore stochastic. We verify their seed-stability in Table S5, where the cohort-mean F1 macro shifts by at most ±0.01 across 10 seeds. These generation settings are shared by every LLM-class row of Table 2, by the visual-modality variants VLM-Gate and Agent-Gate (tool-loop schedule in the supplemental information, section S7), and by the flat-vs. hierarchical comparison (supplemental section S9), whose only experiment-specific knob is the FlowSOM clustering granularity (*K* set to the leaf count per depth, 16/38/83). Full LLM-C2S and LLM-Gate prompts, including the JSON output schema validated against, are in supplemental sections S4.2 and S4.1. Per-paradigm output cardinality and the per-cell propagation rule (rectangle containment, cluster membership, *Unassigned* residual) are summarized in supplemental section S5.

### Robustness protocol

Two perturbation families are applied to the test split without retraining any method. In population depletion, the target candidate’s prevalence in the parent is subsampled toward *ρ* ∈ {100%, 10%, 1%} of nominal. In calibration drift, channel signal is rescaled to emulate batch-to-batch variation. Each family comprises nine curated clinical or technical scenarios. Scenario definitions, the scenario × cohort grid, and full per-scenario results are in supplemental section S8.

### Flat-vs.-hierarchical protocol

The flat and hierarchical annotators share one backbone per run (GPT-5.4 or Qwen3.6- 27B), one canonical input parquet per sample, and one leaf vocabulary derived from the same gating tree. Flat C2S clusters each sample with FlowSOM (*K* set to the leaf count at each depth) and labels every cluster in a single LLM call. The LLM-Gate cascade walks the gating tree using its own predicted parent at each step, so no ground-truth labels leak. Leaf-level annotation is scored as a multi-label task with multi-label F1 (macro/micro). Pipeline details, the flat cell-sentence prompt, and per-sample breakdowns are in supplemental section S9.

### Statistical analysis

The *Mean* column of Table 2 averages 11 per-cohort scores, so the cohort is the natural unit for a paired test. We compare each LLM-Gate backbone against UNITO, the stronger trained baseline on F1 macro and Hull IoU, with a two-sided Wilcoxon signed-rank test on the 11 paired cohort means (*n* = 11). Table S6 reports the result. No metric separates the two method classes at *α* = 0.05. With *n* = 11 the test’s power is limited, and a null result does not establish equivalence. We therefore describe the comparison as an unresolved difference on the chosen metrics rather than as a win for either side.

### Compute and hardware

All open-weight LLMs are served with vLLM^32^ on a single node of eight NVIDIA H200 GPUs (141 GB each). The closed-weight model (GPT-5.4) is accessed through its vendor API and uses no local accelerators. The generation settings of the previous subsection are held fixed across this hardware. As a representative cost, a full zero-shot evaluation sweep with Qwen3.6-27B completes in roughly 2 hours of wall-clock on this node (≈16 GPU-hours). The trained baselines are fit and evaluated on separate nodes, UNITO on 4 × H200 in about 2 hours total (≈8 GPU-hours), and cyMAE on 4 × A100 in about 6 hours total (≈24 GPU-hours).

### Ethics

All 11 cohorts are publicly released human-subject cytometry studies. Institutional review board (IRB) approval and informed-consent context are documented in each cohort’s original publication. Supplemental section S2.2 (Table S3) lists the per-cohort source publications. CytoGate-Bench re-curates only the published cell-by-marker matrices and expert gating annotations, and no new human-subject data was collected for this work. Each cohort is used under its original license or data-use agreement. We redistribute harmonized data for all 11 cohorts, releasing the openly licensed cohorts under their original licenses and the access-controlled Lyoplate cohorts under custom terms commensurate with the ImmPort user agreement. Per-cohort licenses and our release terms are detailed in supplemental section S2.2. We see no significant potential risks. All source cohorts are de-identified public human-subject data, so re-curating them introduces no re-identification risk and no new personal data is collected.

Further details regarding the methods can be found in the supplemental information.

## RESOURCE AVAILABILITY

### Lead contact

Requests for further information and resources should be directed to and will be fulfilled by the lead contact, Dokyoon Kim.

### Materials availability

This study did not generate new materials.

### Data and code availability

- The harmonized per-step benchmark data (arcsinh cell-by-marker matrices, per- step annotations, gating trees, and the frozen evaluation splits) for all 11 cohorts have been deposited in a single Harvard Dataverse dataset at https://dataverse.harvard.edu/dataset.xhtml?persistentId=doi:10.7910/DVN/RDW4UL and are publicly available as of the date of publication. The dataset carries custom terms that state each cohort’s licensing (Acute2020/2021 and Vaccine: CC-BY 4.0; FRDR COVID-19: CC0 1.0; FR-FCM-Z74D and Bjornson: FlowRepository open data-sharing terms). The four HIPC Lyoplate cohorts, sourced from ImmPort/ImmuneSpace, are included under terms commensurate with the ImmPort user agreement (no re-identification, attribution to ImmPort and the source study, and redistribution only under the same terms). We additionally release extraction scripts that rebuild the per-step Lyoplate data from the source FCS files for users who prefer to obtain them directly under that agreement. Our added annotation layer (per-step annotations, gating trees, evaluation splits) is released under CC-BY 4.0.
- All original evaluation and harmonization code has been deposited at https://github.com/leebyounghan/cytof-qa-pipeline and is publicly available as of the date of publication under the MIT license.
- Any additional information required to reanalyze the data reported in this paper is available from the lead contact upon request.

## AUTHOR CONTRIBUTIONS

Conceptualization, J.K., B.L., K.-A.S., and D.K.; methodology, J.K. and B.L.; software, J.K. and B.L.; investigation, J.K. and B.L.; data curation, J.K. and B.L.; formal analysis, J.K. and B.L.; writing – original draft, J.K. and B.L.; writing – review & editing, J.K., B.L., N.A., M.I., M.L.M., M.E.L., C.-U.J., S.A.A., A.E.B., Shwetank, A.R.G., E.J.W., K.-A.S., and D.K.; supervision, K.-A.S. and D.K.; funding acquisition, K.-A.S. and D.K.

## DECLARATION OF GENERATIVE AI AND AI-ASSISTED TECHNOLOGIES

During the preparation of this work, the authors used Claude (Anthropic) to assist with editing and reformatting the manuscript. After using this tool, the authors reviewed and edited the content as needed and take full responsibility for the content of the published article.

## SUPPLEMENTAL INFORMATION

Document S1. Supplemental methods and notes (sections S1–S10), Figures S1–S10, and Tables S1–S19.

## REFERENCES

1. Lu, C., Lu, C., Lange, R.T., Yamada, Y., Hu, S., Foerster, J., Ha, D., and Clune, J. (2026). Towards end-to-end automation of AI research. Nature 651, 914–919. 10.1038/s41586-026-10265-5.

2. Huang, K., Zhang, S., Wang, H., Qu, Y., Lu, Y., Roohani, Y., Li, R., Qiu, L., Li, G., Zhang, J., et al. (2025). Biomni: A general-purpose biomedical AI agent. bioRxiv. 10.1101/2025.05.30.656746.

3. Alber, S., Chen, B., Sun, E., Isakova, A., Wilk, A.J., and Zou, J. (2025). CellVoyager: AI CompBio agent generates new insights by autonomously analyzing biological data. bioRxiv. 10.1101/2025.06.03.657517.

4. Li, H., Shaham, U., Stanton, K.P., Yao, Y., Montgomery, R.R., and Kluger, Y. (2017). Gating mass cytometry data by deep learning. Bioinformatics 33, 3423–3430. 10.1093/bioinformatics/btx448.

5. Arvaniti, E., and Claassen, M. (2017). Sensitive detection of rare disease-associated cell subsets via representation learning. Nature Communications 8, 14825. 10.1038/ncomms14825.

6. Kaushik, A., Dunham, D., He, Z., Manohar, M., Desai, M., Nadeau, K.C., and Andorf, S. (2021). CyAnno: A semi-automated approach for cell type annotation of mass cytometry datasets. Bioinformatics 37, 4164–4171. 10.1093/bioinformatics/btab409.

7. Kim, J., Ionita, M., Lee, M., McKeague, M.L., Pattekar, A., Painter, M.M., Wagenaar, J., Truong, V., Norton, D.T., Mathew, D., et al. (2024). Cytometry masked autoencoder: An accurate and interpretable automated immunophenotyper. Cell Reports Medicine 5, 101808. 10.1016/j.xcrm.2024.101808.

8. Ding, S., Bhattacharya, S., and Butte, A.J. (2025). ImmuneFM: Pre-training foundation model from cytometry data for immunology research. bioRxiv. 10.1101/2025.07.09.664020.

9. Van Gassen, S., Callebaut, B., Van Helden, M.J., Lambrecht, B.N., Demeester, P., Dhaene, T., and Saeys, Y. (2015). FlowSOM: Using self-organizing maps for visualization and interpretation of cytometry data. Cytometry Part A 87, 636–645. 10.1002/cyto.a.22625.

10. Levine, J.H., Simonds, E.F., Bendall, S.C., Davis, K.L., Amir, E.D., Tadmor, M.D., Litvin, O., Fienberg, H.G., Jager, A., Zunder, E.R., et al. (2015). Data-driven phenotypic dissection of AML reveals progenitor-like cells that correlate with prognosis. Cell 162, 184–197. 10.1016/j.cell.2015.05.047.

11. Aghaeepour, N., Finak, G., Consortium, F., Consortium, D., Hoos, H., Mosmann, T.R., Brinkman, R., Gottardo, R., and Scheuermann, R.H. (2013). Critical assessment of automated flow cytometry data analysis techniques. Nature Methods 10, 228–238. 10.1038/nmeth.2365.

12. Martini, P., Mohammadi, M., Thrun, M.C., Blumenthal, D.B., and Krause, S.W. (2026). Towards automated gating of clinical flow cytometry data. bioRxiv. 10.64898/2026.01.10.698765.

13. Liu, P., Pan, Y., Chang, H.-C., Wang, W., Fang, Y., Xue, X., Zou, J., Toothaker, J.M., Olaloye, O., Gonzalez Santiago, E., et al. (2025). Comprehensive evaluation and practical guideline of gating methods for high-dimensional cytometry data: Manual gating, unsupervised clustering, and auto-gating. Briefings in Bioinformatics 26, bbae633. 10.1093/bib/bbae633.

14. Chen, J., Ionita, M., Feng, Y., Lu, Y., Orzechowski, P., Garai, S., Hassinger, K., Bao, J., Wen, J., Duong-Tran, D., et al. (2025). Automated cytometric gating with human-level performance using bivariate segmentation. Nature Communications 16, 1576. 10.1038/s41467-025-56622-2.

15. Montante, S., Yokosawa, D., Li, L., Butyaev, A., Malek, M., Movassaghi, R., Michalchuk, Q., Hsu, C.-T.J., Shmil, D., Rahim, A., et al. (2025). Citizen science gamers enable automated flow cytometry gating through machine learning. bioRxiv. 10.1101/2025.10.07.679685.

16. Malek, M., Taghiyar, M.J., Chong, L., Finak, G., Gottardo, R., and Brinkman, R.R. (2015). flowDensity: Reproducing manual gating of flow cytometry data by automated density-based cell population identification. Bioinformatics 31, 606–607. 10.1093/bioinformatics/btu677.

17. Levine, D., Rizvi, S.A., Lévy, S., Pallikkavaliyaveetil, N., Zhang, D., Chen, X., Ghadermarzi, S., Wu, R., Zheng, Z., Vrkic, I., et al. (2024). Cell2Sentence: Teaching large language models the language of biology. In Proceedings of the 41st international conference on machine learning (ICML) Proceedings of machine learning research., pp. 27299–27325.

18. Rizvi, S.A., Levine, D., Patel, A., Zhang, S., Wang, E., Perry, C.J., Constante, N.M., He, S., Zhang, D., Tang, C., et al. (2025). Scaling large language models for next-generation single-cell analysis. bioRxiv. 10.1101/2025.04.14.648850.

19. Zhang, F., Liu, T., Zhu, Z., Wu, H., Wang, H., Zhou, D., Zheng, Y., Wang, K., Wu, X., and Heng, P.-A. (2025). CellVerse: Do large language models really understand cell biology? In Advances in neural information processing systems 38 (NeurIPS 2025) datasets and benchmarks track.

20. Hou, W., and Ji, Z. (2024). Assessing GPT-4 for cell type annotation in single-cell RNA-seq analysis. Nature Methods 21, 1462–1465. 10.1038/s41592-024-02235-4.

21. Fang, Y., Jin, Q., Xiong, G., Jin, B., Zhong, X., Ouyang, S., Yang, Y., Zhang, A., Han, J., and Lu, Z. (2026). Cell-o1: Training LLMs to solve single-cell reasoning puzzles with reinforcement learning. Bioinformatics, btag208.

22. Xiao, Y., Liu, J., Zheng, Y., Xie, X., Hao, J., Li, M., Wang, R., Ni, F., Li, Y., Luo, J., et al. (2024). CellAgent: An LLM-driven multi-agent framework for automated single-cell data analysis. arXiv preprint arXiv:2407.09811.

23. Zhang, H., Sun, Y.H., Hu, W., Cui, X., Ouyang, Z., Cheng, D., Zhang, X., and Zhang, B. (2025). CompBioAgent: An LLM-powered agent for single-cell RNA-seq data exploration. bioRxiv. 10.1101/2025.03.17.643771.

24. Huang, D., Li, H., Li, W., Zhang, H., Dickson, P., Zhan, M., Miller, J.P., Cruchaga, C., Province, M., Chen, Y., et al. (2025). OmniCellAgent: Towards AI co-scientists for scientific discovery in precision medicine. bioRxiv. 10.1101/2025.07.31.667797.

25. Bu, D., Sun, J., Li, K., He, Z., Huang, W., Hu, J., Zhang, S., Lei, S., Huo, P., Wang, Z., et al. (2026). Empowering AI data scientists using a multi-agent LLM framework with self-evolving capabilities for autonomous, tool-aware biomedical data analyses. Nature Biomedical Engineering. 10.1038/s41551-026-01634-6.

26. Mehandru, N., Hall, A.K., Melnichenko, O., Dubinina, Y., Tsirulnikov, D., Bamman, D., Alaa, A., Saponas, S., and Malladi, V.S. (2025). BioAgents: Bridging the gap in bioinformatics analysis with multi-agent systems. Scientific Reports 15, 39036. 10.1038/s41598-025-25919-z.

27. Dip, S.A., Zafor, A., Paul, B.K., Shuvo, U.A., Emon, M.I., Wang, X., and Zhang, L. (2025). LLM4Cell: A survey of large language and agentic models for single-cell biology. arXiv preprint arXiv:2510.07793.

28. Liu, F., Eisenschlos, J., Piccinno, F., Krichene, S., Pang, C., Lee, K., Joshi, M., Chen, W., Collier, N., and Altun, Y. (2023). DePlot: One-shot visual language reasoning by plot-to-table translation. In Findings of the association for computational linguistics: ACL 2023, pp. 10381–10399.

29. Mukhopadhyay, S., Qidwai, A., Garimella, A., Ramu, P., Gupta, V., and Roth, D. (2024). Unraveling the truth: Do VLMs really understand charts? A deep dive into consistency and robustness. In Findings of the association for computational linguistics: EMNLP 2024 (Association for Computational Linguistics), pp. 16696–16717. 10.18653/v1/2024.findings-emnlp.973.

30. Huang, J., Chen, X., Mishra, S., Zheng, H.S., Yu, A.W., Song, X., and Zhou, D. (2024). Large language models cannot self-correct reasoning yet. In The twelfth international conference on learning representations (ICLR).

31. Masry, A., Long, D.X., Tan, J.Q., Joty, S., and Hoque, E. (2022). ChartQA: A benchmark for question answering about charts with visual and logical reasoning. In Findings of the association for computational linguistics: ACL 2022 (Association for Computational Linguistics), pp. 2263–2279. 10.18653/v1/2022.findings-acl.177.

32. Kwon, W., Li, Z., Zhuang, S., Sheng, Y., Zheng, L., Yu, C.H., Gonzalez, J.E., Zhang, H., and Stoica, I. (2023). Efficient memory management for large language model serving with PagedAttention. In Proceedings of the 29th symposium on operating systems principles (SOSP ’23) (Association for Computing Machinery), pp. 611–626. 10.1145/3600006.3613165.

33. Coppard, V., Szep, G., Georgieva, Z., Howlett, S.K., Jarvis, L.B., Rainbow, D.B., Suchanek, O., Needham, E.J., Mousa, H.S., Menon, D.K., et al. (2024). FlowAtlas: An interactive tool for high-dimensional immunophenotyping analysis bridging FlowJo with computational tools in Julia. Frontiers in Immunology 15, 1425488. 10.3389/fimmu.2024.1425488.

34. Finak, G., Langweiler, M., Jaimes, M., Malek, M., Taghiyar, J., Korin, Y., Raddassi, K., Devine, L., Obermoser, G., Pekalski, M.L., et al. (2016). Standardizing flow cytometry immunophenotyping analysis from the human ImmunoPhenotyping consortium. Scientific Reports 6, 20686. 10.1038/srep20686.

35. Montante, S., Yokosawa, D., and Brinkman, R. (2025). flowMagic gating benchmark: Automated and manual cell population annotation for flow cytometry data analysis. 10.20383/103.01352.

36. Bjornson-Hooper, Z.B., Fragiadakis, G.K., Spitzer, M.H., Chen, H., Madhireddy, D., Hu, K., Lundsten, K., McIlwain, D.R., and Nolan, G.P. (2022). A comprehensive atlas of immunological differences between humans, mice, and non-human primates. Frontiers in Immunology 13, 867015. 10.3389/fimmu.2022.867015.

