## Supplemental Text, Figures, and Tables for "CytoGate-Bench: an LLM benchmark for cross-panel cell gating in cytometry"

**Document S1. Supplemental methods and notes (sections S1–S10), Figures S1–S10, and Tables S1–S19.**

Section S1 places per-step gating among existing annotation methods, related to the introduction. Section S2 documents cohort provenance and preprocessing, related to Table 1. Section S3 proves that composing per-step decisions recovers leaf-level annotation, related to the methods and Figure 5. Sections S4 and S5 reproduce the production prompt templates verbatim and specify the per-paradigm output handling, both related to the methods. Section S6 consolidates the experimental protocol, metrics, compute, significance testing, and deployment cost, related to Table 2. Section S7 details the visual-modality harnesses, related to Figure 3. Section S8 reports per-scenario robustness, related to Figure 4. Section S9 details the flat-vs.-hierarchical pipelines, related to Figure 5. Section S10 is the reproducibility statement. References are numbered independently of the main reference list, in the order they are first cited in this document.

### S1 Why per-step is the right primitive

The main text adopts per-step cell-type annotation as the task formulation without arguing the choice. This section supplies the biological rationale and surveys flat-annotation alternatives, justifying the choice and substantiating why CytoGate-Bench focuses on the LLM family.

*Two challenges any annotation method must address.*

**(i) Cross-cohort misalignment.** Across cytometry studies, neither the input feature space (the antibody panel) nor the target cell-type space (the populations of interest) aligns out of the box. The combinatorial space of marker subsets, candidate populations, and analytical plans is effectively unbounded, so what is required is not transfer between two specific cohorts but generalization across this space. **(ii) Hierarchical, multi-label assignment.** Cell types form a hierarchy, and human annotation proceeds top-down through it, starting from coarse categories (e.g., T cell) and progressively refining into finer sub-types (e.g., $CD4^+$ T cell → memory $CD4^+$ T cell). Because the hierarchy has multiple legitimate resolutions, the same cell simultaneously carries several labels along the hierarchy. The same multiplicity also arises along different analytical axes. A $CD4^+$ T cell may be labeled *memory* under one marker-pair partition (CCR7 × CD45RA, *differentiation markers*) and *activated* under another (HLA-DR × CD38, *activation markers*), both correct. Any computational method must represent this hierarchical, multi-label structure. A single flat label per cell cannot.

*Existing flat-annotation approaches and their failure modes.*

Existing computational cytometry annotation approaches share a flat output structure (a single label per cell, drawn from a fixed set, with no hierarchy), and three families dominate, none of which addresses both challenges. *Clustering paired with manual labeling* (FlowSOM[1], PhenoGraph[2]) partitions cells unsupervised. A human then assigns a label to each cluster, and per-cell annotations emerge by cluster membership. The workflow is not fully automated and requires fresh manual labeling per cohort. *Supervised classifiers* (DeepCyTOF[3], CellCnn[4], CyAnno[5]) fit on a labeled cohort, binding both the marker set and the label set at training. Neither transfers across panels. *Cytometry foundation models* (cyMAE[6], ImmuneFM[7]) pretrain on large unlabeled cytometry corpora but still bind a fixed label set via a classifier head fine-tuned for cell typing. None of the three bridges the misalignment of challenge (i), and the flat output common to all three cannot represent the hierarchical, multi-label structure required by challenge (ii). The predictions are therefore opaque, with no auditable trace explaining how a cell was assigned its label.

*Per-step annotation is a natural primitive.*

An alternative shape of computation already exists in cytometry, and it builds on knowledge we already have. The cell-type hierarchy specifies, for every population, which parent it derives from and which marker pair distinguishes it from siblings, that is, how the type was defined in the first place. Rather than mapping a cell to a label in a single prediction, per-step annotation walks this hierarchy top-to-bottom as a gating tree. At each step the hierarchy fixes the parent population and the relevant marker pair, and the local decision is which sub-population each cell of the parent belongs to. Manual gating instantiates this in human practice. Its automated members are flowDensity[8] (kernel-density per-step gating) and UNITO[9] (per-step segmentation). We refer to this collection of per-step methods as the *gating-strategy class*. Given a *gating tree* that names, at each step, which marker pair separates which sub-populations, any leaf in the hierarchy, no matter how rare or how unfamiliar, is reachable by composing per-step decisions. Solve one step reliably, and every annotation in the hierarchy becomes accessible without retraining. Per-step annotation is therefore a natural primitive for cell-type annotation.

*Coverage of existing methods against four desiderata.*

Existing methods cover only partial corners of (Flat / Per-step) × (Clustering / Supervised / LLM), and none satisfies all four desiderata simultaneously, namely **(A)** automation/scalability, **(E)** hierarchical-trace explainability, **(F)** freedom from a fixed input/target space, **(Z)** zero-shot operation via prior knowledge. Table S1 positions the landscape. The empty corner is *per-step LLM-based annotation*, the gating-strategy class's per-step composition combined with LLMs' panel-agnostic input, open-vocabulary output, and prior-knowledge-driven inference. This is the corner CytoGate-Bench is designed to evaluate.

Table S1. Coverage of cell-type annotation methods, related to Figure 1. Methods are placed on a task × family grid and scored against four desiderata: (A) automation, (E) hierarchical-trace explainability, (F) input/target-space freedom, (Z) zero-shot via prior knowledge. CytoGate-Bench targets the empty (per-step, LLM) corner.

| Task | Method (family) | A | E | F | Z |
|---|---|---|---|---|---|
| Flat | FlowSOM (Clustering) | × | × | ✓ | × |
| Flat | cyMAE / ImmuneFM (Cytometry FM) | ✓ | × | × | × |
| Flat | Cell-o1 (LLM, scRNA-seq) | ✓ | × | ✓ | ✓ |
| Per-step | flowDensity (Clustering) | × | ✓ | ✓ | × |
| Per-step | UNITO (Supervised) | ✓ | ✓ | × | × |
| Per-step | **CytoGate-Bench target (LLM)** | ✓ | ✓ | ✓ | ✓ |

### S2 Cohort details and preprocessing

#### S2.1 Per-cohort channels and split sizes

Table S2 expands Table 1 of the main text with the channel-level breakdown and per-split sample counts. *Channels* = all channels used for gating (protein markers plus cleanup channels: bead, viability dye, DNA-content, light-scatter, Time, Gaussian event-shape; empty mass-cytometry background channels are excluded). Train / val / test counts follow the donor-stratified 1:1:2 split (seed 42), with adjustments for small cohorts (e.g. FR-FCM-Z74D HC, 5 samples).

Table S2. Per-cohort channels and split sizes (expansion of Table 1 of the main text), related to Table 1. *Markers* = protein channels; *Channels* = all gating channels. MDIPA = Maxpar Direct Immune Profiling Assay. Train / Val / Test = donor-stratified split targeting a 1:1:2 ratio, seed 42; realized counts deviate where donor grouping constrains the assignment (e.g., Lyoplate, FR-FCM-Z74D Tissue) or the cohort is small (FR-FCM-Z74D HC).

| Cohort | Platform | Lineage focus | Markers | Channels | Train | Val | Test |
|---|---|---|---|---|---|---|---|
| Acute2020 | CyTOF (MDIPA) | Whole immune (62-step comprehensive) | 30 | 43 | 6 | 7 | 12 |
| Acute2021 | CyTOF (MDIPA) | Whole immune (62-step comprehensive) | 30 | 43 | 16 | 16 | 33 |
| Vaccine | CyTOF (MDIPA) | Whole immune (62-step comprehensive) | 30 | 43 | 42 | 42 | 85 |
| FR-FCM-Z74D (HC) | Flow | T cell (detailed) | 18 | 18 | 2 | 1 | 2 |
| FR-FCM-Z74D | Flow | T cell (detailed) | 18 | 18 | 6 | 11 | 15 |

| Cohort | Platform | Lineage focus | Markers | Channels | Train | Val | Test |
|---|---|---|---|---|---|---|---|
| (Tissue) | | | | | | | |
| Lyoplate (T cell) | Flow | T cell (basic) | 8 | 10 | 18 | 21 | 24 |
| Lyoplate (Treg) | Flow | Treg | 8 | 10 | 18 | 21 | 24 |
| Lyoplate (B cell) | Flow | B cell | 8 | 10 | 18 | 21 | 24 |
| Lyoplate (DC / Mono / NK) | Flow | DC / Mono / NK | 8 | 10 | 18 | 21 | 24 |
| FRDR COVID-19 | Flow | Myeloid | 10 | 18 | 56 | 56 | 111 |
| Bjornson | CyTOF | Phospho-signaling (8-step) | 40 | 44 | 16 | 16 | 32 |
| *Total (11 cohorts)* | | | | | *216* | *233* | *386* |

#### S2.2 Per-cohort provenance and gate source

The 11 cohorts are re-curated public flow / mass-cytometry studies, each released through one of four manual-gating ecosystems (OMIQ workflow, FlowJo WSP, R `flowWorkspace GatingSet`, or pre-computed label CSV). Table S3 records the source repository, gating tool, and number of original FCS samples per cohort, together with the per-cohort exclusions imposed during harmonization. IRB approval and informed consent for each cohort are documented in the source publication cited in the table.

Table S3. Per-cohort source provenance, related to Table 1. *Tool* = source format; *Released / Used* = FCS files released vs. retained after harmonization. *Used* counts samples surviving harmonization; the per-cohort totals in Table 1 of the main text reflect the donor-stratified 1:1:2 split-resident count, which is two samples lower in aggregate (Acute2020: 25 vs. 26; FR-FCM-Z74D (HC): 5 vs. 6) due to split assignment.

| Cohort | Source | Tool | Released | Used | Exclusions |
|---|---|---|---|---|---|
| Acute2020 | Acute COVID[6] | OMIQ workflow | 27 | 26 | 1 sample without GT export |
| Acute2021 | I-SPY[6] | OMIQ workflow | 65 | 65 | none |
| Vaccine | ImmuneHealth COVID-vac[6] | OMIQ workflow | 177 | 169 | 8 samples without GT export |
| FR-FCM-Z74D (HC) | FlowRepository FR-FCM-Z74D[10] | FlowJo WSP → R | 6 | 6 | none |
| FR-FCM-Z74D (Tissue) | FlowRepository FR-FCM-Z74D[10] | FlowJo WSP → R | 32 | 32 | none |
| Lyoplate (T cell) | HIPC Lyoplate[11] | GatingSet → R | 63 | 63 | none |
| Lyoplate (Treg) | HIPC Lyoplate[11] | GatingSet → R | 63 | 63 | none |
| Lyoplate (B cell) | HIPC Lyoplate[11] | GatingSet → R | 63 | 63 | none |
| Lyoplate (DC/Mono/NK) | HIPC Lyoplate[11] | GatingSet → R | 63 | 63 | none |
| FRDR COVID-19 (Panel 8) | FRDR COVID-19[12], BD LSRFortessa X20 | FlowJo WSP → R | 237 | 223 | 11 filename-mismatched, 3 non-standard tree |
| Bjornson | Nolan-lab cross-species phospho-CyTOF[13] | OMIQ workflow | 64 | 64 | none |
| *Total* | | | *860* | *837* | |

*Licensing and data release.*

The source cohorts carry heterogeneous terms. The three CyTOF cohorts of Kim et al.[6] (Acute2020, Acute2021, Vaccine) are released under CC-BY 4.0, and FRDR COVID-19[12] under CC0 1.0. FR-FCM-Z74D[10] and the Bjornson cohort[13] are public on FlowRepository under ISAC open data-sharing terms. No per-dataset Creative Commons tag is attached, and reuse is governed by attribution to the source publication. The four HIPC Lyoplate cohorts[11] are *not* openly licensed. They are distributed through ImmPort/ImmuneSpace under the ImmPort user agreement, which requires free registration, prohibits re-identification, and permits redistribution only under terms commensurate with the agreement. Our release therefore bundles the harmonized arcsinh cell-by-marker matrices for all 11 cohorts in a single Dataverse dataset whose custom terms state each cohort's license, with the openly licensed cohorts under their original licenses and the four Lyoplate cohorts under terms that mirror the ImmPort agreement (no re-identification, attribution to ImmPort and the source study accession, and redistribution only under the same terms). We additionally release the extraction scripts that rebuild the per-step Lyoplate data from the source FCS files for users who prefer to obtain them directly from ImmuneSpace under that agreement. Our added layer (the per-step annotations, gating trees, and evaluation splits) is released under CC-BY 4.0 and the accompanying code under the MIT License, while each underlying cohort retains its original license as listed above.

#### S2.3 Bio-marker transformation: arcsinh

All antibody / fluorophore-conjugated channels (*protein*), together with viability dye, DNA intercalator, normalization-bead, Gaussian event-shape, and background isotope channels, are transformed by $\mathrm{arcsinh}(x/\mathrm{cofactor})$ before storage. The cofactor is platform-specific, $5$ for mass cytometry (CyTOF) and $150$ for conventional flow cytometry, following the standard cytometry conventions for each modality. Light-scatter (`FSC-*`, `SSC-*`) and technical (`Time`, `Event_length`) channels are not arcsinh-transformed. Instead, they are clipped to the per-sample $[1,99]$ percentile range and min-max scaled to $[0,10]$. All channels are stored as `float16` parquet columns with zstd compression, identical across cohorts. The cofactor used for each cohort is recorded per cohort alongside the channel-type assignment, and all gate-polygon vertex coordinates are stored in the same arcsinh space, so no further transform is needed at evaluation time to align gates with cells.

#### S2.4 Compensation

Mass cytometry uses isotopic mass tags rather than fluorophores and therefore has no spectral spillover. The CyTOF cohorts (Acute2020/2021, Vaccine, Bjornson) contribute compensation-free expression. The conventional-flow cohorts inherit per-cohort compensation matrices from their source workspaces. Cohorts harvested from FlowJo workspaces (FRDR COVID-19, FR-FCM-Z74D) are extracted with `flowWorkspace` in R, which applies the workspace-stored compensation matrix to the linear-scale FCS values before our arcsinh transform. Lyoplate enters our pipeline through legacy `GatingSetList` objects whose stored expression is already compensated. The Acute2020/2021 and Vaccine OMIQ workflows include manual cleanup gates upstream of cell-type annotation (bead removal, debris, doublet, viability), which are recorded as the first three to seven steps of each cohort's gating tree rather than discarded. Per-step accuracy thus rewards methods that handle cleanup gates correctly.

#### S2.5 Gate extraction

Gate definitions are harvested from each cohort's source format and serialized to a uniform per-sample JSON keyed by step index, with each step listing one or more gates as a polygon (or rectangle expanded to four vertices) in the same arcsinh-transformed marker space as the parquet expression columns. Three extraction paths are used:

- **OMIQ API** (Acute2020/2021, Vaccine, Bjornson). Per-cohort workflow + `GatingTask` are queried through the OMIQ REST API, and `filterContainers` provide the polygon vertices and the per-file overrides (`perFileFilters`) for sample-specific gate adjustments. Mass-cytometry channel names are mapped from gating-tree markers via the isotope suffix (`145Nd_CD4` → `Nd145Di`). Flow channels are mapped through a per-cohort `channel_map` aligning instrument channel names with marker columns. OMIQ `CompoundFilter` virtual gates defined as Boolean NOT compositions (e.g., `nnCD4` = CD4 NOT CD4Naive) are not represented as polygons in OMIQ and are skipped at extraction. Their child gates remain reachable through the parent population mask.

- **FlowJo WSP via R** (FRDR COVID-19, FR-FCM-Z74D). `flowWorkspace::parseWorkspace` parses each WSP, and `gh_pop_get_data` extracts compensated expression together with one Boolean column per gate name. A per-cohort `GATE_TO_CATEGORY` dictionary maps the FlowJo gate names (which are workspace-author conventions, not standardized) onto the gating-tree categories used by the benchmark.

- **Legacy GatingSetList** (Lyoplate). The HIPC-released `GatingSetList` objects are loaded with `flowWorkspace`, sub-sampled to one panel per cohort (B-cell, T-cell, DC, Treg), and exported with `gh_pop_get_data`. Live-singlet pre-filtering (debris, doublets, dead) is applied as the workspace specifies and is not exposed as an annotation step in our gating tree. The remaining Boolean gate columns are mapped onto the gating-tree categories per panel.

For all three paths, the extracted per-sample gates are validated against the gating tree by checking that every step's parent expression resolves to a well-defined parent mask and that polygon vertex coordinates lie within the per-sample $[1,99]$ percentile range of their channel. Samples whose gates fail to extract, for example through a missing ground-truth export or a filename mismatch, or that contain a degenerate gate are excluded (Table S3, "Exclusions" column). A step whose parent population is empty in an otherwise valid sample is not an exclusion. The sample is retained and that step contributes no evaluation instance.

#### S2.6 Per-cell gate annotation

The gating tree defines one annotation step per (parent population, marker pair) pair. For each cell that lies in a step's parent mask, the cell is checked against every child gate of that step in order, and the first matching child becomes the cell's per-step label. Cells in the parent that match no listed child are left as missing (`NaN`) for that step and propagate to *Unassigned* downstream, as specified in the task definition above. We follow a strict *gate-positive only* convention. Only categories with an explicit gate in the source data appear in the gating tree, and we do not invent derived negative categories (*e.g.*, *Dead*, *Doublets*, *Other*, *Non-Treg*) for unmatched cells. The single exception is when a negative subset is itself used as a downstream parent (e.g., `non_Tfh` parents step 6 in some panels), in which case the source must already provide an explicit gate for it. This discipline ensures every non-NaN label in our parquet outputs is grounded in the original source-recorded gating, never inferred from omission, and that per-step accuracy is computed against the same labels the original analyst would have inspected.

### S3 Per-step composition recovers flat classification

This section formalizes the claim that per-step gating is no less expressive than flat classification on the gating tree. Every flat-classification policy admissible on the tree is realizable by composing per-step rules, and a per-step solver with bounded per-node error yields a flat classifier whose error is bounded linearly by tree depth.

*Notation.*

Let $H=(V,E)$ be a finite rooted tree with root $v_{\text{root}}$, internal node set $V_I := V \backslash L(H)$, and leaf set $L(H)$ naming the terminal cell types. The depth of $H$ is $D := \max_{c\in\mathcal{X}} |\pi^*(c)| - 1$, where $\pi^*(c) = (v_0, v_1, \ldots, v_{K(c)})$ denotes the unique ground-truth root-to-leaf path of cell $c \in \mathcal{X}$ ($v_0 = v_{\text{root}}$, $v_{K(c)} \in L(H)$, and $K(c) \le D$ for all $c$). We write $\psi^*_{\text{leaf}}(c) := v_{K(c)} \in L(H)$ for the ground-truth leaf of $c$. Each internal node $v \in V_I$ specifies a per-step task: a parent population $P_v \subseteq \mathcal{X}$ (the cells of any sample that reach $v$ under the ground-truth path), an ordered marker pair $(x_v, y_v)$, and a candidate child set $C_v$ equal to the set of children of $v$ in $H$. We extend the codomain with a sentinel $\perp$ standing for *Unassigned*. A *per-step solver* is a family

$$\psi = \{\psi_v\}_{v \in V_I},$$
$$\psi_v : \mathcal{X} \to C_v \cup \{\perp\}.$$

We define each $\psi_v$ on all of $\mathcal{X}$ rather than only on $P_v$. A flat classifier $g$ may route a cell $c$ through nodes that lie off its ground-truth path $\pi^*(c)$, so the recursion below must be able to query $\psi_v$ at such cells. On $P_v$, the ground-truth solver $\psi^* = \{\psi_v^*\}_{v \in V_I}$ satisfies $\psi_v^*(c) = v_{k+1}$ whenever $v = v_k$ lies on $\pi^*(c)$. Equivalently, $\psi_v^*(c) \in C_v$ records the expert child of $v$ for $c$. (Off $P_v$, $\psi_v^*$ is irrelevant to the statements below, and we may take $\psi_v^*(c) = \perp$ by convention.)

The *composed flat classifier* $F_\psi : \mathcal{X} \to L(H) \cup \{\perp\}$ is defined recursively by descending $H$ from $v_{\text{root}}$ guided by $\psi$. Concretely, $F_\psi(c) = \mathrm{descend}(v_{\text{root}}, c)$ where

$$\mathrm{descend}(v, c) := \begin{cases} v, & v \in L(H), \\ \perp, & \psi_v(c) = \perp, \\ \mathrm{descend}(\psi_v(c), c), & \text{else.} \end{cases}$$

Each recursive call moves to a strictly deeper node of the finite tree $H$, so the recursion terminates after at most $D$ calls.

**Proposition S1 (Composition along a target path).**

Let $c \in \mathcal{X}$ and let $\pi = (v_0, v_1, \dots, v_K)$ be any root-to-leaf path in $H$, i.e., $v_0 = v_{\text{root}}$, $v_K \in L(H)$, and $(v_k, v_{k+1}) \in E$ for $0 \le k < K$. If

$$\psi_{v_k}(c) = v_{k+1} \qquad (0 \le k < K),$$

then $F_\psi(c) = v_K$.

*Proof.* For $k \in \{0, 1, \dots, K\}$ let $S(k)$ denote the statement *"the recursion* $\mathrm{descend}$ *invoked on* $(v_{\text{root}}, c)$ *reaches* $v_k$ *after exactly* $k$ *recursive calls."*

*Base case* ($k = 0$). By definition $F_\psi(c) = \mathrm{descend}(v_{\text{root}}, c) = \mathrm{descend}(v_0, c)$, so $S(0)$ holds vacuously.

*Inductive step.* Assume $S(k)$ for some $k < K$. Then $\mathrm{descend}$ is currently invoked on $(v_k, c)$, and $v_k \in V_I$ since $k < K$ and $\pi$ is a root-to-leaf path. The hypothesis gives $\psi_{v_k}(c) = v_{k+1}$, and $(v_k, v_{k+1}) \in E$ ensures $v_{k+1} \in C_{v_k}$. In particular $\psi_{v_k}(c) \neq \perp$, so the third branch of $\mathrm{descend}$ fires and the recursion descends to $\mathrm{descend}(v_{k+1}, c)$. This establishes $S(k+1)$.

By induction, $S(K)$ holds. The recursion reaches $(v_K, c)$, and since $v_K \in L(H)$ the first branch returns $v_K$. Therefore $F_\psi(c) = v_K$.

Proposition S1 is stated abstractly over an arbitrary root-to-leaf path $\pi$. The two corollaries below instantiate it with two different choices of $\pi$. Setting $\pi = \pi^*(c)$ (the ground-truth path) and $v_{k+1} = \psi_{v_k}^*(c)$ yields the agreement-with-ground-truth statement used for error compounding. Setting $\pi = \pi_g(c)$ (the path induced by a flat classifier $g$, defined below) yields representational completeness.

**Corollary S2 (Error compounding).**

Fix a probability space carrying the randomness of $\psi$ (e.g., training randomness, stochastic decision rules). The cell $c$ is fixed. Suppose every per-step rule errs with probability at most $\varepsilon$ on its parent population:

$$\Pr[\psi_v(c) \neq \psi_v^*(c)] \ \le \ \varepsilon$$
for every $v \in V_I$ and $c \in P_v$.

Then for every $c \in \mathcal{X}$ with ground-truth path of length $K = K(c)$,

$$\Pr\left[F_\psi(c) \neq \psi^*_{\text{leaf}}(c)\right] \;\leq\; K\varepsilon \;\leq\; D\varepsilon.$$

Averaging over an independent population draw $c \sim \mathcal{D}$ on the product probability space (Fubini), the same bound holds in expectation with $K$ replaced by $\mathbb{E}_{c\sim\mathcal{D}}[K(c)] \leq D$.

*Proof.* Write $\pi^*(c) = (v_0, v_1, \dots, v_K)$, and let $E_k := \{\psi_{v_k}(c) \neq \psi^*_{v_k}(c)\}$ for $k = 0,1,\dots,K-1$. Since $c \in P_{v_k}$ for each such $k$ (the $v_k$ lie on $\pi^*(c)$), the per-node hypothesis yields $\Pr[E_k] \leq \varepsilon$. If $K = 0$ then $\pi^*(c) = (v_0)$ with $v_0 \in L(H)$, so $\mathrm{descend}(v_{\text{root}}, c)$ returns $v_0 = \psi^*_{\text{leaf}}(c)$ immediately and the claim is trivial. Assume $K \geq 1$.

On the complement $\bigcap_{k=0}^{K-1} E_k^c$, we have $\psi_{v_k}(c) = \psi^*_{v_k}(c) = v_{k+1}$ for all $0 \leq k < K$, so Proposition S1 applied with $\pi = \pi^*(c)$ gives $F_\psi(c) = v_K = \psi^*_{\text{leaf}}(c)$. By contrapositive,

$$\{F_\psi(c) \neq \psi^*_{\text{leaf}}(c)\} \;\subseteq\; \bigcup_{k=0}^{K-1} E_k,$$

and the union bound gives

$$\begin{aligned}\Pr\left[F_\psi(c) \neq \psi^*_{\text{leaf}}(c)\right] &\leq \sum_{k=0}^{K-1} \Pr\,[E_k] \\ &\leq K\varepsilon \;\leq\; D\varepsilon.\end{aligned}$$

For the population statement, integrate the pointwise bound $\Pr[F_\psi(c) \neq \psi^*_{\text{leaf}}(c) \mid c] \leq K(c)\varepsilon$ against $\mathcal{D}$ (Fubini on the product space of training randomness and $c \sim \mathcal{D}$) and apply $K(c) \leq D$.

**Remark S1.** The union bound is loose. Once descend leaves the ground-truth path at some step $k^* < K$, the subsequent queries $\psi_{v_k}(c)$ for $k > k^*$ are never made by the recursion, so $E_{k^*+1}, \dots, E_{K-1}$ cannot contribute to the failure event. A tighter bound replaces $\bigcup_k E_k$ by the event that the *first* disagreement occurs on the path. The linear-in-$D$ statement above suffices for our purposes.

**Corollary S3 (Representational completeness).**

For every flat classifier $g: \mathcal{X} \to L(H)$, there exists a per-step solver $\psi^g$ such that $F_{\psi^g} \equiv g$.

*Proof.* For each $c \in \mathcal{X}$ let $\pi_g(c) = (u_0, u_1, \dots, u_{K_g(c)})$ denote the unique root-to-leaf path in $H$ with $u_0 = v_{\text{root}}$ and $u_{K_g(c)} = g(c)$. Uniqueness follows from $H$ being a tree (each leaf has a unique ancestor chain). Define, for every $v \in V_I$ and every $c \in \mathcal{X}$,

$$\psi^g_v(c) \;:=\; \begin{cases} u_{k+1}, & \text{if } v = u_k,\ k < K_g(c), \\ \perp, & \text{otherwise.} \end{cases}$$

Because each $\psi^g_v$ is defined on all of $\mathcal{X}$, the recursion descend may freely query $\psi^g_v$ even at cells $c$ whose ground-truth path does not pass through $v$.

If $K_g(c) = 0$ then $g(c) = u_0 = v_{\text{root}} \in L(H)$, and $\mathrm{descend}(v_{\text{root}}, c)$ returns $v_{\text{root}} = g(c)$ at the first branch. Otherwise $K_g(c) \geq 1$, and by construction $\psi^g_{u_k}(c) = u_{k+1}$ for every $0 \leq k < K_g(c)$. Applying Proposition S1 with target path $\pi = \pi_g(c)$ yields $F_{\psi^g}(c) = u_{K_g(c)} = g(c)$.

*Remark.*

Proposition S1 and Corollaries S2–S3 together justify per-step as the productive primitive for evaluation. *(i) Composition* (Proposition S1) with *error compounding* (Corollary S2) shows that solving the local primitive carries flat-classification accuracy along with it, with depth $D$ as the only multiplicative cost. The gating trees in CytoGate-Bench are of moderate depth (section S2), so per-step accuracy translates directly into flat accuracy. *(ii) Representational completeness* (Corollary S3) shows per-step is no less expressive than flat. *(iii)* But per-step is strictly more *local*. Each $\psi_v$ depends only on the local task

$(P_v, x_v, y_v, C_v)$, so a method that solves per-step needs no global view of the panel or the leaf vocabulary, which is the cross-cohort, panel-agnostic property that motivates the benchmark. We therefore evaluate per-step accuracy as the headline metric. Flat-classification accuracy follows by composition, and the empirical translation is verified in our flat-vs.-hierarchical comparison.

### S4 Prompt templates for the LLM-class methods

This section reproduces the production prompt templates for the two LLM-class methods (LLM-C2S, LLM-Gate). Both use a shared user-prompt skeleton with four blocks, namely `[Context]` (modality, parent path, focal marker pair, parent cell count), `[Categories]` (the candidate-child set $C_v \cup \{\text{Unassigned}\}$), `[Description]` (the curated marker priors $D_v$), and a method-specific data block, followed by a JSON output schema. The examples below are real prompt renderings emitted by the benchmark pipeline (Acute2020 / 994570_Normalized, step 18: CD38 × CD14 monocyte split). Numeric histograms are abbreviated with "..." for compactness. The full templates are in the released code.

#### S4.1 LLM-Gate prompt

The data block is a per-axis 1-D histogram for each of the focal marker pair, with pre-extracted density peaks and valleys (P$i$ / V$i$) annotated as positional landmarks on the unicode bar plot. The model must return one axis-aligned rectangle per named category in JSON. *Unassigned* is the residual and is never gated explicitly.

**LLM-Gate user prompt**

```
[Context]
Modality: CyTOF (arcsinh cofactor: 5)
Path: Step17_CD11c|CD14 == TotalMonocyte -> [CD38 vs CD14]
Step: 18
X-axis marker: CD38
Y-axis marker: CD14
Parent cells: 14,622

[Categories]
ClassicalMono, TransitionalMono, NonclassicalMono, Unassigned

[Description]
ClassicalMono: classical monocytes -- main dense blob upper-right
  (CD38 high, CD14 high).
TransitionalMono: intermediate monocytes -- mid-cluster
  (CD38 mid, CD14 mid).
NonclassicalMono: patrolling monocytes -- lower-left
  (CD38 low, CD14 low-to-dim).
Unassigned: cells whose (x, y) position falls outside every named
  category's gate at this step.

[Axis distribution]
### CD38 (x-axis), range [0.00, 6.52], n=14622 parent cells
... unicode bar plot (40 bins) ...
P1: x=4.48, frac=0.46

### CD14 (y-axis), range [0.00, 6.60], n=14622 parent cells
... unicode bar plot (40 bins) ...
P1: x=0.16, frac=0.04
```

```
P2: x=4.06, frac=0.29
V1: x=1.16 (between P1<->P2)

## Output Format
Return a JSON object keyed by category name. For each named category,
output either a single rectangle dict (default) or a list of rectangle
dicts (multi-mode populations); coordinates are in the same numeric
units as the axis distribution above. Cells inside multiple
rectangles are resolved at evaluation time (smallest-area wins).

{
  "ClassicalMono":     {"x_min": ..., "x_max": ...,
                         "y_min": ..., "y_max": ...,
                         "rationale": "..."},
  "TransitionalMono":  {...},
  "NonclassicalMono":  {...}
}
```

The accompanying system prompt fixes three guardrails. (i) The model first estimates the expected proportion of each category from biology before reading the histogram, preventing rare populations from overrunning dominant modes. (ii) Every boundary is anchored to a histogram landmark (P$i$ or V$i$) rather than to axis extremes. (iii) A category may be declared *absent* (`{"absent": true, "rationale": ...}`) when biology and the axis distribution agree it has no mass. This is the depleted-population path that drives the robustness behavior. The system prompt (shared with the zero-shot panel evaluation) is reproduced below in abridged form.

**LLM-Gate system prompt (abridged)**

```
You are an expert in single-cell CyTOF and Flow Cytometry gating.
For each step you see the parent population's distribution on a 2D
(x, y) plane; define the gates that separate the listed categories
on that plane.

## What you produce
For every named category, output an axis-aligned rectangle
{x_min, x_max, y_min, y_max} in the same numeric units as the axis
distribution shown -- do not invent a new scale.

## Step 1 -- Estimate expected proportion from biology
Before reading the histogram, estimate each category's expected
proportion within this parent (rare <10%, common 10-50%, dominant
>50%) from the description, tips, and immunology prior. Use it as a
sanity check: a rare category must not cover the dominant mode.

## Unassigned holds the residual
A cell is Unassigned when it falls outside every named gate. Do NOT
output a gate for Unassigned.

## Declaring a category absent
Emit {"absent": true, "rationale": ...} ONLY when (1) biology allows
the category to be missing, (2) the histogram shows no mode there,
and (3) the expected proportion is ~0%. If unsure, emit the best
```

```
(even tight) gate instead. (...)

## Positive vs. negative
Positive/negative is relative to the parent's distribution, not a
fixed threshold. Pitfall: in pre-gated panels a "negative" cluster
can sit at moderate-to-high absolute values; anchor boundaries to
the reported peak (Pi) / valley (Vi) landmarks, never to axis-zero
or axis-max. (...)

## Non-overlap
Gates must not overlap; each cell falls in at most one named gate.

## Multi-rectangle gates
Default to ONE rectangle per category; emit a list only when the
category forms spatially-disconnected clusters that one rectangle
cannot cover without sweeping in a different population. (...)

## Output format
A single JSON object keyed by category name; each value is a
rectangle dict, a list of rectangle dicts, or an absent dict, each
with a one-sentence rationale.
```

**S4.2 LLM-C2S prompt**

LLM-C2S clusters the parent on the $(x_v, y_v)$ plane (flowDensity or FlowSOM, reported in the variant truncated to the top highly variable protein (HVP) markers) and serializes each cluster's representative cell as a rank-encoded "cell sentence". The same `[Context]` / `[Categories]` / `[Description]` header is reused. The data block lists one cell per cluster.

**LLM-C2S user prompt**

```
[Context]
Modality: CyTOF (arcsinh cofactor: 5)
Path: Step17_CD11c|CD14 == TotalMonocyte -> [CD38 vs CD14]
Step: 18
Parent cells: 14,622
Representative cells in this batch: 16

[Categories]
ClassicalMono, TransitionalMono, NonclassicalMono, Unassigned

[Description]
... same description block as LLM-Gate ...

[Top HVP markers (excl. axis)]
CD45RA, CCR6, HLA-DR, CD11c, CD11b, CD16, CD33, CD64, CD86, CD123

## cell_1 (n=4,356, 29.8%)
[axis] CD38(4.61), CD14(4.78)
[hvp]  HLA-DR(3.82) > CD11b(3.57) > CD11c(3.34) > CD64(2.91)
       > CD33(2.41) > CD86(2.11) > CCR6(0.40) > CD45RA(0.20)
       > CD16(0.18) > CD123(0.12)
```

```
## cell_2 (n=2,189, 15.0%)
[axis] CD38(2.14), CD14(2.71)
[hvp]  CD11c(3.10) > HLA-DR(2.84) > CD11b(2.41) > CD33(2.05)
       > ...

## cell_3 (n=1,512, 10.3%)
[axis] CD38(0.71), CD14(0.42)
[hvp]  CD16(3.21) > CD11c(2.84) > HLA-DR(2.31) > CCR6(1.42)
       > ...

... 13 more cluster representatives (cell_4 ... cell_16) ...

## Output Format
Return one line per cluster: "[cluster_id] <label>", where label is
one of {ClassicalMono, TransitionalMono, NonclassicalMono,
Unassigned}. The full parent population is recovered by propagating
each cluster's label to every cell that was assigned to it during
the upstream grouping step.
```

The system prompt is reproduced below in abridged form.

**LLM-C2S system prompt (abridged)**

```
You are an expert in single-cell CyTOF and Flow Cytometry gating.
Classify each anchor cell into one of the listed categories by
drawing the gate that defines that category and checking whether the
cell falls inside it.

## Cell sentence
Each token is Marker(arcsinh_value), ordered by descending
expression. Each "## cell_N" header carries (n=N_cells, X.X%) -- the
cluster's parent-cell count and share. Size is a confidence prior on
stability, NOT a category prior.

## Categories define what to classify
Read [Description] with [Categories]; the name is a label, not the
specification. Match the cell's expression profile to the targeted
phenotype, not the literal name and not by elimination.

## Unassigned holds the residual
A cell is Unassigned when it falls outside every category's gate --
not merely when you are uncertain which named category fits.

## Positive vs. negative
Relative to the parent's distribution at this step, not a fixed
absolute threshold.

## Marker reference
arcsinh-transformed protein / lineage / bead / DNA / gaussian /
live-dead channels (relative ordering meaningful); technical
(Time, Event_length) and scatter (FSC, SSC) channels are linearly
```

```
rescaled to 0-10.

## Consistency check
A single high marker is weak evidence; confirm each call against the
expected co-expression of the cell sentence's other top markers.
```

The cell sentence is HVP-truncated for the reported runs (`hvp10`, the top-10 panel-wide HVP markers per cluster, sorted high→low after the axis markers).

### S5 Paradigm-specific output handling

Methods whose natural output does not project onto $C_v \cup \{\text{Unassigned}\}$ are coerced as follows. (i) LLM-class JSON outputs are validated as axis-aligned rectangles. On parse failure, every cell of the step is predicted *Unassigned*. (ii) Cells outside every returned rectangle are routed to *Unassigned* per the task-definition rule, and a cell inside multiple overlapping rectangles takes the label of the smallest-area rectangle containing it. For clustering paradigms, cluster IDs are mapped to candidate labels by per-cluster LLM labeling.

*Output cardinality and paradigm contrast.*

The two LLM-class paradigms commit to outputs at different scales, which has direct consequences for both cost and the propagation rule of (ii) above. **LLM-C2S** consumes a multi-marker phenotype per pre-formed cluster and emits one label per cluster ($K$ labels per step, $K \in [2,20]$ depending on the clustering primitive). Per-cell predictions follow by cluster membership, so any error at the upstream grouping stage (for example, a single cluster straddling two candidate populations) upper-bounds the achievable per-cell accuracy regardless of the downstream LLM call. **LLM-Gate** consumes the marker-pair distribution as two per-axis 1-D histograms in the same arcsinh units as the input expression matrix, and emits exactly one axis-aligned rectangle $B_c \subseteq \mathbb{R}^2$ per candidate. A $|P_{v,s}|$-cell annotation thus reduces to an $\mathcal{O}(|C_v|)$-rectangle output regardless of population size, and the containment rule of (ii) turns the rectangles into per-cell labels deterministically (cells contained in no rectangle default to *Unassigned*, and cells in multiple overlapping rectangles take the smallest-area rectangle's label). The two paradigms therefore diverge in both *what the LLM sees* (a per-cluster phenotype profile vs. a marker-pair distribution) and *what it commits to* (a per-cluster label vs. a per-category geometric rule).

### S6 Experimental setup and reproducibility

This section consolidates the protocol summarized in the main text, namely the method roster, the train/val/test regime, the metric definitions, and inference settings.

#### S6.1 Method roster and training regime

Table S4 lists the three method groups compared throughout the main text, together with the input view each method consumes per step, whether it is fit on cohort labels or evaluated zero-shot, and the backbones used. The 1:1:2 train/val/test partition is donor-stratified with seed 42 (section S2.1, Table S2). The *trained baselines* (UNITO, cyMAE) are fit per cohort on train+val and evaluated on test, while *LLM-class methods* (LLM-C2S, LLM-Gate, VLM-Gate, Agent-Gate) see no labels and operate purely on the per-step inputs $(P_{v,s}, x_v, y_v, C_v, D_v)$ formalized in the methods (main text).

Table S4. Method roster, related to Table 2. *Input view*: what the method consumes at one annotation step. *Regime*: *fit* = trained on cohort train+val labels; *zero-shot* = no label access. *Backbones*: rows shown in Table 2 of the main text.

| Group | Method | Input view | Regime | Backbone(s) |
|---|---|---|---|---|
| Trained | UNITO[9] | 2D pair $(x_v, y_v)$ | fit per-cohort | — |
| | cyMAE[6] | all markers | fit per- | — |

| Group | Method | Input view | Regime | Backbone(s) |
|---|---|---|---|---|
| | | | cohort | |
| LLM-C2S | density-grid + LLM[8] | per-cluster phenotype (flowDensity) | zero-shot | Cell-o1 7B, Qwen3.5 4B/27B, Qwen3.6 27B, Gemma 4 26B-A4B/31B, GPT-5.4 |
| | clustering + LLM[1] | per-cluster phenotype (FlowSOM) | zero-shot | (same six backbones) |
| LLM-Gate | LLM-Gate | two 1-D histograms | zero-shot | (same six backbones) |
| | VLM-Gate | + 2-D scatter image | zero-shot | GPT-5.4 only |
| | Agent-Gate | + scatter image + render tool | zero-shot | GPT-5.4 only |

Cell-o1 (7B)[14] is a Qwen2.5-7B fine-tuned on transcriptomic cell-type reasoning. It is included as a domain-tuned reference and is the only fine-tuned LLM in the roster.

#### S6.2 Aggregation and metrics

Per-cohort scores are computed by averaging per-step F1 macro, balanced accuracy (BA), and Hull IoU over all (sample, step) instances in the cohort's test split. The *Mean* column of Table 2 of the main text averages these per-cohort scores with equal weight across the 11 cohorts (cohort, not task, is the aggregation unit, so panel diversity is not dominated by the sample-heavy CyTOF cohorts). F1 macro is the unweighted mean of per-candidate F1 over $C_v \cup \{\text{Unassigned}\}$. BA is the mean of per-candidate recall over the candidates of that set with nonzero ground-truth support in the instance, following the scikit-learn convention. Hull IoU is the intersection-over-union between the convex hulls of predicted and ground-truth cells assigned to the same candidate in the $(x_v, y_v)$ arcsinh plane, with each point set trimmed to its per-axis $[1,99]$ percentile range before the hull is computed to reduce outlier sensitivity.

#### S6.3 Inference and generation settings

All LLM backbones are run zero-shot, with a maximum completion budget of $32768$ tokens and a single generation attempt per step. On a parse failure every cell of the step is predicted *Unassigned* (section S5). Open-weight reasoning backbones (Qwen3.5/3.6, Gemma 4) additionally receive a thinking-token budget of $8096$ (`tb8096` variant). Decoding follows each backbone's recommended setting. The closed-weight GPT-5.4 is greedy (temperature 0), whereas the open-weight reasoning backbones sample at temperature 1.0, top-$p$ 0.95, and top-$k$ 20 (Qwen) / 64 (Gemma), since greedy decoding degrades these reasoning modes. The open-weight rows are therefore stochastic. We verify their seed-stability in Table S5, where the cohort-mean F1 macro shifts by at most $\pm 0.01$ across 10 seeds. These generation settings are shared by every LLM-class row of Table 2 of the main text, by the visual-modality variants VLM-Gate and Agent-Gate (tool-loop schedule in section S7), and by the flat-vs. hierarchical comparison (section S9), whose only experiment-specific knob is the FlowSOM clustering granularity ($K$ set to the leaf count per depth, $16/38/83$). Full LLM-C2S and LLM-Gate prompts, including the JSON output schema validated against, are in section S4.2, S4.1. Per-paradigm output cardinality and the per-cell propagation rule (rectangle containment, cluster membership, *Unassigned* residual) are summarized in section S5.

Table S5. Seed stability of the open-weight LLM-Gate rows, related to Table 2. Cohort-mean F1 macro / BA / Hull IoU as mean±std over 10 random generation seeds (seeds 1–7, 42, 123, 456), with $\max_c \sigma_{\text{F1}}$ the largest per-cohort standard deviation of F1 macro. The LLM-Gate values in Table 2 of the main text are these seed means. Despite stochastic sampling (section S6.3), the aggregate metrics move by $\leq \pm 0.01$ across seeds. The closed-weight GPT-5.4 row is greedy and thus seedless.

| Backbone | F1 macro | BA | Hull IoU | $\max_c \sigma_{\text{F1}}$ |
|---|---|---|---|---|
| Qwen3.5-4B | $0.624 \pm 0.010$ | $0.710 \pm 0.008$ | $0.561 \pm 0.012$ | $0.038$ |
| Qwen3.5-27B | $0.656 \pm 0.003$ | $0.738 \pm 0.003$ | $0.611 \pm 0.003$ | $0.028$ |
| Qwen3.6-27B | $0.657 \pm 0.002$ | $0.743 \pm 0.002$ | $0.612 \pm 0.003$ | $0.015$ |
| Gemma 4 26B A4B | $0.640 \pm 0.002$ | $0.731 \pm 0.002$ | $0.588 \pm 0.003$ | $0.013$ |

| Backbone | F1 macro | BA | Hull IoU | $\max_c \sigma_{\mathrm{F1}}$ |
|---|---|---|---|---|
| Gemma 4 31B | $0.656 \pm 0.001$ | $0.741 \pm 0.001$ | $0.614 \pm 0.001$ | $0.007$ |

#### S6.4 Compute and hardware

All open-weight LLMs are served with vLLM[15] on a single node of eight NVIDIA H200 GPUs (141 GB each). The closed-weight model (GPT-5.4) is accessed through its vendor API and uses no local accelerators. The generation settings of section S6.3 are held fixed across this hardware. As a representative cost, a full zero-shot evaluation sweep with Qwen3.6-27B completes in roughly 2 hours of wall-clock on this node (≈16 GPU-hours). The trained baselines are fit and evaluated on separate nodes, UNITO on $4 \times$ H200 in about 2 hours total (≈8 GPU-hours), and cyMAE on $4 \times$ A100 in about 6 hours total (≈24 GPU-hours).

#### S6.5 Cross-cohort significance testing

The *Mean* column of Table 2 of the main text averages 11 per-cohort scores, so the cohort is the natural unit for a paired test. We compare each LLM-Gate backbone against UNITO, the strongest trained baseline, with a two-sided Wilcoxon signed-rank test on the 11 paired cohort means ($n = 11$). Table S6 reports the result. No metric separates the two method classes at $\alpha = 0.05$. With $n = 11$ the test's power is limited, so these are failures to detect a difference and not evidence of equivalence. We therefore describe the comparison as an unresolved difference on the chosen metrics rather than as a win for either side.

Table S6. Wilcoxon signed-rank tests over the 11 cohort means, each LLM-Gate backbone against UNITO, related to Table 2. $\Delta =$ UNITO − LLM, so a positive Δ favors UNITO. Two-sided $p$; n.s. = not significant at $\alpha = 0.05$.

| Backbone | Metric | UNITO | LLM | Δ | $p$ | Verdict |
|---|---|---|---|---|---|---|
| GPT-5.4 | F1 macro | 0.690 | 0.676 | +0.014 | 0.46 | n.s. |
| | BA | 0.757 | 0.768 | −0.011 | 0.83 | n.s. |
| | Hull IoU | 0.665 | 0.608 | +0.057 | 0.15 | n.s. |
| Qwen3.6-27B | F1 macro | 0.690 | 0.657 | +0.033 | 0.21 | n.s. |
| | BA | 0.757 | 0.743 | +0.014 | 0.64 | n.s. |
| | Hull IoU | 0.665 | 0.612 | +0.053 | 0.13 | n.s. |

#### S6.6 Deployment cost, latency, and stability

*Cost and latency.*

Annotating one sample costs one LLM call per gating-tree node, so the call count is fixed by the tree and is independent of the number of cells in the sample. Calls inside an empty sub-tree are elided, so the effective count falls when a branch is rejected early. The per-call cost depends on the backbone and the serving environment. The open-weight Qwen3.6-27B runs on the single node described in section S6.4. Table S7 reports end-to-end wall-clock to annotate one sample on Acute2021 with GPT-5.4 under sequential execution, at all three tree depths. The per-step median is $4.5$–$5.6$ s, of which the LLM call is a roughly fixed $1$–$3$ s and the remainder is label propagation given the predicted gate. Steps within a tree level are independent given their parent, so the sequential figures are an upper bound.

Table S7. Per-sample cascade latency on Acute2021 (GPT-5.4, sequential execution), related to Table 2.

| Depth | Steps | Median | Mean (range) | Per-step |
|---|---|---|---|---|
| L1 | 22 | 123 s | 123 s (58–210) | 5.6 s |
| L2 | 41 | 216 s | 215 s (80–443) | 5.3 s |
| L3 | 62 | 277 s | 304 s (79–891) | 4.5 s |

*Stability.*

Table S5 reports seed stability of the per-step LLM-Gate rows. The cascade compounds those per-step decisions, so we additionally measure run-to-run reproducibility of the end-to-end cascade (the flat-vs.-hierarchical harnesses of section S9). Every cohort-depth setting is repeated three times with Qwen3.6-27B and reported as mean $\pm$ s.d. in Table S8. Spread is small (s.d. $\leq 0.045$ across all 17 settings) and does not grow with tree depth in any cohort (on Acute2020 it is $0.006/0.017/0.006$ at L1 / L2 / L3), so a deeper gating tree does not destabilize the resulting annotation. The two largest spreads are Acute2021 L1 ($\pm 0.045$) and Z74D-H ($\pm 0.039$), the latter a small-cohort effect at $n = 2$ rather than decoder noise.

Table S8. Run-to-run stability of the cascade (Qwen3.6-27B, three repeats, mean $\pm$ s.d. of multi-label F1 macro), related to Figure 5. $n$ = test samples. Bold = best.

| Cohort | Depth | $n$ | Flat C2S | LLM-Gate cascade |
|---|---|---|---|---|
| Acute2020 | L1 (16) | 12 | $0.249 \pm 0.006$ | $\mathbf{0.551} \pm 0.006$ |
| Acute2020 | L2 (38) | 12 | $0.119 \pm 0.003$ | $\mathbf{0.444} \pm 0.017$ |
| Acute2020 | L3 (83) | 12 | $0.043 \pm 0.002$ | $\mathbf{0.257} \pm 0.006$ |
| Acute2021 | L1 (16) | 33 | $0.243 \pm 0.003$ | $\mathbf{0.369} \pm 0.045$ |
| Acute2021 | L2 (38) | 33 | $0.123 \pm 0.000$ | $\mathbf{0.308} \pm 0.022$ |
| Acute2021 | L3 (83) | 33 | $0.037 \pm 0.002$ | $\mathbf{0.176} \pm 0.017$ |
| Vaccine | L1 (16) | 85 | $0.303 \pm 0.002$ | $\mathbf{0.536} \pm 0.006$ |
| Vaccine | L2 (38) | 85 | $0.160 \pm 0.001$ | $\mathbf{0.453} \pm 0.014$ |
| Vaccine | L3 (83) | 85 | $0.054 \pm 0.002$ | $\mathbf{0.257} \pm 0.006$ |
| Bjornson | full | 32 | $0.471 \pm 0.012$ | $\mathbf{0.871} \pm 0.012$ |
| Z74D-H | full | 2 | $0.268 \pm 0.015$ | $\mathbf{0.563} \pm 0.039$ |
| Z74D-T | full | 15 | $0.234 \pm 0.006$ | $\mathbf{0.399} \pm 0.019$ |
| FRDR | full | 111 | $\mathbf{0.187} \pm 0.002$ | $0.114 \pm 0.006$ |
| LP-D | full | 24 | $0.078 \pm 0.006$ | $\mathbf{0.248} \pm 0.011$ |
| LP-B | full | 24 | $0.083 \pm 0.010$ | $\mathbf{0.546} \pm 0.022$ |
| LP-T | full | 24 | $0.141 \pm 0.008$ | $\mathbf{0.657} \pm 0.003$ |
| LP-Tr | full | 24 | $0.028 \pm 0.003$ | $\mathbf{0.226} \pm 0.005$ |

### S7 Visual-modality ladder: implementation detail

This section details the VLM-Gate and Agent-Gate harnesses summarized in the main text and reported in Figure 3 of the main text. Both share LLM-Gate's text prompt, system prompt, output schema (one axis-aligned rectangle $B_c \subseteq \mathbb{R}^2$ per candidate), and per-cell propagation (rectangle containment with *Unassigned* residual). They differ only in what the user message carries and whether the model is given a tool to call.

*VLM-Gate.*

The user message gains a single PNG image, a decoration-free 2-D scatter of the parent population in the $(x_v, y_v)$ plane, produced by a pre-render pass (matplotlib, fixed arcsinh axes, no title/legend/grid). Text blocks, system prompt, output schema, and per-cell propagation are byte-identical to LLM-Gate. Only the user message gains the image part.

*Agent-Gate.*

On top of VLM-Gate, the model is given one tool, `render_gate_overlay(rectangle)`, which re-renders the model's proposed rectangle on top of the parent scatter and returns the resulting overlay PNG. The conversation runs for up to four turns with the following turn-by-turn `tool_choice` schedule:

- **Turn 1:** `tool_choice="required"`. Without it the model skips the loop entirely and commits on the first turn.
- **Turns 2–3:** `tool_choice="auto"`. The model decides whether to revise its rectangle and call the tool again, or commit.
- **Turn 4:** `tool_choice="none"`. The model must commit a final rectangle in JSON form.

Output schema and per-cell propagation are unchanged from LLM-Gate. Overlay rendering happens in-process. The overlay images are streamed back to the model and not persisted.

*Extended numerical detail.*

By sign test on the 23,041 paired (cohort, sample, step, category) instances summarized in the main text, Agent-Gate emits a smaller rectangle than LLM-Gate in $72\%$ of instances (vs. $26\%$ larger) and smaller than VLM-Gate in $62\%$ (vs. $31\%$ larger), both well above the chance rate of $50\%$. The Agent-Gate vs. VLM-Gate median area ratio is $0.96$. Mean turns per step on the 6,402-step Agent-Gate run is $2.07$ (median 2), with the forced first-turn call accounting for $\sim 93\%$ of all tool invocations. Only $5$–$12\%$ of steps per cohort trigger a second tool call. Extending `max_turns` would therefore be unlikely to alter the average outcome.

*Replication on an open-weight backbone.*

Table S10 repeats the gate-area measurement with Qwen3.5-27B. Agent-Gate's median rectangle is smaller than LLM-Gate's in all 11 cohorts, ranging from $-3\%$ to $-62\%$, so the tightening induced by the self-verification loop is not a property of the closed-weight backbone. The intermediate VLM-Gate step does not replicate as cleanly. On Qwen3.5-27B its median area exceeds LLM-Gate's in three cohorts (Z74D-H, Z74D-T, LP-T), whereas on GPT-5.4 it shrinks monotonically in all 11. Adding the image is therefore backbone-dependent. Adding the loop is not.

Table S9. Per-cohort visual-modality results on Qwen3.5-27B, related to Figure 3, expanding the Qwen3.5-27B mean line of Figure 3A. Bold = best per metric.

| | LLM-Gate | | VLM-Gate | | Agent-Gate | |
|---|---|---|---|---|---|---|
| Cohort | F1 | IoU | F1 | IoU | F1 | IoU |
| A2020 | **0.667** | **0.607** | **0.667** | **0.607** | 0.649 | 0.570 |
| A2021 | **0.656** | **0.598** | 0.649 | 0.587 | 0.630 | 0.548 |
| Vacc | 0.673 | 0.631 | **0.678** | **0.635** | 0.650 | 0.585 |
| Bj | 0.816 | **0.787** | **0.827** | 0.760 | 0.792 | 0.709 |
| Z74D-H | 0.631 | 0.674 | **0.637** | **0.690** | 0.628 | 0.659 |
| Z74D-T | 0.549 | 0.605 | **0.559** | **0.607** | 0.550 | 0.575 |
| FRDR | 0.680 | 0.636 | **0.698** | **0.646** | 0.672 | 0.589 |
| LP-D | 0.631 | 0.502 | **0.689** | **0.550** | 0.645 | 0.481 |
| LP-B | **0.650** | **0.548** | 0.604 | 0.510 | 0.581 | 0.461 |
| LP-T | **0.622** | **0.630** | 0.610 | 0.628 | 0.602 | 0.622 |
| LP-Tr | 0.640 | 0.531 | **0.674** | **0.571** | 0.657 | 0.534 |
| *Mean* | 0.656 | 0.613 | **0.663** | **0.617** | 0.642 | 0.576 |

Table S10. Gate area on an open-weight backbone (Qwen3.5-27B), related to Figure 3. Per-cohort median rectangle area in the $(x_v, y_v)$ arcsinh plane; *Shrink* = Agent-Gate relative to LLM-Gate.

| Cohort | LLM-Gate | VLM-Gate | Agent-Gate | Shrink |
|---|---|---|---|---|
| A2020 | 8.10 | 7.60 | 6.72 | −17% |
| A2021 | 6.16 | 6.00 | 5.10 | −17% |
| Vacc | 7.42 | 7.00 | 6.11 | −18% |
| Bj | 13.61 | 10.26 | 8.25 | −39% |
| Z74D-H | 9.84 | 10.48 | 9.53 | −3% |
| Z74D-T | 11.86 | 12.14 | 10.81 | −9% |
| FRDR | 18.80 | 15.53 | 12.78 | −32% |
| LP-D | 3.59 | 3.20 | 2.52 | −30% |
| LP-B | 3.50 | 2.05 | 1.32 | −62% |
| LP-T | 3.75 | 3.94 | 3.43 | −8% |
| LP-Tr | 3.05 | 2.78 | 2.20 | −28% |

*Qualitative comparison.*

Figure S1 renders the three methods' rectangles side-by-side on four representative steps. The monotone shrinkage from LLM-Gate → VLM-Gate → Agent-Gate is visible per row. On Bjornson step 7 (CD16 × CD11b), for example, Agent-Gate's monocyte rectangle retains only the core of the cluster while LLM-Gate covers the full distribution. The truncated periphery falls into *Unassigned* rather than misrouting to a neighboring candidate, so label-level F1 absorbs only a mild penalty even when Hull IoU drops sharply.

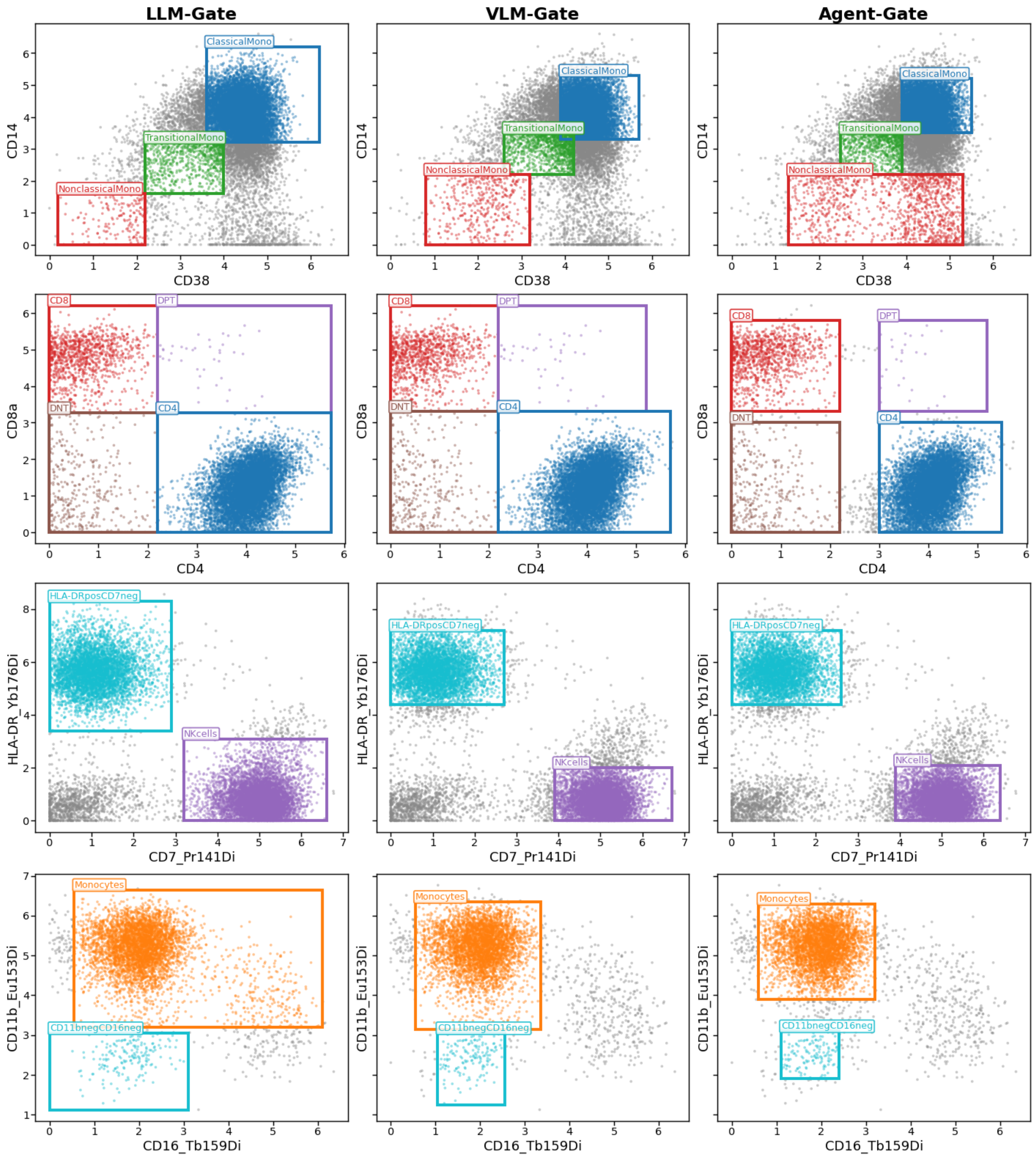


Figure S1. Qualitative gate comparison: visual-modality ladder, related to Figures 2 and 3. Rows: four steps from Acute2020 sample 994570 (steps 18, 29) and Bjornson sample R15W11 (steps 6, 7). Columns: LLM-Gate, VLM-Gate, Agent-Gate. Gray = parent cells; colored = candidate assignments by rectangle containment, boundary in matching color. LLM backbone: GPT-5.4.

### S8 Per-scenario robustness: scenario × cohort breakdown

This section expands Tables S13–S15 with per-cohort cell-counts within each robustness scenario. Each cell of Tables S14–S16 is the number of test-split eval instances triggered by the (scenario, cohort) pair (sum over its (cohort, step) defining triples of test-split sample counts). Empty cells indicate the scenario is not defined for that cohort. Tables S11–S12 below define each scenario.

#### S8.1 Scenario definitions

The depletion and calibration-drift experiments draw from two curated scenario libraries shipped with the benchmark (one for population depletion, one for calibration drift). Each scenario fixes *what* is perturbed and *where* (cohorts, gating steps), and is paired with a clinical or technical rationale grounding the perturbation in conditions a practicing immunologist would expect. The (scenario, cohort, step) triples are listed in Tables S14, S16.

Table S11. The nine clinical population-depletion scenarios, related to Figure 4. Each scenario sub-samples the named population at the relevant gating step to a target prevalence $\rho \in \{10\%, 1\%\}$ of nominal (the unperturbed data serve as the $\rho = 100\%$ reference in Table S13); the category list is left intact so the test is whether the model correctly labels the populations that remain.

| # | Scenario | Depleted population | Clinical rationale |
|---|---|---|---|
| D1 | `hiv` | $CD4^+$ T cells | HIV late-stage / anti-CD4 therapy |
| D2 | `rituximab` | $CD20^+$ B cells | Anti-CD20 (rituximab, ocrelizumab) |
| D3 | `b_cell_aplasia` | $CD19^+$ B cells | Marrow B-cell aplasia (post anti-CD20) |
| D4 | `immunosenescence` | Naive T cells | Elderly thymic involution |
| D5 | `neutropenia` | Granulocytes / PMN | Post-chemotherapy / idiopathic |
| D6 | `pdc_depletion` | Plasmacytoid DCs | HIV, severe COVID-19, measles |
| D7 | `nonclassical_mono_loss` | Patrolling ($CD14^{dim}$ $CD16^+$) monocytes | Severe COVID-19 / sepsis |
| D8 | `th17_axis_collapse` | Th17 cells | Secukinumab / ustekinumab (anti-IL17 / anti-IL23) |
| D9 | `treg_depletion_daclizumab` | Tregs ($CD25^{hi}$ $FoxP3^+$) | Daclizumab (anti-CD25) / IPEX |

Table S12. The nine synthetic calibration-drift scenarios, related to Figure 4. Each scenario rescales the listed channels by a multiplicative factor on raw signal, calibrated to push 10–30% of cells across the gate boundary (the lot-variability regime); GT labels are unchanged. “dim” = factor $< 1$, “bright” = factor $> 1$.

| # | Scenario | Markers (direction) | Technical rationale |
|---|---|---|---|
| C1 | `lineage_dim` | CD3, CD4, CD8, CD19, CD20 (dim) | Aged antibody / partial conjugation |
| C2 | `ccr7_internalization` | CCR7 (dim) | Freeze-thaw or in-tube CCR7 internalization |
| C3 | `cd45ra_drift` | CD45RA (bright) | Lot-to-lot brightness increase |
| C4 | `treg_marker_collapse` | FoxP3, CD25 (dim); CD127 (bright) | Intracellular Treg-marker dim staining |
| C5 | `mono_subset_shift` | CD16 (bright) | CD16 spillover at monocyte subset boundaries |
| C6 | `dc_marker_drift` | CD11c, CD123, CD303 (dim) | Lot variability on narrow-dynamic-range DC markers |
| C7 | `bead_residual` | Bead channels (dim, mass cytometry) | Post-EQ4 bead-normalization residual |
| C8 | `viability_drift` | Live/Dead (bright) | Storage / freeze-thaw intensifying viability stain |

| # | Scenario | Markers (direction) | Technical rationale |
|---|---|---|---|
| C9 | `panel_global` | All `protein` channels (bright) | Whole-panel calibration drift (instrument-level) |

#### S8.2 Population depletion (scenario × cohort)

Table S13 expands Figure 4 of the main text to the full nine-scenario × three-$\rho$ grid, with per-row results for UNITO (fit on train+val, not retrained per $\rho$), cyMAE (a train+val-fit foundation model), and the zero-shot GPT-5.4 and Qwen3.6-27B LLM-Gate rows. Each cell reports *recall on the depleted candidate* / *recall on the non-depleted candidates* at that step, averaged across per-sample eval instances.

Table S13. Per-scenario depletion robustness (numerical companion to Figure 4 of the main text), related to Figure 4. Scenario codes `D1`–`D9` are defined in Table S11. Recall on the depleted candidate / recall on the non-depleted candidates at the same gating step at $\rho \in \{100\%, 10\%, 1\%\}$ of nominal prevalence; $\rho = 100\%$ is the nominal reference. UNITO and cyMAE are fit on train+val at nominal composition and not retrained per $\rho$; GPT-5.4 and Qwen3.6-27B are zero-shot. *Mean* averages the 9 scenarios.

| Method (Train) | $\rho$ | D1 | D2 | D3 | D4 | D5 | D6 | D7 | D8 | D9 | *Mean* |
|---|---|---|---|---|---|---|---|---|---|---|---|
| UNITO (train+val) | 100% | 0.990/0.904 | 0.994/0.780 | 0.981/0.993 | 0.916/0.836 | 0.866/0.930 | 0.411/0.483 | 0.000/0.926 | 0.883/0.935 | 0.805/0.954 | 0.761/0.860 |
| | 10% | 0.741/0.851 | 0.996/0.367 | 0.925/0.989 | 0.807/0.771 | 0.894/0.892 | 0.386/0.491 | 0.000/0.917 | 0.849/0.926 | 0.777/0.951 | 0.708/0.795 |
| | 1% | 0.294/0.814 | 0.990/0.031 | 0.770/0.982 | 0.638/0.749 | 0.735/0.841 | 0.259/0.487 | 0.000/0.905 | 0.769/0.925 | 0.725/0.942 | 0.575/0.742 |
| cyMAE (train+val) | 100% | 0.886/0.854 | 0.995/0.659 | 0.941/0.948 | 0.833/0.767 | 0.927/0.835 | 0.795/0.763 | 0.824/0.840 | 0.796/0.752 | 0.827/0.766 | 0.869/0.798 |
| | 10% | 0.886/0.854 | 0.995/0.659 | 0.943/0.948 | 0.833/0.767 | 0.926/0.835 | 0.797/0.763 | 0.813/0.840 | 0.796/0.751 | 0.857/0.766 | 0.872/0.798 |
| | 1% | 0.889/0.854 | 0.996/0.657 | 0.946/0.948 | 0.834/0.767 | 0.927/0.835 | 0.784/0.763 | 0.861/0.840 | 0.790/0.751 | 0.849/0.766 | 0.875/0.798 |
| GPT-5.4 (ZS) | 100% | 0.981/0.925 | 0.999/0.578 | 0.866/0.937 | 0.955/0.811 | 0.962/0.813 | 0.825/0.827 | 0.797/0.778 | 0.803/0.956 | 0.502/0.901 | 0.855/0.836 |
| | 10% | 0.944/0.942 | 0.993/0.476 | 0.854/0.908 | 0.895/0.764 | 0.939/0.821 | 0.732/0.762 | 0.795/0.767 | 0.765/0.958 | 0.490/0.863 | 0.823/0.807 |
| | 1% | 0.919/0.801 | 0.983/0.353 | 0.844/0.863 | 0.909/0.755 | 0.872/0.729 | 0.556/0.688 | 0.775/0.753 | 0.755/0.957 | 0.562/0.786 | 0.797/0.743 |
| Qwen3.6-27B (ZS) | 100% | 0.999/0.957 | 1.000/0.353 | 0.997/0.997 | 0.973/0.867 | 0.955/0.861 | 0.935/0.889 | 0.649/0.811 | 0.797/0.955 | 0.336/0.885 | 0.849/0.842 |
| | 10% | 0.981/0.966 | 0.998/0.303 | 0.988/0.999 | 0.906/0.839 | 0.835/0.840 | 0.881/0.825 | 0.618/0.784 | 0.743/0.970 | 0.378/0.861 | 0.814/0.821 |
| | 1% | 0.927/0.918 | 0.996/0.258 | 0.967/0.989 | 0.909/0.830 | 0.639/0.712 | 0.907/0.778 | 0.572/0.765 | 0.733/0.971 | 0.324/0.827 | 0.775/0.783 |

Table S14. Per-cohort eval-instance counts $n$ per depletion scenario, related to Figure 4. Cohort abbreviations: A2020/A2021 = Acute2020/2021, Vacc = Vaccine, Z74D-H/T = FR-FCM-Z74D HC/Tissue, LP-T/Tr/B/D = Lyoplate T/Treg/B/DC, FRDR = FRDR COVID-19, Bj = Bjornson. *Total* matches per-scenario $n$ in Table S13.

| Scenario | A2020 | A2021 | Vacc | Z74D-H | Z74D-T | LP-T | LP-Tr | LP-B | LP-D | FRDR | Bj | *Total* |
|---|---|---|---|---|---|---|---|---|---|---|---|---|
| hiv | 12 | 33 | 85 | – | 15 | 24 | – | – | – | – | 32 | 201 |
| rituximab | – | – | – | – | – | – | – | 24 | – | – | – | 24 |
| b_cell_aplasia | 12 | 33 | 85 | – | – | – | – | – | – | 111 | 32 | 273 |
| immunosenescence | 24 | 66 | 170 | 4 | 30 | 48 | – | – | – | – | – | 342 |
| neutropenia | 12 | 33 | 85 | – | – | – | – | – | – | 111 | 32 | 273 |
| pdc_depletion | 12 | 33 | 85 | – | – | – | – | – | 24 | 111 | – | 265 |
| nonclassical_mono_loss | – | – | – | – | – | – | – | – | – | 111 | – | 111 |
| th17_axis_collapse | 12 | 33 | 85 | 2 | 15 | – | – | – | – | – | – | 147 |
| treg_depletion_daclizumab | – | – | – | 2 | 15 | – | 24 | – | – | – | – | 41 |
| *Total* | 84 | 231 | 595 | 8 | 75 | 72 | 24 | 24 | 24 | 444 | 96 | 1677 |

#### S8.3 Calibration drift (scenario × cohort)

Table S15 expands the calibration-drift companion to Figure 4 of the main text to the full nine-scenario grid, reporting ΔF1 macro / ΔHull IoU (perturbed minus nominal) for UNITO, cyMAE, and the zero-shot GPT-5.4 and Qwen3.6-27B LLM-Gate rows under each calibration-drift scenario.

Table S15. Per-scenario calibration-drift robustness, related to Figure 4. Scenario codes C1–C9 are defined in Table S12. $\Delta = F_{\text{pert}} - F_{\text{nom}}$ for F1 macro / Hull IoU under nine synthetic channel-scale perturbations. UNITO and cyMAE are fit on train+val at nominal calibration and not retrained per scenario; GPT-5.4 and Qwen3.6-27B are zero-shot. *Mean* averages the 9 scenarios.

| Method (Train) | C1 | C2 | C3 | C4 | C5 | C6 | C7 | C8 | C9 | *Mean* |
|---|---|---|---|---|---|---|---|---|---|---|
| UNITO (train+val) | −0.053/ −0.096 | −0.028/ −0.041 | −0.091/ −0.123 | −0.011/ −0.024 | −0.012/ −0.020 | +0.029/ +0.026 | −0.053/ −0.015 | −0.002/ +0.006 | −0.021/ −0.036 | −0.027/ −0.036 |
| cyMAE (train+val) | −0.132/ −0.107 | −0.045/ −0.069 | −0.026/ −0.035 | −0.018/ +0.026 | −0.012/ −0.010 | −0.082/ −0.069 | −0.002/ −0.001 | −0.021/ −0.025 | −0.027/ −0.031 | −0.040/ −0.036 |
| GPT-5.4 (ZS) | +0.012/ +0.003 | −0.010/ −0.013 | −0.012/ −0.016 | −0.026/ −0.038 | −0.012/ −0.011 | −0.004/ −0.034 | +0.005/ +0.001 | −0.064/ −0.078 | −0.012/ −0.013 | −0.014/ −0.022 |
| Qwen3.6-27B (ZS) | +0.012/ +0.018 | −0.008/ +0.000 | −0.020/ −0.024 | −0.036/ −0.031 | −0.005/ −0.002 | +0.012/ −0.023 | +0.000/ +0.013 | −0.085/ −0.111 | +0.002/ −0.003 | −0.014/ −0.018 |

Table S16. Per-cohort eval-instance counts $n$ for each calibration-drift scenario, related to Figure 4. Cohort abbreviations as in Table S14. *Total* sums across cohorts (equals the per-scenario $n$ in Table S15).

| Scenario | A2020 | A2021 | Vacc | Z74D-H | Z74D-T | LP-T | LP-Tr | LP-B | LP-D | FRDR | Bj | *Total* |
|---|---|---|---|---|---|---|---|---|---|---|---|---|

| Scenario | A2020 | A2021 | Vacc | Z74D-H | Z74D-T | LP-T | LP-Tr | LP-B | LP-D | FRDR | Bj | *Total* |
|---|---|---|---|---|---|---|---|---|---|---|---|---|
| lineage_dim | 24 | 33 | 85 | 2 | – | 24 | – | 24 | – | 111 | – | 303 |
| ccr7_internalization | – | – | – | 4 | 30 | 48 | – | – | – | – | – | 82 |
| cd45ra_drift | 24 | 66 | 170 | 4 | – | 48 | – | – | – | – | – | 312 |
| treg_marker_collapse | – | – | – | 2 | 15 | – | 24 | – | – | – | – | 41 |
| mono_subset_shift | 12 | 33 | 85 | – | – | – | – | – | – | 222 | – | 352 |
| dc_marker_drift | 12 | – | – | – | – | – | – | – | 24 | 111 | – | 147 |
| bead_residual | 12 | 33 | 85 | – | – | – | – | – | – | – | – | 130 |
| viability_drift | 12 | 33 | 85 | – | – | – | – | – | – | – | – | 130 |
| panel_global | 12 | 33 | 85 | 2 | 15 | 24 | 24 | 24 | 24 | 111 | 32 | 386 |
| *Total* | 108 | 231 | 595 | 14 | 60 | 144 | 48 | 48 | 48 | 555 | 32 | 1883 |

### S9 Flat-vs-hierarchical comparison: pipeline details

This section specifies the two harnesses compared in the flat-vs. hierarchical analysis (Figure 5 of the main text). Both are run with the same two backbones (GPT-5.4 and the open-weight Qwen3.6-27B), one canonical input parquet per sample, and one leaf vocabulary derived from the same gating tree. Generation settings (temperature, max completion tokens) follow section S6.3. They differ only in how the per-cell labeling is decomposed and which LLM call boundary the model sees.

*Shared leaf vocabulary.*

Given a gating tree, the leaf set is the set of categories that never appear in the parent expression of any later step. Non-leaf categories are intermediate gating waypoints (e.g., *Mononuclear*, *CD45*$^+$, $\alpha\beta T$) and are filtered out. A cell that never reaches a leaf along its predicted path collapses to the synthetic class *Discard*, which is scored as a regular category. We sweep three tree depths, namely L1 (16 lineage leaves), L2 (38 identity leaves), and L3 (the full 83-leaf tree, the $62$-step canonical Acute2020 tree). Both harnesses are evaluated against the same depth in each comparison cell of Figure 5 of the main text.

*Flat C2S harness.*

The harness clusters the entire sample once with FlowSOM and asks the LLM to label every cluster with a multi-label leaf set in a single whole-sample call.

*Clustering.* A $10 \times 10$ self-organizing map (random initialization, $30{,}000$ sampled training steps) is trained on all protein channels of the sample (arcsinh-transformed, no z-scoring). The SOM codebook (100 nodes) is then collapsed to $K$ metaclusters via average-linkage agglomerative clustering, with $K$ set to the leaf count at each depth ($16/38/83$). Each cell is assigned to its nearest codebook node and inherits that node's metacluster. The implementation uses the `MiniSom` library for the self-organizing map and scikit-learn's `AgglomerativeClustering` for the metacluster collapse. Per metacluster we record cell count, fraction, and a marker centroid (mean over cluster members) plus a per-cluster cell sentence that orders markers from highest to lowest intra-cluster mean expression.

*Prompt.* A single text prompt is rendered per sample and contains (i) modality and cofactor; (ii) sample identifier with $n_{cells}$ and $K$; (iii) the flat leaf vocabulary plus the sentinel *Discard* (presented as a JSON-line list); (iv) the panel-wide HVP marker ranking; (v) one block per cluster with header `## cluster_k (n=N, X.XX%)` followed by the `Marker(value)` cell sentence sorted by descending value within that cluster. The system prompt instructs the model to return one JSON object keyed by cluster id, each value carrying a `leaves` list (drawn verbatim from the leaf vocabulary or the singleton `["Discard"]`) and a one-sentence `rationale`; multi-label outputs are explicitly permitted when parallel-axis steps in the tree apply (e.g., a CD4 T cell carrying both a memory leaf and an activation leaf).

*Per-cell propagation.* For each cluster the LLM's leaf set is decomposed back to step columns by inverting the leaf→step-column map computed at preprocess time. Cells outside any step's predicted parent receive *Unassigned* in that step's column. The multi-label evaluator collapses cells whose final path contains no leaf to *Discard*.

*LLM-Gate cascade harness.*

The harness walks the gating tree step by step, issuing one LLM call per step.

*Cascade state.* A per-sample state object holds (a) the original cell×marker frame and (b) the predicted $\text{Step}_k$ column for every step the cascade has finished. Ground-truth $\text{Step}_k$ columns are masked from the parent-expression evaluator. At step $k$ the parent population is computed by evaluating the tree's parent expression (e.g., `Step01 == CD3pos & Step02 == CD4`) against the LLM's own predictions for steps $1..k-1$. Errors therefore compound across depth, but every step sees the population the cascade actually produced rather than an oracle parent.

*Per-step LLM call.* The step's marker pair $(x_v, y_v)$ is resolved against the panel's marker→channel map, the predicted-parent mask is computed, and a biaxial-plot prompt is rendered with (i) modality, cofactor, and a path breadcrumb (the parent expression and the step number); (ii) the candidate-child set $C_v \cup \{\text{Unassigned}\}$ verbatim; (iii) per-axis distribution summaries (decile-based bin centers and counts on each marker); (iv) a per-step note (the gating tree's per-step description, optionally overridden by a curated per-(cohort, step) note); (v) the unified rectangle output schema. The system prompt is the same one used by the zero-shot panel evaluation. A `tiebreak=smallest` rule resolves cells contained in multiple gates by assigning the smallest-area gate, which empirically matches a manual annotator's preference for the more specific child.

*Per-cell propagation and sub-tree skip.* The LLM's gate JSON is parsed via the rectangle schema. Cells inside a single gate take that gate's category, and cells in no gate become *Unassigned*. The full-length step column is constructed by writing the per-cell labels onto the parent indices and *Unassigned* on the rest, then committed to the cascade state. If a step's predicted parent mask is empty (e.g., a prior step rejected every cell along that branch), the runner skips just that sub-tree and continues evaluating the rest of the gating tree. The skipped step's column is written as *Unassigned* on every cell and the skip is recorded in the cascade metadata. The multi-label evaluator reads the cascade's per-cell Step columns the same way it reads Flat C2S's cluster-decomposed columns, so the two methods score against the same evaluation under the same leaf vocabulary.

*Cost.* Flat C2S issues exactly one LLM call per sample regardless of tree depth. The LLM-Gate cascade issues at most one call per gating-tree step ($22$, $41$, $62$ for L1, L2, L3 on Acute2020). Calls inside an empty sub-tree are elided.

*Evaluator.*

In this comparison a cell's label is genuinely *multi-label*, even at the leaf level. The gating tree branches along parallel axes, so a single cell can satisfy more than one leaf at once. A $CD4^+$ T cell, for example, may carry both a differentiation leaf (*memory*) and an activation leaf (*activated*). Each cell's ground truth and each method's prediction are therefore a *set* of leaves, drawn from the leaf vocabulary plus the sentinel *Discard*, rather than a single label. The shared evaluator encodes each cell's leaf set as a binary indicator vector over that vocabulary with `MultiLabelBinarizer`, the multi-label one-hot encoder from scikit-learn's `preprocessing` module, and then scores both methods against ground truth using scikit-learn's multi-label metrics, namely F1 (micro / macro / weighted), Jaccard (micro / macro), Hamming loss, and subset accuracy. The macro and micro F1 reported in Figure 5 of the main text are these multi-label scores. As a secondary view, the evaluator also collapses each cell to a single *primary leaf* (the deepest leaf on its predicted path, or *Discard* when the path reaches none) and reports accuracy, balanced accuracy, and a confusion matrix from it. Cells that the two harnesses emit in different row orders are aligned by parquet row index, and per-leaf precision / recall / F1 with the ground-truth support distribution are written as side outputs that do not appear in the main table.

#### S9.1 Flat C2S prompt

This subsection reproduces the system and user prompts driving Flat C2S, illustrated on Acute2020/994570_Normalized at L3, the full $83$-leaf tree ($K = 83$ FlowSOM metaclusters). The user prompt is rendered programmatically per sample. The system prompt is fixed across samples and depths.

*Prompt construction in three pieces.*

The cluster-level cell sentence is the workhorse representation, adapted from the cell-sentence framing of C2S[16] but restated at the *cluster* level rather than the per-cell level. Each FlowSOM metacluster is summarized by its arcsinh-mean expression vector over the panel, serialized into a single line of `Marker(value)` tokens sorted by descending value within that cluster. This makes the high-expression markers (the immunological lineage signature) read first, mirroring how an annotator scans a marker profile. The cluster header carries the cell count and population fraction. The panel-wide HVP ranking precedes the cluster blocks as a global cue. The model is asked for a multi-label leaf assignment per cluster, drawn from the gating tree’s leaf vocabulary plus the synthetic sentinel *Discard*.

**Flat C2S system prompt**

```
You are an expert immunologist annotating cytometry clusters at the
WHOLE-CELL level - one shot per FlowSOM meta-cluster.

Each user message describes ONE sample with N FlowSOM meta-clusters.
For EVERY cluster you must return a JSON list of leaf cell-type
labels (multi-label) drawn ONLY from the user message's
[Leaf categories] section, plus the sentinel "Discard" when no leaf
fits.

## Cluster sentence
Each cluster header reads `## cluster_K (n=N_cells, X.XX%)' - the
cluster's cell count and its share of the sample. Cluster size is a
confidence prior on commitment stability (large clusters give stable
marker averages, tiny clusters <1% are noisier observations) - it
is NOT a category prior: both rare and abundant cell types can show
up as large or small FlowSOM clusters depending on the SOM grid
resolution.

The line below the header lists each cluster's marker profile as
`Marker(arcsinh_value)' tokens, separated by `>' and ordered by
descending value within the cluster (high-to-low expression).
Markers are arcsinh-transformed.

## Leaves only - no waypoints
Pick ONLY leaf labels from the [Leaf categories] section.
Intermediate gating waypoints (e.g. "CD3pos", "CD4", "CD8") are NOT
in the leaf set and must NOT appear in your output.

## Multi-label when parallel axes apply
A single cluster may legitimately carry multiple leaves when the
gating tree has parallel axes (e.g. CD4 differentiation x CD4
activation: a CD4 cluster should pick BOTH a memory leaf AND an
activation leaf when the marker profile supports it). Each picked
leaf populates a different gating step downstream, so over-
```

```
restricting to a single leaf throws away signal.

## Bead vs Discard - DO NOT confuse them
- Bead (when present in [Leaf categories]) is a REAL leaf for
  calibration beads. Pick it ONLY when the cluster's marker profile
  matches calibration beads - typically very high signal on a bead-
  channel marker (e.g., 140Ce_Bead, 165Ho_Bead, 175Lu_Bead) AND no
  concurrent real-cell lineage signal (CD45-, CD3-, CD19-, CD14-,
  CD66b-, etc.). Beads carry no immune receptors - if the cluster
  expresses any lineage marker at meaningful intensity, it is NOT
  a bead.

- Discard is the synthetic sentinel for cluster profiles that
  resemble cleanup-rejected real-cell events: dead cells, doublets,
  debris, mis-acquisition artifacts, or populations not represented
  by ANY listed leaf.

Discard is NOT "I am unsure" - if the cluster's profile is at all
consistent with a listed leaf, commit to that leaf. But likewise,
do not force a leaf onto an obviously-bad cluster (extreme low
CD45 with no lineage signal, viability-positive dead-cell cluster,
doublet-like profile with conflicting markers): pick Discard.

## Positive vs. negative
A marker is "positive" or "negative" RELATIVE to the other clusters
in the same prompt - not against absolute axis-zero. Compare a
cluster's value to the spread you see across clusters. In pre-gated
panels, even "negative" clusters can sit at moderate-to-high
absolute arcsinh values.

## Output format
A single JSON object keyed by cluster id. Each value is
{"leaves": ["<leaf_a>", ...], "rationale": "<one short sentence>"}.
Rationale is a brief biology-grounded justification.

Respond ONLY with the JSON object - no prose or commentary outside
it.
```

**Flat C2S user prompt (excerpt, L3, 2 of 83 clusters)**

```
[Context]
Modality: cytof (cofactor 5.0)
Sample: 994570_Normalized (dataset: Acute2020) -- n_cells=585,732,
        n_clusters=83

[Leaf categories]
Basophil
Bead
CD45hiCD66bpos
CD4Naive/CD38pos
CD4Naive/activated
```

CD4Naive/activated+CD38pos
CD4TCM/CD38pos
CD4TCM/activated
CD4TCM/activated+CD38pos
CD4TEM1/CD38pos
CD4TEM1/activated
CD4TEM1/activated+CD38pos
(... 83 leaves total; CD8, monocyte, NK, B-cell, Th/Treg, and DC leaves omitted for space ...)
Discard

[Top HVP markers (high -> low variance)]
CD66b, CD45, CD11c, CD16, CD25_IL-2Ra, CD161, CD45RO, CD38, CD45RA, CD196_CCR6, CD56_NCAM, CD3, CD127_IL-7Ra, CD194_CCR4, HLA-DR, TCRgd, CD14, CD4, CD183_CXCR3, CD8a, CD123_IL-3R, CD19, CD28, CD27, CD20, CD197_CCR7, CD57, IgD, CD294, CD185_CXCR5

---

### cluster_1 (n=13,598, 2.32%)
CD16(5.76) > CD66b(4.28) > CD45(3.81) > CD11c(3.40) > CD45RO(2.76) > CD194_CCR4(1.94) > CD196_CCR6(1.93) > CD38(1.75) > TCRgd(1.41) > CD45RA(1.40) > CD8a(1.18) > CD127_IL-7Ra(0.97) > CD123_IL-3R(0.92) > HLA-DR(0.66) > CD25_IL-2Ra(0.65) > CD19(0.44) > CD4(0.40) > CD56_NCAM(0.37) > CD14(0.34) > CD27(0.33) > CD28(0.29) > CD20(0.29) > CD3(0.20) > CD183_CXCR3(0.19) > CD294(0.19) > IgD(0.18) > CD57(0.16) > CD161(0.14) > CD197_CCR7(0.14) > CD185_CXCR5(0.07)

### cluster_2 (n=38,773, 6.62%)
CD16(5.64) > CD66b(3.81) > CD45(3.70) > CD11c(3.16) > CD45RO(2.41) > CD38(2.27) > CD194_CCR4(2.21) > TCRgd(1.66) > CD45RA(1.03) > CD123_IL-3R(0.92) > CD196_CCR6(0.86) > CD8a(0.80) > CD19(0.66) > CD127_IL-7Ra(0.51) > CD4(0.41) > CD25_IL-2Ra(0.36) > HLA-DR(0.36) > CD56_NCAM(0.32) > CD14(0.32) > CD28(0.28) > CD27(0.28) > CD20(0.19) > CD3(0.18) > IgD(0.15) > CD161(0.14) > CD183_CXCR3(0.14) > CD57(0.14) > CD294(0.14) > CD197_CCR7(0.12) > CD185_CXCR5(0.07)

(... clusters 3-83 follow the same one-line format ...)

---

### Output Format
Return a single JSON object keyed by cluster id. For each cluster, `leaves' MUST be a JSON list whose elements come from [Leaf categories] (verbatim) - or the singleton ["Discard"]. `rationale' is one short sentence.

~~~
```json
{
  "cluster_1": {"leaves": ["<leaf_a>", "<leaf_b>"],
               "rationale": "..."},
  "cluster_2": {"leaves": ["<leaf_a>", "<leaf_b>"],
               "rationale": "..."}
}
```
~~~

**Flat C2S model response (2 clusters)**

```
{
  "cluster_1": {
    "leaves": ["CD45hiCD66bpos"],
    "rationale": "Very high CD66b with high CD45 and dominant
                  myeloid-like CD16/CD11c pattern fits
                  CD45hiCD66bpos granulocytes."
  },
  "cluster_2": {
    "leaves": ["CD45hiCD66bpos"],
    "rationale": "High CD66b and CD45 with strong CD16/CD11c
                  support a CD45hiCD66bpos granulocyte cluster."
  }
}
```

#### S9.2 Flat-vs-hierarchical: per-sample breakdown

Tables S17–S19 list per-sample multi-label F1 macro (F1m) and F1 micro (F1$\mu$) for both harnesses across the three tree depths. All 12 Acute2020 test samples are shown, with GPT-5.4 and FlowSOM $K$ set to the leaf count per depth (16/38/83). The cascade walks every step, with empty predicted-parent sub-trees skipped as described above, so every sample completes the rest of the tree.

Table S17. Per-sample results at L1 (16 leaves, 22-step tree), related to Figure 5. F1m = multi-label F1 macro; F1$\mu$ = micro.

| | Flat C2S | | LLM-Gate cascade | |
|---|---|---|---|---|
| Sample | F1m | F1$\mu$ | F1m | F1$\mu$ |
| 994570_Normalized | 0.209 | 0.073 | 0.464 | 0.850 |
| 994572_Normalized | 0.298 | 0.723 | 0.567 | 0.862 |
| 994574_Normalized | 0.182 | 0.654 | 0.511 | 0.858 |
| 994585_Normalized | 0.243 | 0.708 | 0.471 | 0.847 |
| 994586_Normalized | 0.202 | 0.690 | 0.507 | 0.899 |
| 994587_Normalized | 0.187 | 0.750 | 0.571 | 0.871 |
| 994591_Normalized | 0.216 | 0.052 | 0.496 | 0.913 |
| HD2020_014_Normalized | 0.252 | 0.390 | 0.602 | 0.670 |
| HD2020_023F_Normalized | 0.190 | 0.116 | 0.504 | 0.864 |
| HD2020_029_Normalized | 0.394 | 0.764 | 0.634 | 0.834 |
| HD2020_051_Normalized | 0.409 | 0.248 | 0.719 | 0.917 |
| ND_Zeus_Normalized | 0.264 | 0.296 | 0.453 | 0.736 |
| **Mean** | **0.254** | **0.455** | **0.541** | **0.843** |

Table S18. Per-sample results at L2 (38 leaves, 41-step tree), related to Figure 5.

| | Flat C2S | | LLM-Gate cascade | |
|---|---|---|---|---|
| Sample | F1m | F1$\mu$ | F1m | F1$\mu$ |
| 994570_Normalized | 0.156 | 0.558 | 0.308 | 0.838 |
| 994572_Normalized | 0.212 | 0.621 | 0.449 | 0.823 |
| 994574_Normalized | 0.055 | 0.631 | 0.366 | 0.756 |
| 994585_Normalized | 0.146 | 0.671 | 0.409 | 0.856 |
| 994586_Normalized | 0.073 | 0.692 | 0.432 | 0.898 |
| 994587_Normalized | 0.084 | 0.667 | 0.462 | 0.654 |
| 994591_Normalized | 0.169 | 0.586 | 0.402 | 0.907 |
| HD2020_014_Normalized | 0.150 | 0.306 | 0.567 | 0.681 |
| HD2020_023F_Normalized | 0.164 | 0.394 | 0.384 | 0.842 |
| HD2020_029_Normalized | 0.246 | 0.538 | 0.590 | 0.838 |
| HD2020_051_Normalized | 0.211 | 0.193 | 0.440 | 0.881 |
| ND_Zeus_Normalized | 0.207 | 0.582 | 0.448 | 0.778 |
| **Mean** | **0.156** | **0.537** | **0.438** | **0.813** |

Table S19. Per-sample results at L3 (83 leaves, 62-step tree), related to Figure 5.

| | Flat C2S | | LLM-Gate cascade | |
|---|---|---|---|---|
| Sample | F1m | F1$\mu$ | F1m | F1$\mu$ |
| 994570_Normalized | 0.055 | 0.477 | 0.150 | 0.844 |
| 994572_Normalized | 0.047 | 0.421 | 0.269 | 0.836 |
| 994574_Normalized | 0.025 | 0.273 | 0.244 | 0.754 |
| 994585_Normalized | 0.050 | 0.480 | 0.224 | 0.808 |
| 994586_Normalized | 0.027 | 0.626 | 0.254 | 0.919 |
| 994587_Normalized | 0.026 | 0.440 | 0.273 | 0.739 |
| 994591_Normalized | 0.057 | 0.465 | 0.348 | 0.913 |
| HD2020_014_Normalized | 0.063 | 0.271 | 0.279 | 0.620 |
| HD2020_023F_Normalized | 0.075 | 0.381 | 0.197 | 0.833 |
| HD2020_029_Normalized | 0.103 | 0.491 | 0.310 | 0.786 |
| HD2020_051_Normalized | 0.098 | 0.598 | 0.224 | 0.882 |
| ND_Zeus_Normalized | 0.100 | 0.602 | 0.105 | 0.723 |
| **Mean** | **0.061** | **0.460** | **0.240** | **0.805** |

#### S9.3 Flat-vs-hierarchical: per-cell projections, confusion, and cascade traces

To complement the aggregate scores in Figure 5 of the main text and the per-sample numbers above, this subsection presents the per-cell behavior of both harnesses on the L1/L2/L3 sweep, all on the same representative sample (Acute2020/994570_Normalized, ~586K cells, with a 5K-cell UMAP subsample shared across panels). Figures S2–S4 show the L1/L2/L3 UMAP views. Figures S5–S7 report the row-normalized primary-leaf confusion matrices for both methods at each depth, and Figures S8–S10 show the LLM-Gate cascade's per-step biaxial gates as the cascade unfolds.

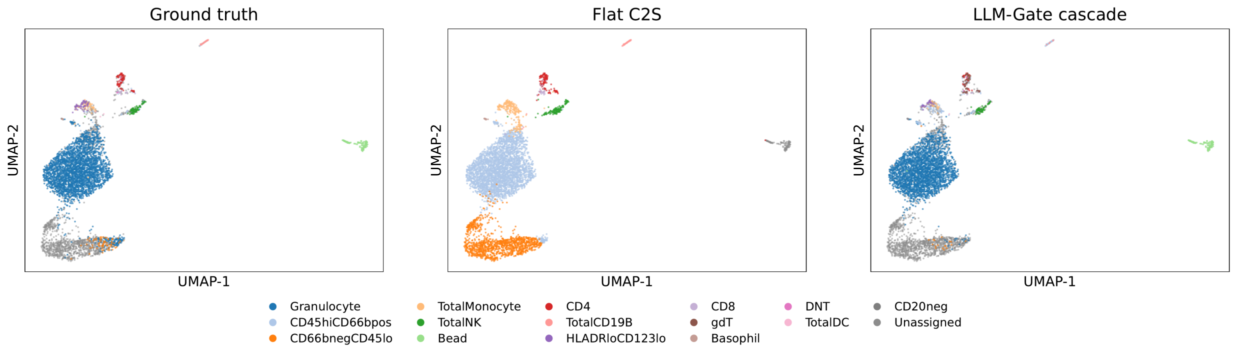


Figure S2. Per-cell UMAP at L1 depth (16 leaves) on Acute2020/994570_Normalized; cells colored by primary leaf under (left) ground truth, (middle) Flat C2S, (right) LLM-Gate cascade, related to Figure 5. The cascade reproduces manual-gating structure including cleanup leaves (*Bead*, *Unassigned*); Flat C2S smears these into adjacent lineages.

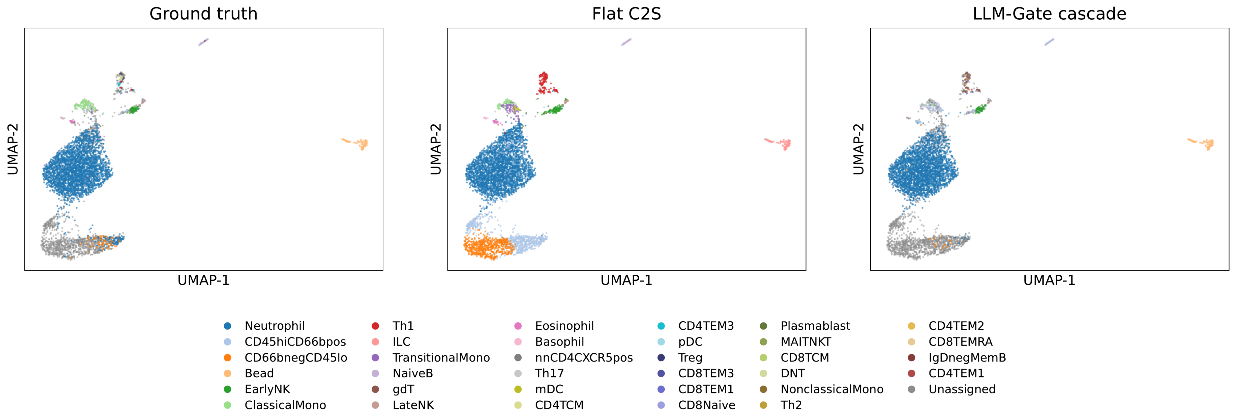


Figure S3. Per-cell UMAP at L2 depth (38 identity-level leaves, 41-step tree); same sample and embedding as Figure S2, related to Figure 5. As the leaf vocabulary expands beyond lineage to identity-level (memory/effector/naive subsets), Flat C2S still resolves the major lineage islands but mislabels finer T-subset territory; the LLM-Gate cascade preserves both the cleanup mass and the lineage-island boundaries.

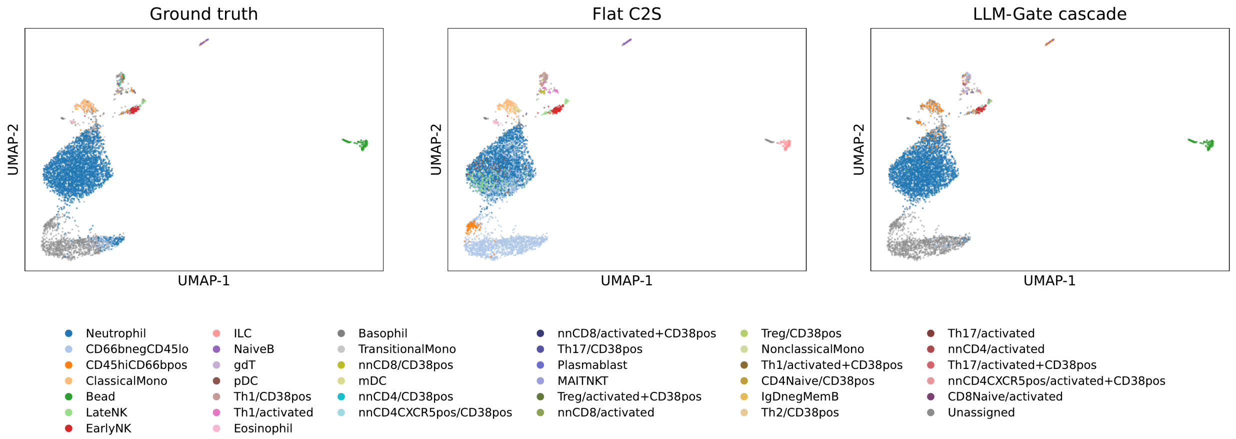


Figure S4. Per-cell UMAP at L3 depth (83 leaves, 62-step full canonical tree); same sample and embedding as Figures S2 and S3, related to Figure 5. At full depth the leaf vocabulary fragments the previously-coherent lineage islands into many activation/CD38 subsets; Flat C2S's cluster-to-leaf mapping degrades sharply (cf. Figure 5 of the main text), while the cascade retains the coarse structure and only loses resolution at the deepest activation-state leaves.

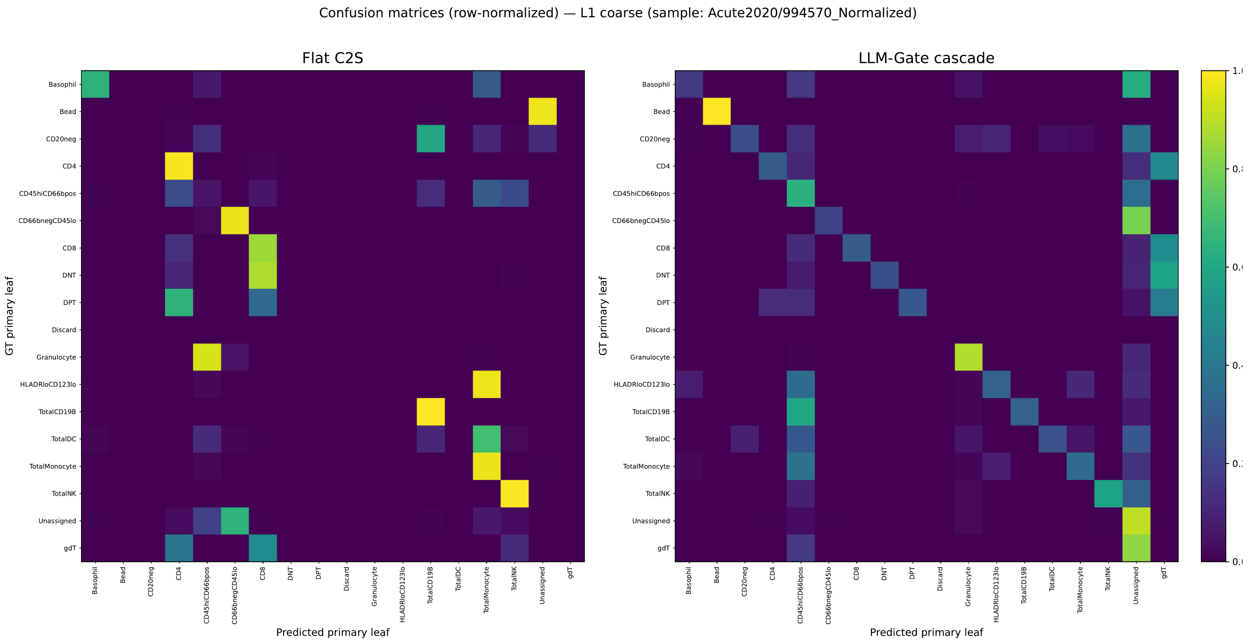


Figure S5. Row-normalized primary-leaf confusion matrices at L1 (Flat C2S vs LLM-Gate cascade), Acute2020/994570_Normalized, related to Figure 5. Diagonal mass on the cascade panel is concentrated on *Bead*, *Granulocyte*, and the major lineage classes; the Flat C2S panel shows the same mass distributed off-diagonal because the harness rarely emits the cleanup-class leaves (*Bead*, *Discard*) at all.

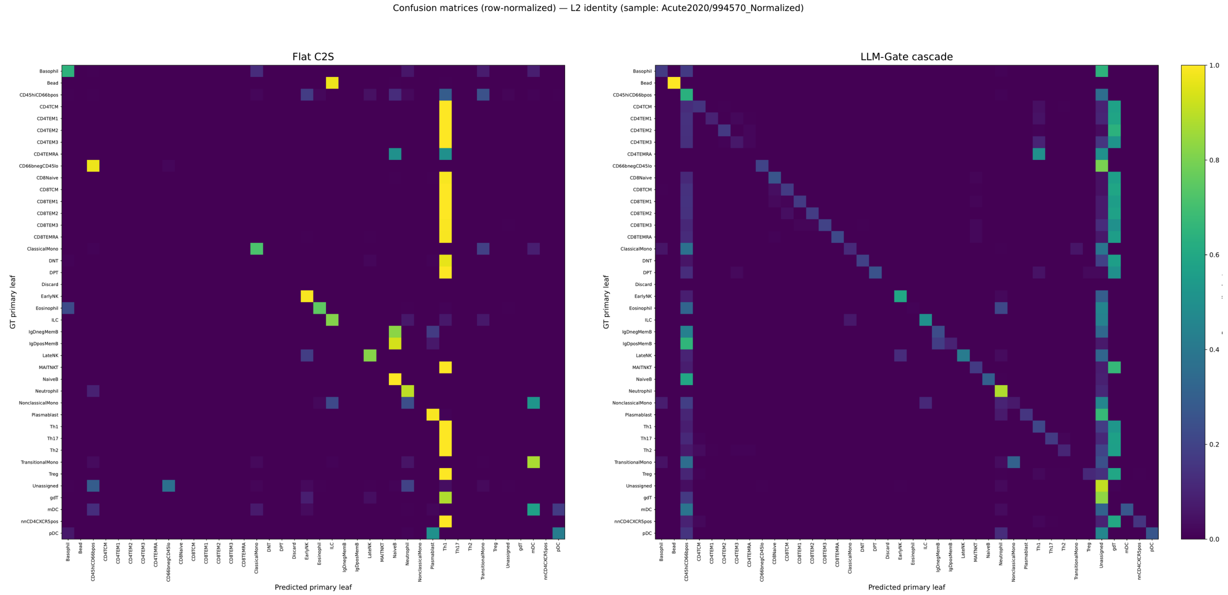


Figure S6. Row-normalized primary-leaf confusion at L2 (38 leaves), Acute2020/994570_Normalized, related to Figure 5.

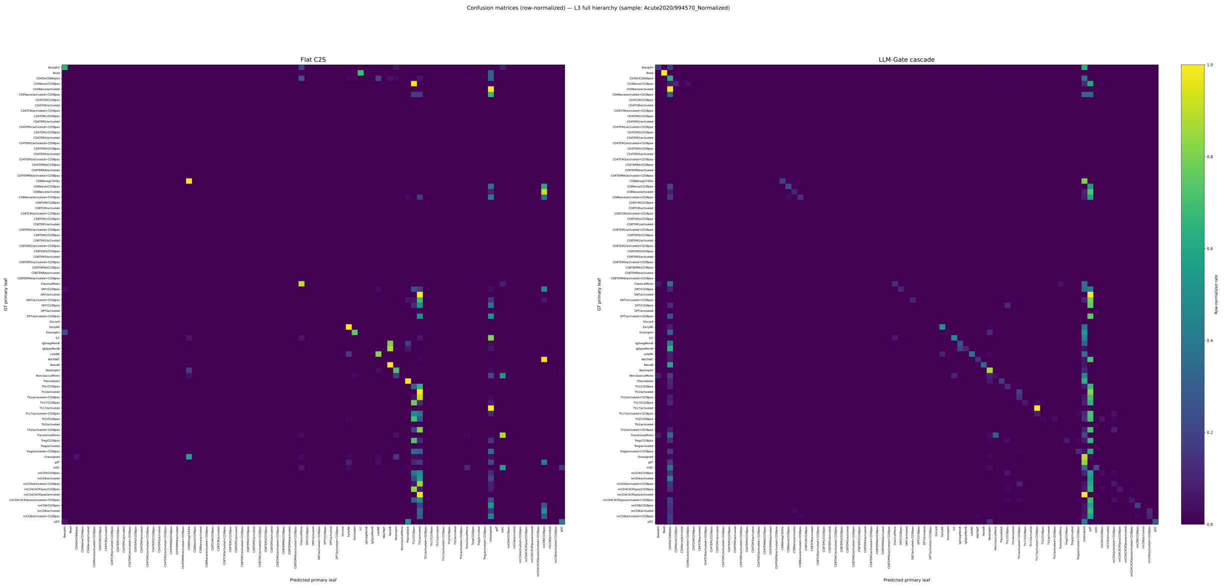


Figure S7. Row-normalized primary-leaf confusion at L3 (83 leaves), Acute2020/994570_Normalized, related to Figure 5. Off-diagonal column-stripes for both methods correspond to the cascade-rejected mass collapsed under *Discard* (or *Unassigned*, when a cell's predicted path terminates before reaching a leaf).

Acute2020 / 994570_Normalized — cascade (22 steps)

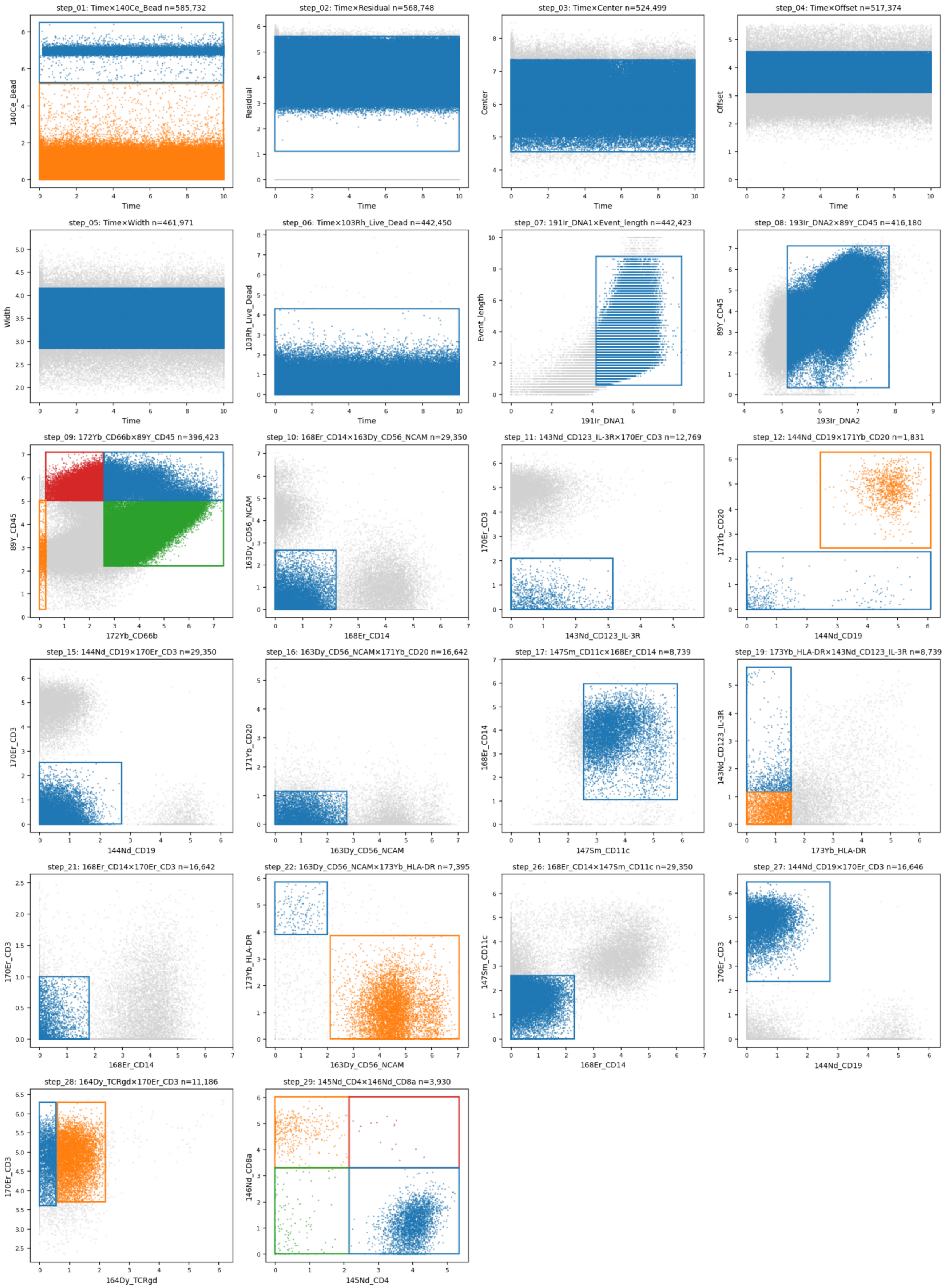

Figure S8. LLM-Gate cascade trace at L1 (22 steps), Acute2020/994570_Normalized: per-step biaxial scatter overlaid with the LLM-emitted rectangle gate that produced the next predicted-parent population, related to Figure 5. The trace illustrates how the cascade compounds: each panel's surviving cells are exactly the input to the panel below it.

Acute2020 / 994570_Normalized — cascade (41 steps)

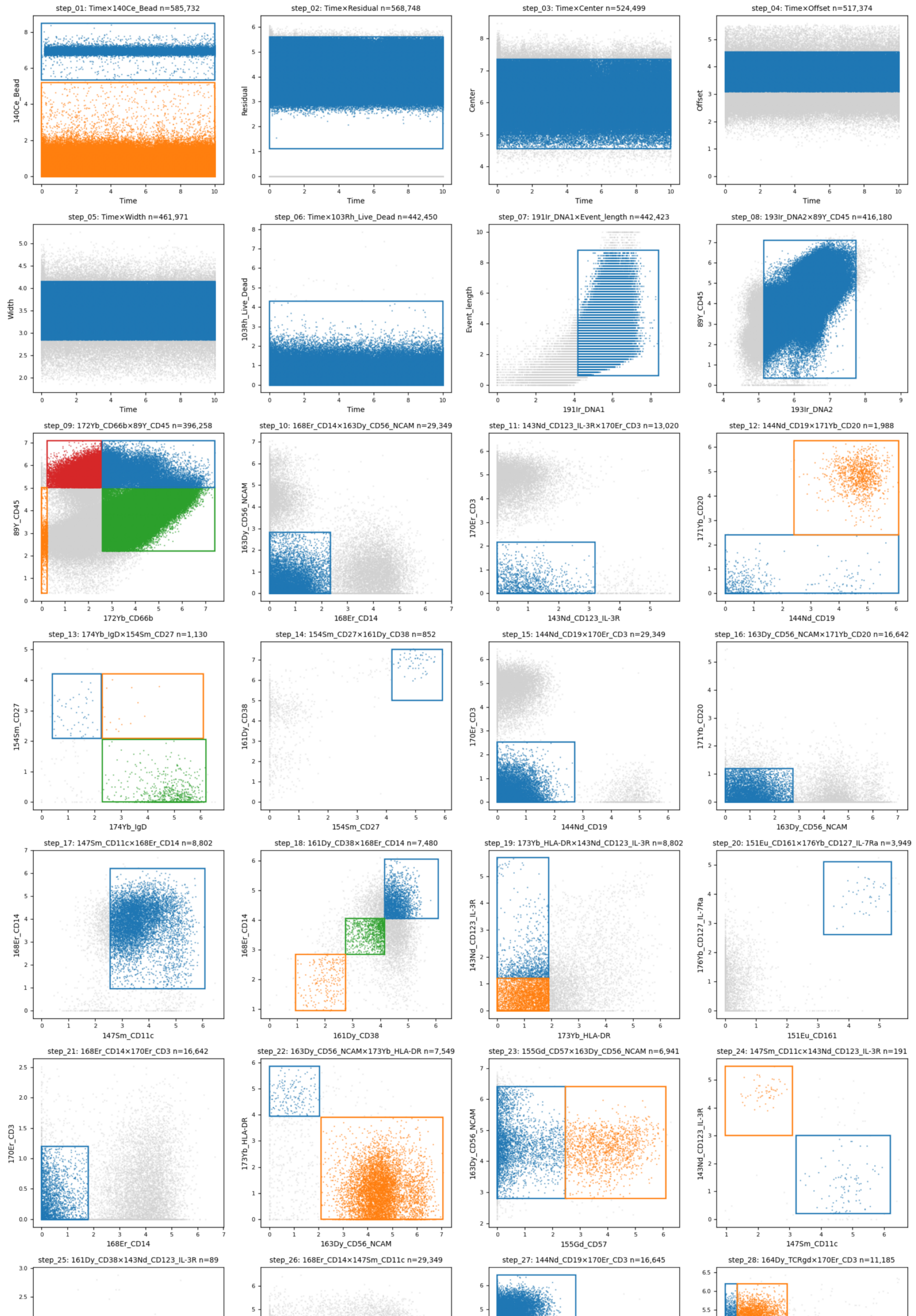

Figure S9. LLM-Gate cascade trace at L2 (41 steps), Acute2020/994570_Normalized, related to Figure 5.

Acute2020 / 994570_Normalized — cascade (62 steps)

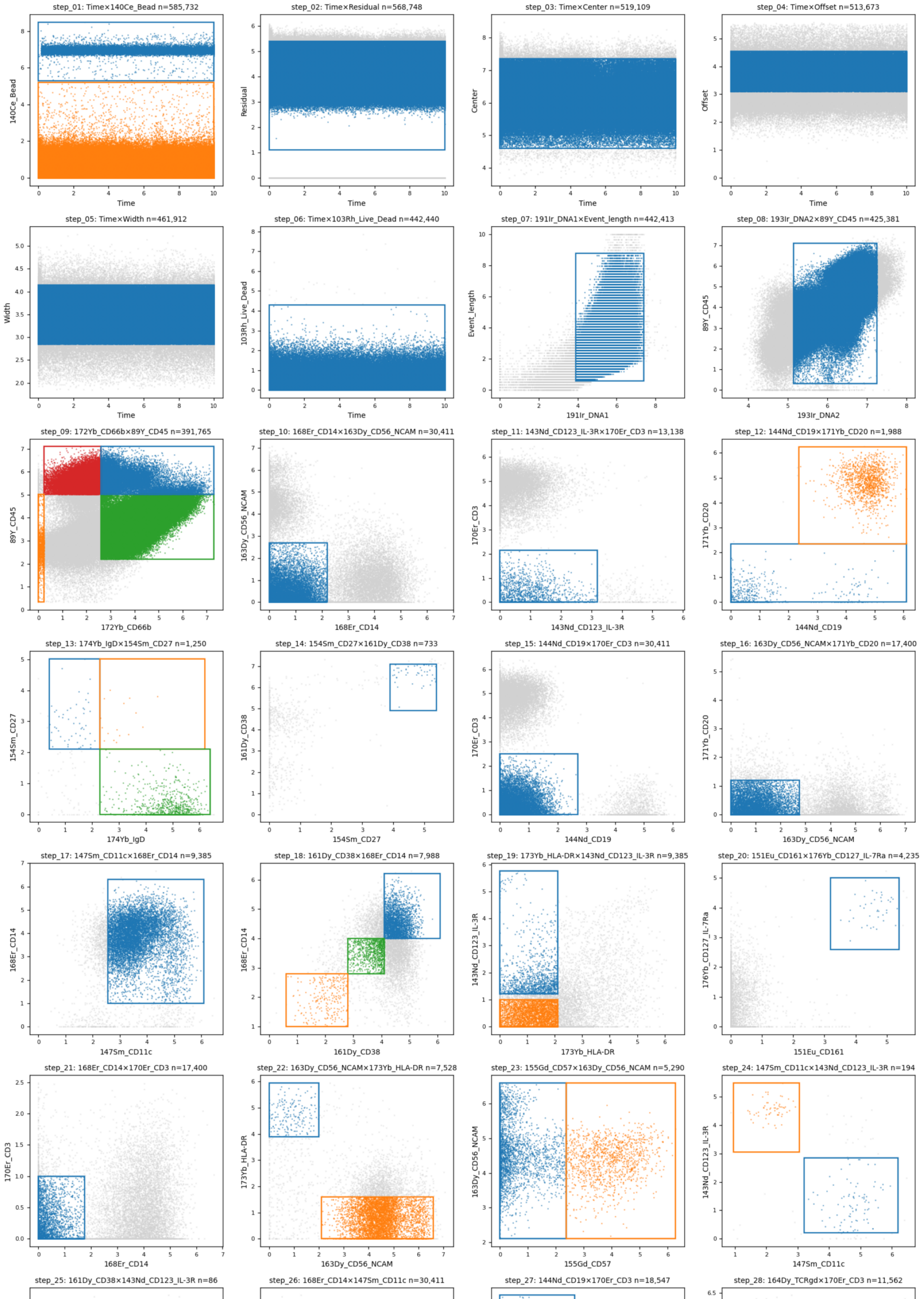

Figure S10. LLM-Gate cascade trace at L3 (62 steps), Acute2020/994570_Normalized, related to Figure 5.

### S10 Reproducibility statement

All experiments are run zero-shot with fixed generation settings. The closed-weight backbone is decoded greedily, and the open-weight reasoning backbones use model-card sampling (temperature 1.0, top-$p$ 0.95, top-$k$), whose seed-stability is reported in Table S5 (section S6.3). The method roster, backbones, and the donor-stratified 1:1:2 train/val/test split with seed 42 are specified in section S6.1 and section S2.1. Metric definitions (F1 macro, balanced accuracy, Hull IoU) and the cohort-level aggregation are given in section S6.2. The complete LLM-C2S and LLM-Gate prompts, the JSON output schema validated against, and the per-paradigm output handling are reproduced verbatim in section S4 and section S5. Per-cohort source, format, and harmonization exclusions are listed in section S2.2. Serving and hardware (vLLM on $8 \times$ H200) are reported in section S6.4. We release the evaluation code and the frozen per-step test split with the submission.

### REFERENCES


1. Van Gassen, S., Callebaut, B., Van Helden, M.J., Lambrecht, B.N., Demeester, P., Dhaene, T., and Saeys, Y. (2015). FlowSOM: Using self-organizing maps for visualization and interpretation of cytometry data. Cytometry Part A *87*, 636–645. https://doi.org/10.1002/cyto.a.22625.

2. Levine, J.H., Simonds, E.F., Bendall, S.C., Davis, K.L., Amir, E.D., Tadmor, M.D., Litvin, O., Fienberg, H.G., Jager, A., Zunder, E.R., et al. (2015). Data-driven phenotypic dissection of AML reveals progenitor-like cells that correlate with prognosis. Cell *162*, 184–197. https://doi.org/10.1016/j.cell.2015.05.047.

3. Li, H., Shaham, U., Stanton, K.P., Yao, Y., Montgomery, R.R., and Kluger, Y. (2017). Gating mass cytometry data by deep learning. Bioinformatics *33*, 3423–3430. https://doi.org/10.1093/bioinformatics/btx448.

4. Arvaniti, E., and Claassen, M. (2017). Sensitive detection of rare disease-associated cell subsets via representation learning. Nature Communications *8*, 14825. https://doi.org/10.1038/ncomms14825.

5. Kaushik, A., Dunham, D., He, Z., Manohar, M., Desai, M., Nadeau, K.C., and Andorf, S. (2021). CyAnno: A semi-automated approach for cell type annotation of mass cytometry datasets. Bioinformatics *37*, 4164–4171. https://doi.org/10.1093/bioinformatics/btab409.

6. Kim, J., Ionita, M., Lee, M., McKeague, M.L., Pattekar, A., Painter, M.M., Wagenaar, J., Truong, V., Norton, D.T., Mathew, D., et al. (2024). Cytometry masked autoencoder: An accurate and interpretable automated immunophenotyper. Cell Reports Medicine *5*, 101808. https://doi.org/10.1016/j.xcrm.2024.101808.

7. Ding, S., Bhattacharya, S., and Butte, A.J. (2025). ImmuneFM: Pre-training foundation model from cytometry data for immunology research. bioRxiv. https://doi.org/10.1101/2025.07.09.664020.

8. Malek, M., Taghiyar, M.J., Chong, L., Finak, G., Gottardo, R., and Brinkman, R.R. (2015). flowDensity: Reproducing manual gating of flow cytometry data by automated density-based cell population identification. Bioinformatics *31*, 606–607. https://doi.org/10.1093/bioinformatics/btu677.

9. Chen, J., Ionita, M., Feng, Y., Lu, Y., Orzechowski, P., Garai, S., Hassinger, K., Bao, J., Wen, J., Duong-Tran, D., et al. (2025). Automated cytometric gating with human-level performance using bivariate segmentation. Nature Communications *16*, 1576. https://doi.org/10.1038/s41467-025-56622-2.

10. Coppard, V., Szep, G., Georgieva, Z., Howlett, S.K., Jarvis, L.B., Rainbow, D.B., Suchanek, O., Needham, E.J., Mousa, H.S., Menon, D.K., et al. (2024). FlowAtlas: An interactive tool for high-dimensional immunophenotyping analysis bridging FlowJo with computational tools in Julia. Frontiers in Immunology *15*, 1425488. https://doi.org/10.3389/fimmu.2024.1425488.

11. Finak, G., Langweiler, M., Jaimes, M., Malek, M., Taghiyar, J., Korin, Y., Raddassi, K., Devine, L., Obermoser, G., Pekalski, M.L., et al. (2016). Standardizing flow cytometry immunophenotyping analysis from the human ImmunoPhenotyping consortium. Scientific Reports *6*, 20686. https://doi.org/10.1038/srep20686.

12. Montante, S., Yokosawa, D., and Brinkman, R. (2025). flowMagic gating benchmark: Automated and manual cell population annotation for flow cytometry data analysis. https://doi.org/10.20383/103.01352.

13. Bjornson-Hooper, Z.B., Fragiadakis, G.K., Spitzer, M.H., Chen, H., Madhireddy, D., Hu, K., Lundsten, K., McIlwain, D.R., and Nolan, G.P. (2022). A comprehensive atlas of immunological differences between humans, mice, and non-human primates. Frontiers in Immunology *13*, 867015. https://doi.org/10.3389/fimmu.2022.867015.

14. Fang, Y., Jin, Q., Xiong, G., Jin, B., Zhong, X., Ouyang, S., Yang, Y., Zhang, A., Han, J., and Lu, Z. (2026). Cell-o1: Training LLMs to solve single-cell reasoning puzzles with reinforcement learning. Bioinformatics, btag208.

15. Kwon, W., Li, Z., Zhuang, S., Sheng, Y., Zheng, L., Yu, C.H., Gonzalez, J.E., Zhang, H., and Stoica, I. (2023). Efficient memory management for large language model serving with PagedAttention. In Proceedings of the 29th symposium on operating systems principles (SOSP '23) (Association for Computing Machinery), pp. 611–626. https://doi.org/10.1145/3600006.3613165.

16. Rizvi, S.A., Levine, D., Patel, A., Zhang, S., Wang, E., Perry, C.J., Constante, N.M., He, S., Zhang, D., Tang, C., et al. (2025). Scaling large language models for next-generation single-cell analysis. bioRxiv. https://doi.org/10.1101/2025.04.14.648850.
